# The psychedelic DOI decreases the modularity of the cortical network by increasing cortico-cortical and thalamo-cortical functional influence

**DOI:** 10.64898/2026.07.20.739656

**Authors:** Roman Doronin, Eduard Stroukov, Georg B. Keller

**Affiliations:** Friedrich Miescher Institute for Biomedical Research, Basel, Switzerland; Faculty of Science, University of Basel, Basel, Switzerland

## Abstract

Psychedelics are powerful modulators of conscious perception, hypothesized to exert their effects by reorganizing large-scale brain functional connectivity. The circuit mechanisms underlying this reorganization remain unknown. Here, using widefield calcium imaging in awake mice, we found that the psychedelic DOI, in a 5-HT2A-dependent manner, acutely decreased the modularity of the cortical network by increasing correlations between anterolateral (AL) and posteromedial (PM) cortical domains. Functional influence measurements revealed that DOI increased both thalamo-cortical and long-range cortico-cortical influence onto the AL domain. We speculate that these increases in functional influence are driving the DOI-induced increase in the coupling between cortical domains. Our results provide a mouse correlate of the network reorganization seen under psychedelics in human neuroimaging and identify thalamo-cortical and cortico-cortical changes as candidate circuit mechanisms for this reorganization.

## INTRODUCTION

In humans, the psychedelic experience is accompanied by a dramatic functional disintegration of large-scale networks manifested as reduced functional connectivity within established brain networks, and enhanced connectivity between them (Beneš et al. 2025; Timmermann et al. 2023; Dai et al. 2023; Felix Müller et al. 2018; Preller et al. 2018; Palhano-Fontes et al. 2015; Lebedev et al. 2015; Gaddis et al. 2022; Carhart-Harris et al. 2013; Girn et al. 2026). Of these between-network changes, the most consistent pattern is an increase in functional connectivity between primary sensory areas and higher-level cognitive regions (Girn et al. 2026).

However, these findings offer limited access to the circuit mechanisms that are responsible for this effect. There are many different circuit mechanisms that could give rise to the observed changes in functional connectivity. For example, the increased between-network connectivity could result from an increase in cortico-cortical connections. Layer 5 (L5) neurons, the main contributor to long-range cortico-cortical projections, densely express 5-HT2A receptors (Andrade and Weber 2010), which are hypothesized to mediate psychedelic effects in humans (Vollenweider et al. 1998), and their cellular and firing properties are directly altered by psychedelics (Schmitz et al. 2022). Another possible circuit mechanism would be an alteration in the influence of subcortical inputs to cortex. Because a single subcortical source can project to multiple cortical areas, it provides a common input shared across them. Changes in the strength of such common input to cortex can result in changes to functional connectivity within cortex. One candidate for such modulatory input is higher-order thalamus. Higher-order thalamic connections span large parts of cortex and are well positioned to modulate large-scale cortical networks (Shine 2021). Moreover, thalamic neurons also express 5-HT2A receptors (López-Giménez et al. 2001; Barre et al. 2016) and in humans LSD administration induces widespread changes in thalamic resting-state functional connectivity with various cortical regions. In particular, thalamic functional connectivity with sensorimotor cortices is increased (Preller et al. 2018; F. Müller et al. 2017), but decreased with association cortices (Preller et al. 2018).

These cortico-cortical and thalamic candidates are not mutually exclusive, and human imaging offers limited opportunity for mechanistic circuit approaches to either. Doing so requires an animal model in which large-scale cortical activity can be measured under psychedelics and the implicated circuit elements can be interrogated. This approach is still hampered by a lack of clear understanding of whether psychedelics induce a rearrangement of the activity in large-scale cortical networks in the mouse, analogous to that observed in humans, and what these networks are. Here we leverage the advantages of the mouse model to investigate the circuit-level basis of acute psychedelic-induced network reorganization. We confirm that large-scale activity patterns in mouse cortex are dominated by a partition into an anterolateral (AL) and a posteromedial (PM) domain and show that the psychedelic DOI dissolves the functional separation between them in a 5-HT2A-dependent manner. Combining widefield calcium imaging with optogenetic functional influence mapping, we identify two candidate circuit-level substrates for this reorganization: an increase in thalamo-cortical drive to both the AL and PM domains, and an increase in cortico- cortical coupling between PM and AL. Together, these findings provide a mouse correlate of the human network reorganization and identify candidate circuit substrates for it.

## RESULTS

### Functional domains in mouse cortex

Functional networks have been previously characterized in mouse cortex (Vanni et al. 2017; Vafaii et al. 2024; Kochalka et al. 2026). To evaluate how DOI modulates these networks, we first sought to reproduce the reported baseline network architecture using widefield calcium imaging of dorsal cortex. We expressed a GCaMP variant using a retroorbital deposit of an AAV-PHP.eB vector to label neurons brain-wide (**Figure 1A**; **Methods**). For all imaging experiments, mice were head-fixed on a treadmill in a virtual reality setup and could freely locomote (**Figure 1B**) (Leinweber et al. 2014). Widefield data were downsampled using a grid of 818 regions of interest (ROIs) aligned to anatomical landmarks (**Figure 1A**). Based on calcium data recorded during quiet rest (**Methods**), we then first characterized the spatial structure of calcium activity in dorsal cortex using a correlation-based approach. For each of the 818 ROIs we calculated its correlation of calcium activity with that of all other ROIs. This gave us a seed correlation map for each of the 818 ROIs (**Figure 1C**). These maps were then thresholded (at 35% between minimum and maximum) to generate contour lines separating areas of high and low correlations. We then overlaid all of these contour lines. The result of this is what we refer to as a seed correlation contour map and visualizes cortical domains based on similarity of calcium activity (**Figure 1D**). We found that activity in dorsal cortex appeared to exhibit two domains, an anterolateral (AL) one and a posteromedial (PM) one. The AL domain covers lateral section of MOs and MOp as well as parts of somatosensory cortex, whereas the PM domain consists of medial MOs and MOp, sensory areas of cortex and retrosplenial cortex. To confirm the robustness of this domain structure, we used two additional approaches. First, we performed spectral clustering of ROIs, which operates directly on the pairwise activity correlation matrix. This approach treats the ROIs as nodes in a network and uses the eigenvectors of the connectivity matrix to project the data into a lower-dimensional space, allowing us to resolve complex, non-convex functional clusters. This recovered a similar AL-PM separation, with a two-cluster solution yielding the highest silhouette coefficient (**Figure 1E, S1A-B**). Second, we computed functional connectivity gradients, frequently used in fMRI analysis, that characterize the dominant axes of variation of functional connectivity (Margulies et al. 2016). Again, we found that the AL domain and the PM domain occupied different extremes of the first cortical gradient values (**Figure 1F**). This division into two domains is consistent with previous reports that cortical activity is segregated into functional modules (Vanni et al. 2017). The AL domain also emerges as a distinct cluster when parcellating calcium activity in dorsal cortex into different numbers of clusters (Vafaii et al. 2024). We therefore conclude that the dominant segregation of calcium activity in mouse dorsal cortex is into an AL and a PM domain.

**Figure 1.**
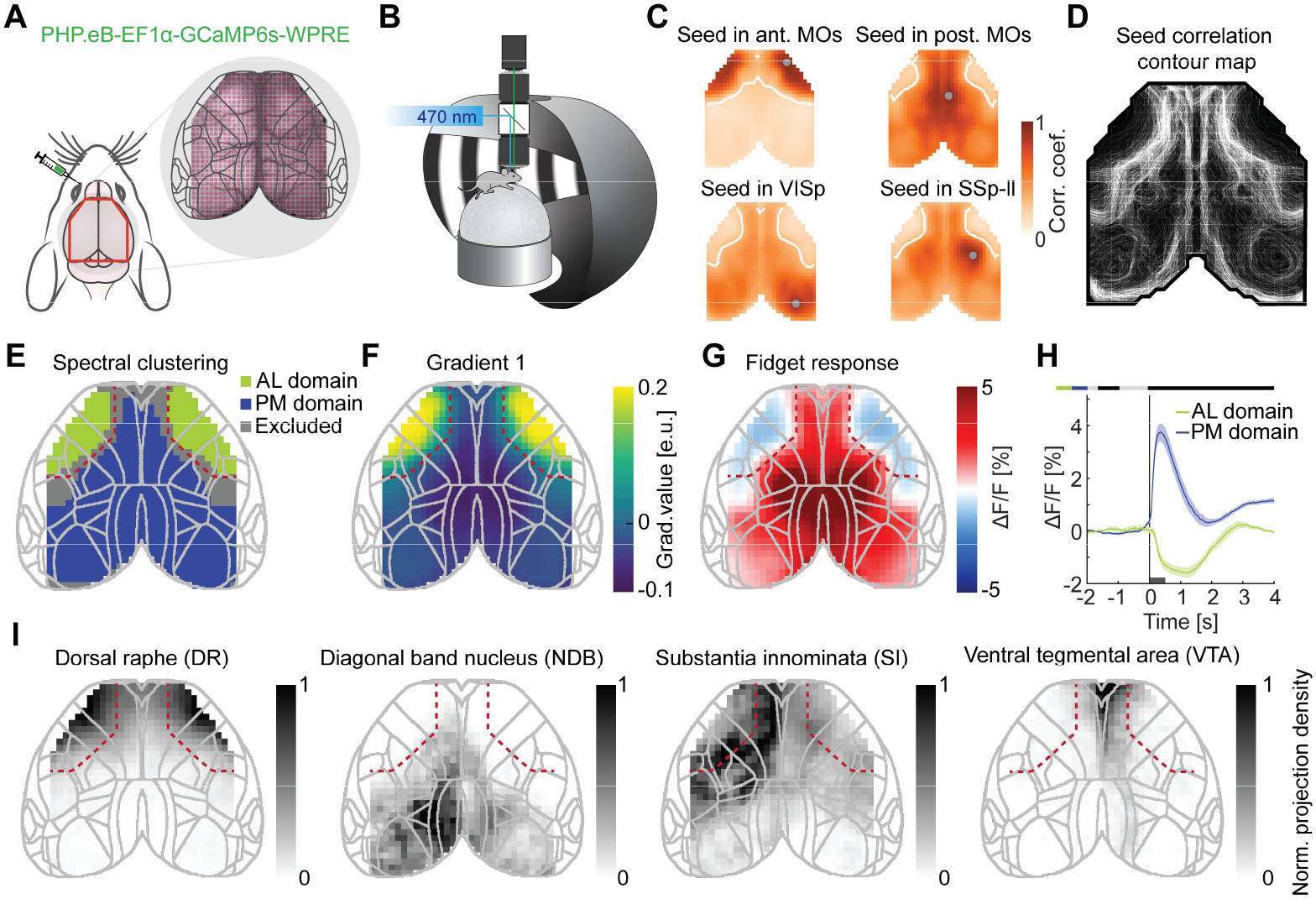
Calcium activity exhibits a two-domain organization in mouse cortex: anterolateral (AL) and posteromedial (PM). (**A**) Left: Schematic showing retroorbital AAV deposits used to express GCaMP6s brain-wide and the approximate extent of the cranial thin-skull window (marked in red) for the widefield recordings. Right: Example widefield image of the thin-skull cortical window registered to the grid of 818 ROIs and Allen Mouse Brain Common Coordinate Framework (CCFv3). (**B**) Schematic of the widefield setup. Mice were head-fixed on a treadmill and explored a virtual reality corridor. Calcium activity in dorsal cortex was recorded with a widefield fluorescence microscope (**Methods**). (**C**) Seed correlation maps for four example seeds in anterior and posterior secondary motor area (MOs), primary visual area (VISp), and primary somatosensory area, lower limb (SSp-ll). The grey dot on each map marks the respective seed. Correlation with the seed is indicated by the color, white lines are isocontour lines drawn at 35% of the distance between minimum and maximum correlation. (**D**) Seed correlation contour map obtained by overlaying isocontour lines of all 818 seed correlation maps. (**E**) Spectral clustering of the activity correlation matrix with 2 clusters reveals an anterolateral (AL) and a posteromedial (PM) domain. We excluded ROIs whose silhouette coefficient was below 0.5, indicating a low confidence in the cluster assignment. For all following analyses we used these regions as the AL and PM domains. Here and elsewhere, the red dashed line marks the boundary between AL and PM domains as defined by spectral clustering. (**F**) The first cortical gradient of the average correlation matrix aligns with the primary division into AL and PM domains. (**G**) Spatial map of fidget responses (averaged in the 0-0.5 s window from the fidget onset). (**H**) Activity traces of fidget responses for the AL and PM domains. Short dark gray bar marks the analysis window used in **G**. Here and elsewhere, the shading represents one standard deviation of the hierarchical bootstrap estimate of the mean. Horizontal bar above the plot indicates time bins in which the responses are statistically different from each other (gray: not significant, black: p < 0.05). Colored bars to the left indicate which data are compared. (**I**) Normalized axonal projection densities in the cortical volume of different neuromodulatory systems obtained from the Allen Mouse Brain Connectivity Atlas.

To test how much of this functional distinction can be explained by hemodynamic occlusion effects (Attwell et al. 2010; Ma et al. 2016; Waters 2020; Yogesh et al. 2025), we repeated these experiments in a cohort of mice that expressed EGFP. While EGFP fluorescence exhibited domains, these did not correspond to AL or PM and instead reflected transverse and sagittal sinuses (**Figure S1C-D**). Thus, within the frequency range used for our analyses (**Methods**), hemodynamic occlusion cannot account for the AL–PM partition observed in calcium activity.

Having defined AL and PM domains from ongoing activity, we next tested whether they are also functionally distinguishable by their responses to behavioral events. Interestingly, we found the clearest functional separation with behavioral fidgets, spontaneous brief motor movements that mice perform during quiet rest (Ramadan et al. 2022). During behavioral fidgets, the PM domain exhibited a strong positive response, while the AL domain decreased its activity (**Figure 1G-H**).

The differential engagement of AL and PM during rest and fidgets suggests that the two domains are differentially recruited across behavioral states. Since behavioral state in cortex is shaped in large part by ascending neuromodulatory input (Lee and Dan 2012; Shine 2019), we asked whether the anatomical projection patterns of major neuromodulatory systems map onto the AL-PM partition. For this, we used the Allen Mouse Brain Connectivity Atlas (Wang et al. 2020), from which we calculated the cortical projection density for some of the major neuromodulatory systems. We found that the serotonergic projections from dorsal raphe (DR) match the AL domain relatively well, while cholinergic projections from the Diagonal band nucleus (NDB) preferentially innervate the PM domain (**Figure 1I**). By contrast, cholinergic projections from substantia innominata (SI) and projections from the ventral tegmental area (VTA) do not show projection patterns reflective of the AL-PM domain structure.

### DOI-induced breakdown of domains

A hallmark of psychedelic action in humans is the breakdown of functional network boundaries - reduced connectivity within and increased connectivity between networks (Dai et al. 2023; Girn et al. 2026; Siegel et al. 2024). Given the overlap between the AL functional domain (**Figure 1E**) and the serotonergic innervation from the dorsal raphe (**Figure 1I**), we speculated that the clear separation we observe between the AL and PM domains, with high within-domain and low between-domain correlations, would provide an ideal model to test how serotonergic psychedelics change large-scale interactions between cortical domains in the mouse. To do this, we used the 5-HT2 receptor agonist DOI, which causes psychedelic experiences in humans (Shulgin and Shulgin 1991). To distinguish acute drug effects from habituation effects, we used a within-animal three-session experimental design (**Figure 2A**): a baseline session (Day 1), a DOI session (Day 3), and a recovery session (Day 5). In humans, the subjective effects of DOI typically last for 24 hours (Shulgin and Shulgin 1991). In mice, similarly, DOI concentration in the blood and forebrain returns to baseline 24 hours after administration and the primary behavioral readout, the head-twitch response (HTR), subsides before this (de la Fuente Revenga et al. 2019). Consistent with this, we found that DOI robustly induced head-twitch responses that returned to baseline levels at the recovery timepoint (**Figure 2B**). Thus, while it is important to consider that DOI may drive plasticity that alters cortical function long-term, the acute effect of the drug has likely subsided at our recovery time point.

**Figure 2.**
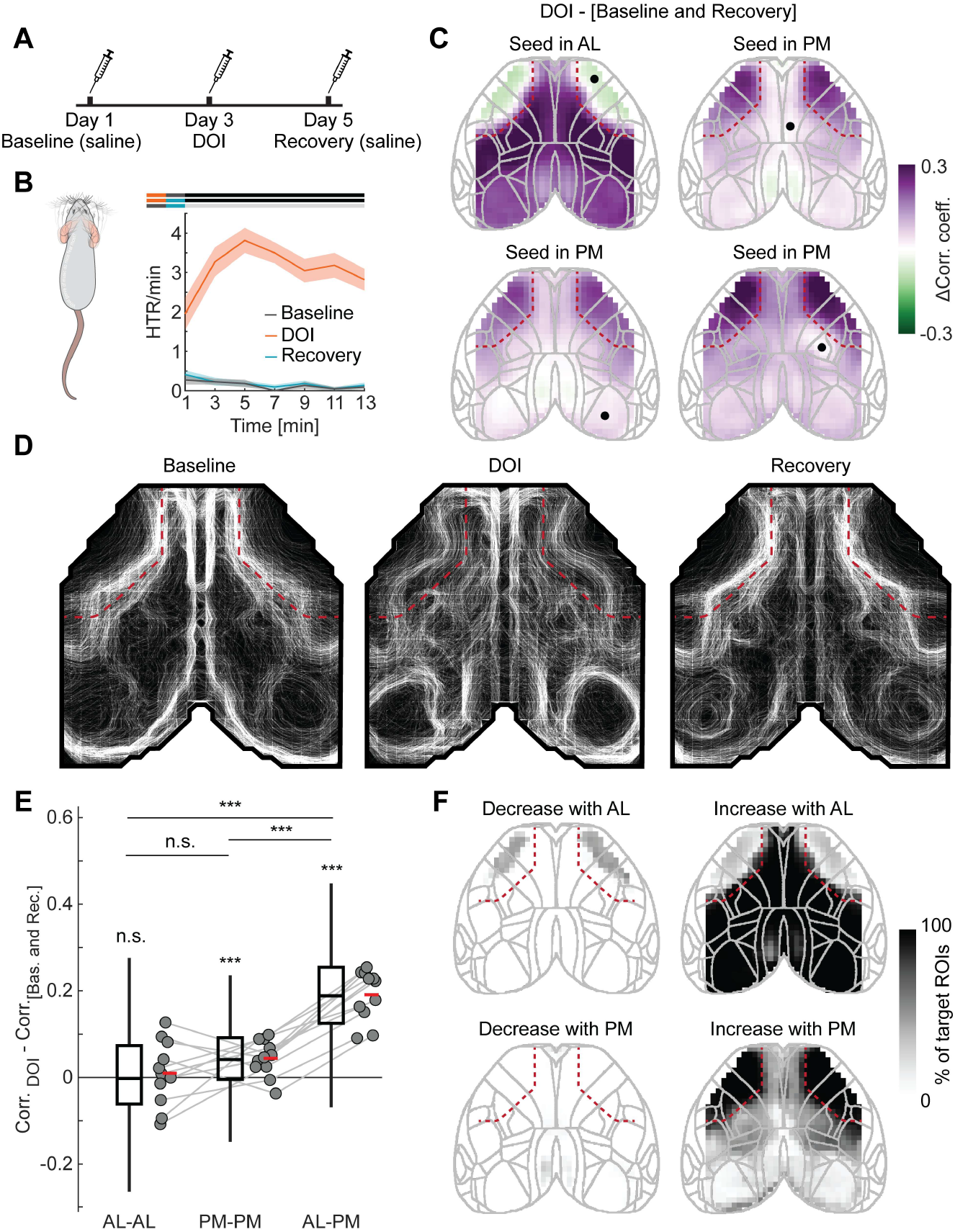
The psychedelic DOI dissolves the separation between the AL and PM domains. (**A**) Schematic of the three-session recording paradigm (days 1, 3, and 5). Each mouse was recorded following a baseline saline injection (2 days before DOI), a DOI injection, and a recovery saline injection (2 days after DOI). (**B**) Left: Schematic of the mouse head-twitch response (HTR). Right: Rate of head-twitch responses as a function of time since injection for baseline, DOI, and recovery conditions. (**C**) Difference in seed correlation maps for four example ROIs (black dots) between DOI and combined baseline and recovery conditions. (**D**) Seed correlation contour maps for baseline, DOI, and recovery conditions. (**E**) Change in correlation of activities between DOI and combined baseline and recovery conditions for pairs of ROIs within the AL domain, within the PM domain, and between the AL and PM domains. Boxplots show the distribution of differences in correlations for all pairs of ROIs for all mice. Here and elsewhere, whiskers extend to all data points not considered outliers (**Methods**), boxes mark quartiles and the central horizontal line the median of the distribution. Data points are averages per mouse and gray lines connect data from the same mice. Short horizontal red line indicates the mean over mice. Significance levels: n.s.: not significant; *: p<0.05; **: p<0.01; ***: p<0.001. (**F**) Spatial maps of DOI-induced changes in correlation with the AL domain (top) and the PM domain (bottom). For each ROI, shading indicates the percentage of target-domain ROIs (AL or PM) with which it exhibited a significant decrease (left) or increase (right) in correlation under DOI relative to combined baseline and recovery conditions. E.g. most ROIs in the PM domain exhibited an increase in correlation with close to 100% of the ROIs in the AL domain (Top right panel).

To probe for evidence of a DOI-induced change in the correlation structure of mouse cortical activity, we began by quantifying the effect of DOI on the seed correlation maps. For each seed, we computed the difference of the correlation map between DOI and no drug conditions (**Figure 2C**). For this and all following analyses, unless otherwise noted, we combined data from baseline (Day 1) and recovery (Day 5). The primary effects are unchanged if we just use one or the other (**Figure S2**). We found that for a seed in the AL domain, DOI markedly increased correlations with regions in PM, while resulting in modest decreases for within-AL correlations. Conversely, for seeds in the PM domain, correlations with the AL domain were strongly increased, while correlations within the domain exhibited more modest changes. The net effect of these changes was to reduce the separation between the AL and PM domains in the correlation structure. This can be visualized with the seed correlation contour maps (**Figure 2D**). The sharp boundary separating the AL and PM domains at baseline and recovery is largely dissolved under DOI. We quantified these effects in two ways. First, we calculated change in correlation for ROIs within AL, within PM, or between AL-PM per mouse (**Figure 2E**). We found no evidence of DOI-induced changes in correlations within the AL domain, a small increase within the PM domain, and a strong increase in the correlation of activity between domains. Second, for each ROI we visualized what fraction of AL or PM ROIs exhibited significant increases or decreases of correlation with that ROI (**Figure 2F**). This demonstrated that almost every ROI within a domain increased its correlation with every ROI of the opposite domain. Thus, DOI preferentially increased correlations of activity between the AL and PM domains.

Again, we tested whether the DOI-induced changes in activity correlation structure could be explained by influences of DOI on hemodynamic occlusion (Padawer-Curry et al. 2025) by repeating the experiments in mice expressing EGFP. Hemodynamic occlusion could influence our conclusions if it is differentially altered by DOI in the AL and PM domains, or if it acts as a common occluder, preferentially elevating correlations between weakly correlated regions. To address the first, we investigated whether DOI-induced vascular changes exhibited any domain specificity. We found no such effect (**Figure S3**). Second, we investigated whether an ideal common occluder that is uniform across dorsal cortex could differentially influence weakly and strongly correlated regions to produce the observed changes to AL-PM, AL-AL, and PM-PM correlations simultaneously. To do this, we selected an occluder that could explain either an increase in PM-PM or AL-PM changes and for each quantified how well the other two changes were described by this common occluder. We found no common occluder that could explain all DOI-induced changes in GCaMP activity simultaneously (**Figure S4**). Thus, neither region-specific nor uniform hemodynamic changes can account for the observed changes in correlation patterns.

We next addressed whether this effect of DOI is mediated by the 5-HT2A receptor, through which serotonergic psychedelics are thought to produce their acute effects in humans (Vollenweider et al. 1998). We reasoned that pretreatment with the 5-HT2A antagonist ketanserin (Leysen et al. 1981) should prevent the DOI-induced increase in AL–PM correlations if this effect is 5-HT2A-dependent. To test this, we repeated the experiments in a separate cohort of mice, administering ketanserin 30 minutes prior to DOI (**Figure 3A**). Consistent with previous results (Custodio et al. 2025), ketanserin prevented DOI-induced head-twitch responses (**Figure 3B**). Looking at seed correlation maps, we found that ketanserin plus DOI resulted in an overall reduction in correlations with no clear changes in domain structure (**Figure 3C-D**). Quantifying this we found that ketanserin not only prevented the DOI-induced increase in AL–PM correlations, but reversed the effect (**Figure 3E-F**): AL–PM, AL–AL, and PM–PM correlations all decreased below baseline, indicating a global decrease of cortical correlations. Thus, the DOI-induced increase in the correlation between the AL and PM domains is 5-HT2A dependent.

**Figure 3.**
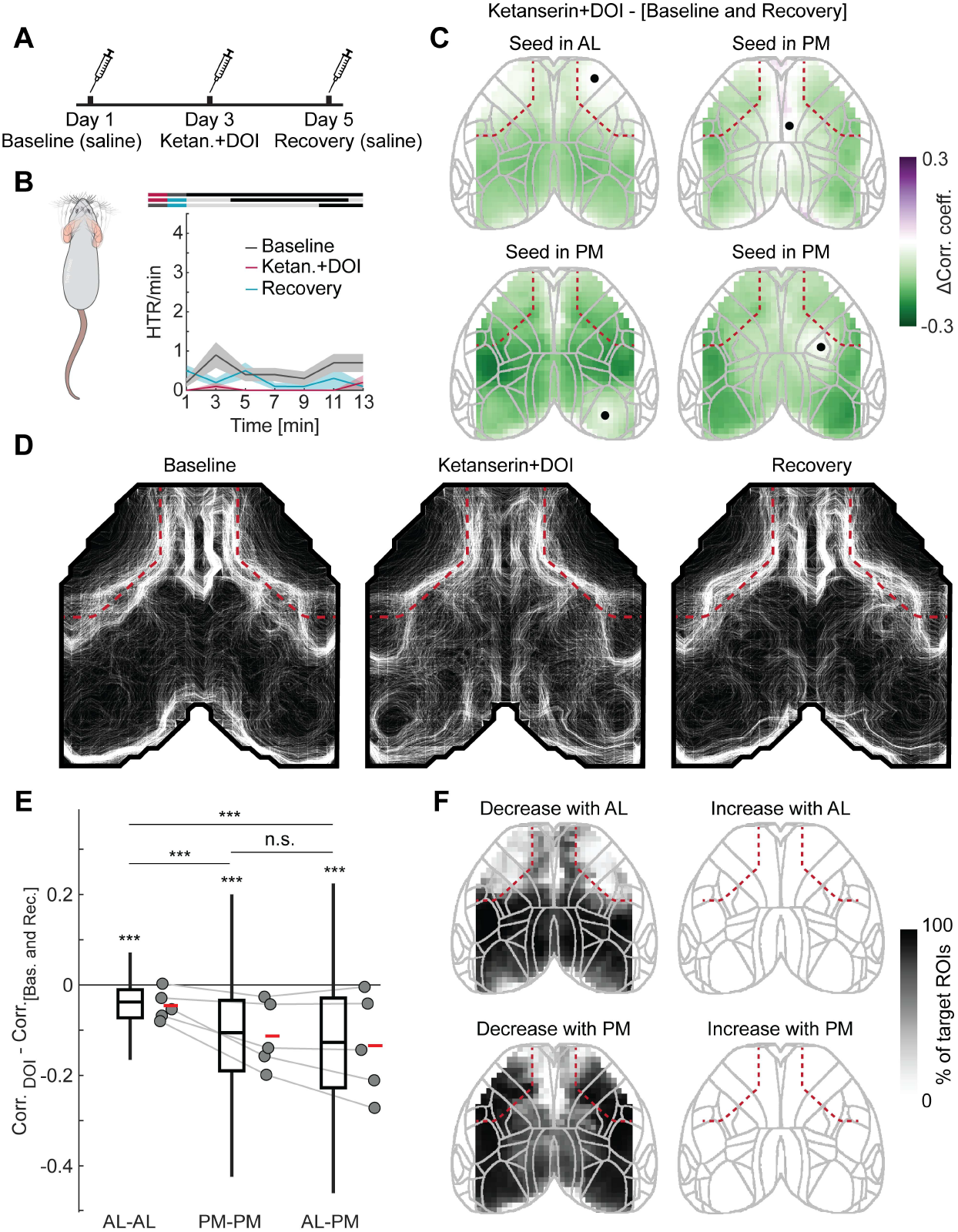
Ketanserin reverses DOI effects on correlations. (**A**) Schematic of the three-session recording paradigm. Each mouse was recorded following saline injection (2 days before ketanserin+DOI), after ketanserin pretreatment followed 30 min later by DOI injection, and a second saline injection (2 days after ketanserin+DOI). (**B**) Head-twitch response as the function of time since DOI injection for baseline, ketanserin+DOI, and recovery conditions. (**C**) Difference in seed correlation maps for four example ROIs (black dots) between ketanserin+DOI and combined baseline and recovery conditions. (**D**) Seed correlation contour maps for baseline, ketanserin+DOI and recovery conditions. (**E**) Change in correlation of activities between ketanserin+DOI and combined baseline and recovery conditions for pairs of ROIs within the AL domain, within the PM domain and between the AL and PM domains. Boxplots show the distribution of differences in correlations for all pairs of ROIs within or between domains for all mice. (**F**) Spatial maps of ketanserin+DOI-induced changes in correlation with the AL domain (top) and the PM domain (bottom). For each ROI, shading indicates the percentage of target-domain ROIs (AL or PM) with which it showed a significant decrease (left) or increase (right) in correlation under ketanserin+DOI relative to combined baseline and recovery conditions.

### Contribution of AL dynamics to the breakdown of domains

The DOI-driven increase in correlation between the AL and PM domains can be understood at two distinct levels. At a statistical level, the variation of activity of each domain can be decomposed into a domain-specific component and a common component. The correlation between domains increases if the common component of activity variation grows in relative weight, or if a domain-specific component decreases. At a circuit mechanism level, each of these statistical changes can be produced by one or more circuit alterations, for example a decrease in the domain-specific component could be caused by a decrease in an input specific to that domain or through domain-intrinsic network mechanisms. We therefore proceeded in two stages. We first used activity and correlation measurements (**Figure 4, 5**) to establish which statistical changes could underlie the increased AL-PM correlation, and subsequently used optogenetic influence mapping (**Figure 6, 7**) to probe for potential circuit mechanisms underlying these changes.

**Figure 4.**
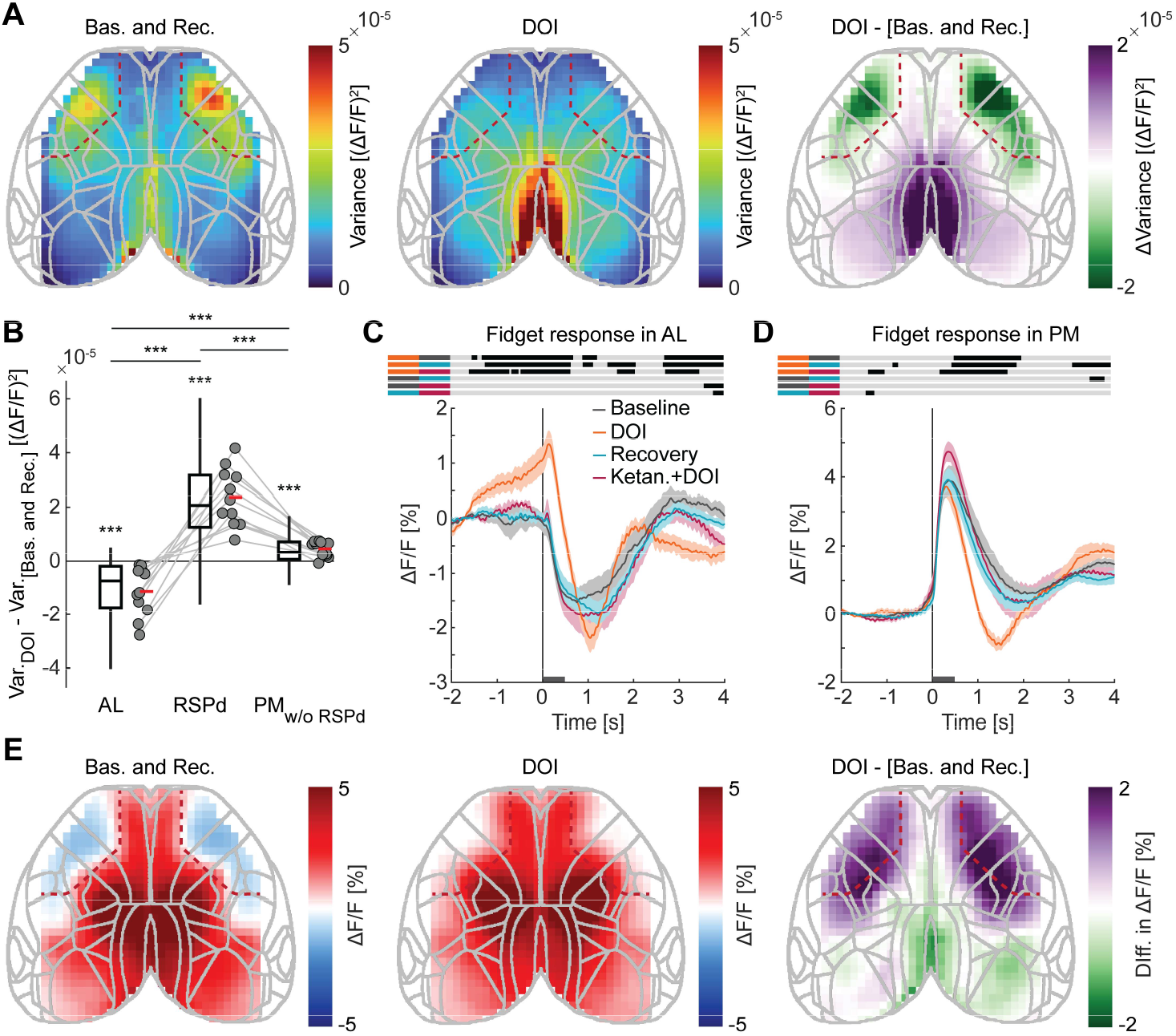
DOI differentially alters calcium activity in AL and PM domains. (**A**) Spatial maps of the variance of calcium activity for the combined baseline and recovery conditions, DOI, as well as the difference between the two. (**B**) Boxplots for the difference in variance between DOI and combined baseline and recovery for the ROIs in the AL domain, RSPd, and the PM domain without RSPd. (**C**) Activity traces of fidget responses in the AL domain for baseline, DOI, recovery, and ketanserin+DOI conditions. Short dark gray bar marks the analysis window used in **E**. Activity traces for baseline, DOI, and recovery conditions are presented for the dataset shown in Figure 1**,2**. Activity traces for the ketanserin+DOI condition are presented for the dataset shown in Figure 3. (**D**) As in **C**, but for the activity traces in the PM domain. (**E**) Spatial maps of the fidget responses for the combined baseline and recovery conditions, DOI, and the difference between the two.

**Figure 5.**
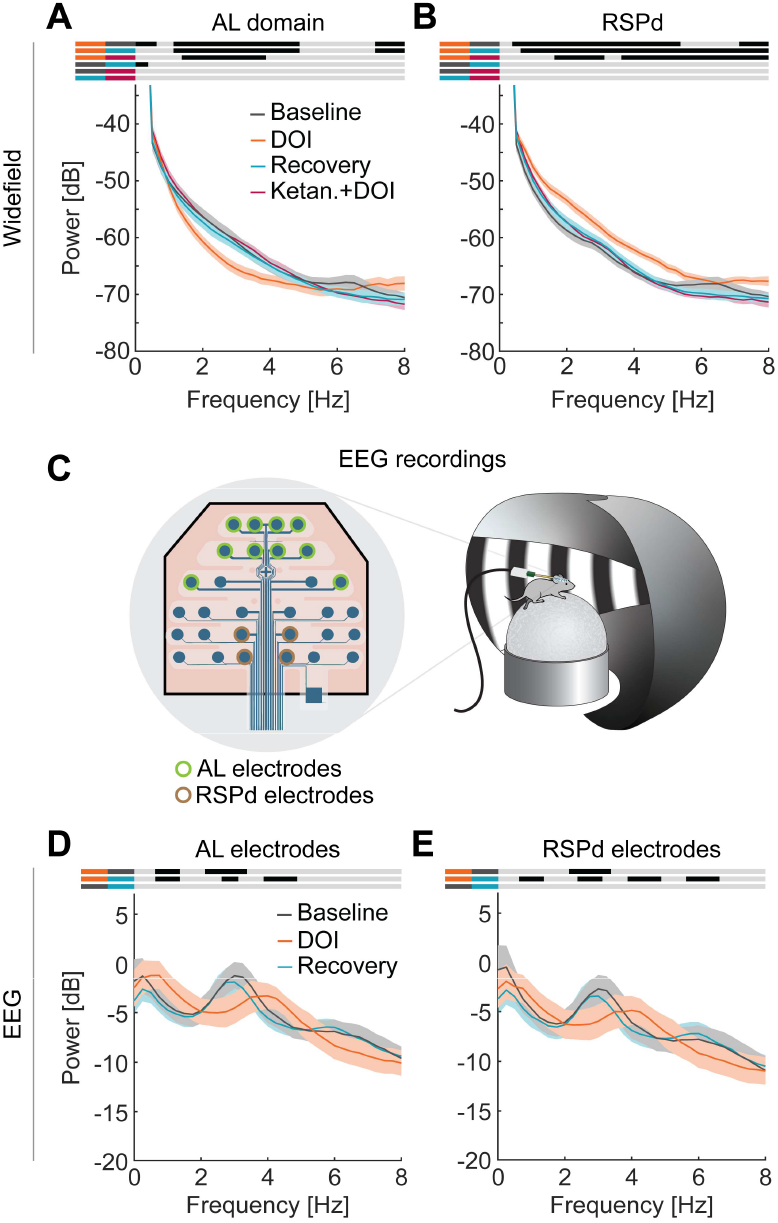
Decrease in AL domain variance of activity is concentrated in the 1-4 Hz band. (**A**) Power spectral density of the widefield calcium signal in the AL domain for baseline, DOI, recovery, and ketanserin+DOI conditions. Spectra for baseline, DOI, and recovery conditions are derived from the dataset shown in Figure 1**,2**. Spectra for the ketanserin+DOI condition are derived from the dataset shown in Figure 3. (**B**) As in **A**, but for RSPd. (**C**) Schematic of the EEG recordings. Left: Schematic of the EEG grid with the marked AL and RSPd electrodes used for the power spectra in **D, E**. Right: EEG grid was placed on top of the thinned skull while mice were head-fixed and allowed to run on the treadmill in the virtual reality corridor. (**D**) Power spectral density of the EEG signal for the AL electrodes in **C** for baseline, DOI, and recovery conditions. (**E**) As in **D**, but for the EEG signal for the RSPd electrodes.

**Figure 6.**
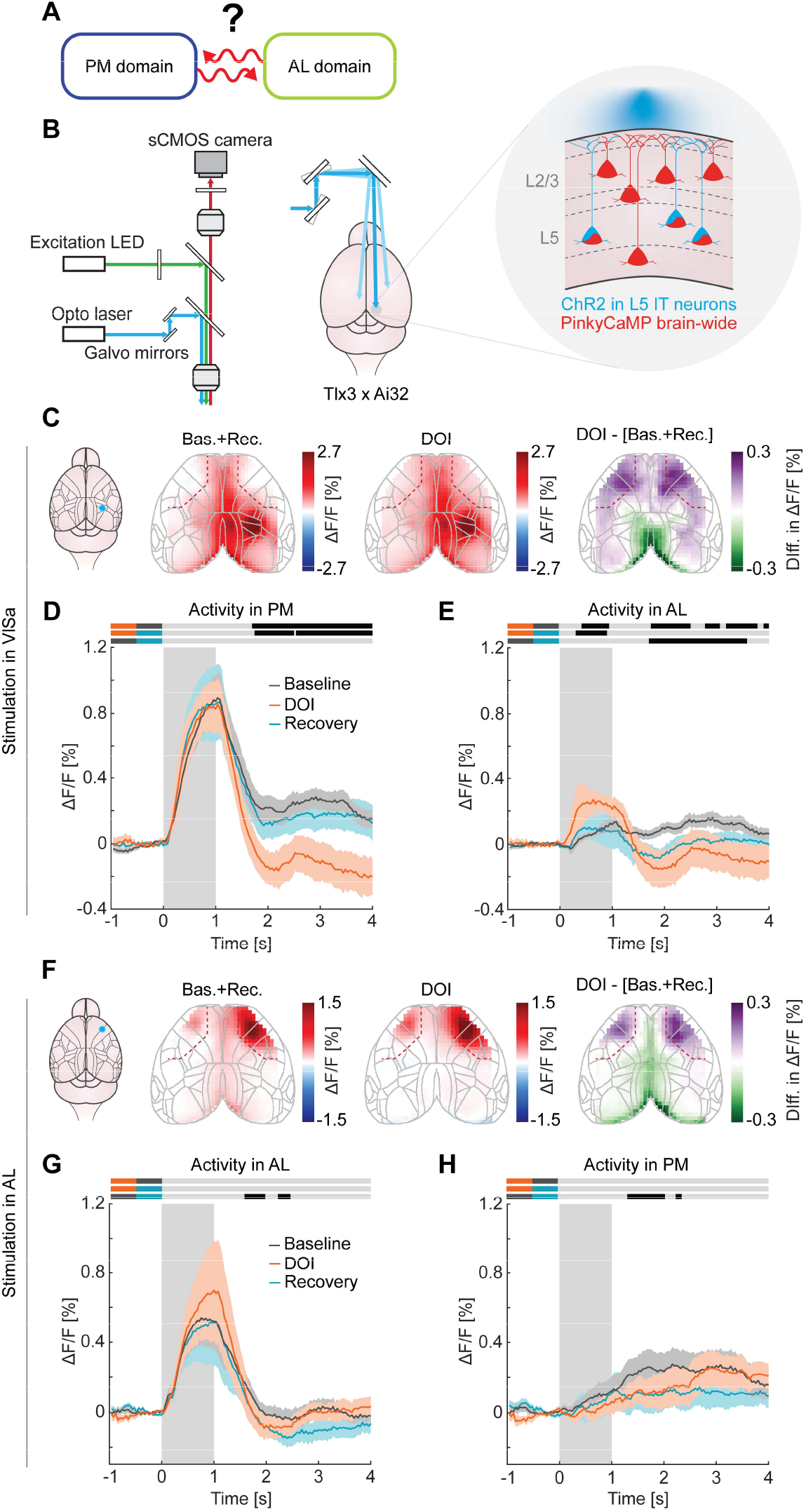
DOI increases cortico-cortical functional influence onto the AL domain. (**A**) Schematic of the hypothesis tested: Does DOI change cortico-cortical functional influence? (**B**) Schematic of the widefield optostimulation setup. A blue laser was targeted with galvo mirrors to different cortical locations to activate channelrhodopsin-2 (ChR2) while simultaneously imaging the activity of the red calcium indicator (PinkyCaMP) across dorsal cortex. ChR2 was expressed in L5 IT Tlx3 neurons by crossing Tlx3-Cre mice with the Ai32 reporter line. (**C**) Left: Schematic of the stimulation location in VISa. Right: Spatial maps of the response to VISa stimulation for the combined baseline and recovery condition, DOI condition, and the difference between the two, in the 0.5-1 s window. (**D**) Mean calcium response of the PM domain to the stimulation in VISa. Gray shading indicates stimulation interval. (**E**) As in **D**, but for the activity in AL. (**F**) As in **C**, but for the stimulation in AL. (**G**) As in **D**, but for the activity in AL, and for the stimulation in AL. (**H**) As in **D**, but for the activity in PM, and for the stimulation in AL.

**Figure 7.**
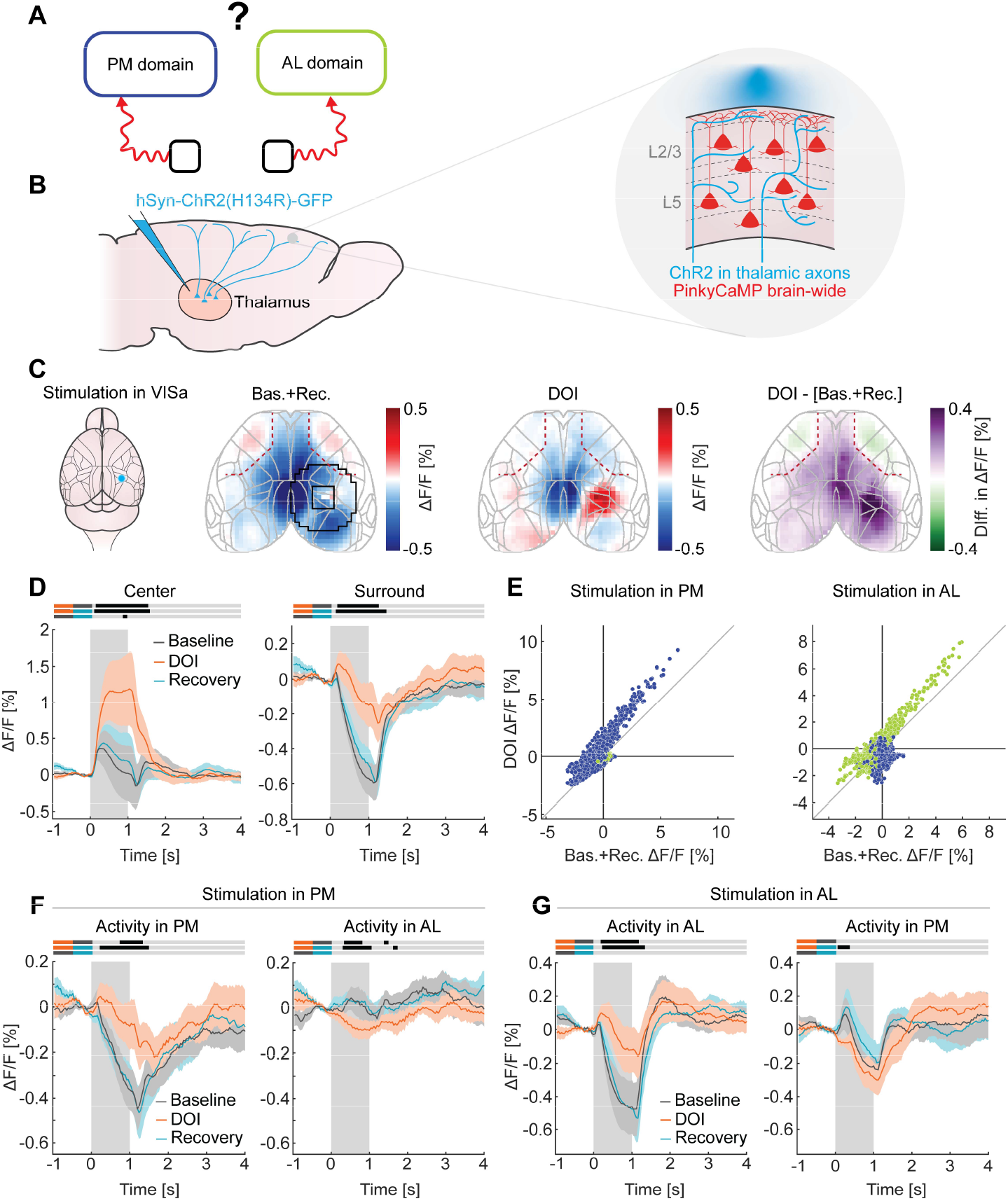
DOI increases thalamo-cortical functional influence in both AL and PM domains. (**A**) Schematic of the hypothesis tested: Does DOI change thalamo-cortical influence onto the AL and PM domains? (**B**) Left: Schematic of viral injection strategy. To express ChR2 in the thalamus, local AAV injections into the thalamus were used. Right: schematic of the expression and stimulation strategy. Thalamo-cortical axons expressing ChR2 were stimulated with a blue laser, while the activity of a red calcium indicator expressed brain-wide was recorded simultaneously. (**C**) Left: Schematic of the stimulation location in VISa. Right: spatial maps of the response to VISa stimulation for the combined baseline and recovery condition, DOI condition, and the difference between the two in a 0.5-1 s window. Black square contour around VISa and the surrounding circle indicate the ROIs used for calculating activity in the center and surround region around the stimulation sites in panel **D**. Note that the shape of the outlines (ideally both circles) are constrained by the ROI boundaries. (**D**) Left: Mean calcium response in the center set of ROIs closest to a stimulation site for the stimulation sites in VISa, RSP, and AL combined. Right: The same for the ROIs in the surround regions around a stimulation site. (**E**) Left: Scatter plot of the mean response of ROIs in 0.5-1 s window for the combined baseline and recovery conditions and DOI condition in response to VISa and RSPd stimulation. Points indicate ROIs of individual mice and are colored according to AL/PM identity. Right: The same but for the stimulation in an AL site. (**F**) Left: Mean calcium response in the PM domain for the combined stimulation sites in VISa and RSPd for baseline, DOI, and recovery conditions. Right: The same, but for the response in AL. (**G**) Left: As in **F**, but for the activity in the AL domain in response to stimulation in the AL domain. Right: The same, but for the response in PM.

An increase in the correlation between AL and PM domains could be the consequence of either an increase in the strength of a common component of activity variation or a decrease of an domain-specific component. These two statistical changes make different predictions for domain activity levels. If the increase in correlation were solely driven by an increase in a common component, we would - on first approximation - expect the amplitude of activity variation to increase in both domains. Conversely, in the case of a reduction of independent component - again on first approximation - one would expect a decrease in the amplitude of activity variation of one (or both) domains. To quantify the strength of changes in activity levels we calculated the variance of the calcium activity. This revealed a strong DOI-driven decrease in variance in the AL domain (**Figure 4A-B**). The PM domain showed only a modest increase in variance of activity overall, with the exception of retrosplenial cortex (RSPd) where we found a strong increase in variance (**Figure 4A-B**). We tested whether these changes could be explained by hemodynamic occlusion and found a DOI-driven reduction in variance, but this reduction was significantly weaker than for GCaMP data, was relatively homogeneous across cortex and did not reflect a domain-specific pattern (**Figure S5**). To test whether the DOI-driven reduction in AL variance could be the result of a general reduction in responsiveness, we investigated the effect of DOI on the fidget responses. Fidget responses were normally associated with a strong increase in activity in the PM domain, and a concurrent decrease in the AL domain (**Figure 1G-H**). Under DOI, fidget responses were altered primarily by the appearance of an increase in anticipatory activity in the AL domain (**Figure 4C, E**). Activity in the PM domain did not show anticipatory activity or a difference in peak activity but exhibited a faster decay of the positive response (**Figure 4D-E**). Pretreatment with ketanserin blocked both effects on the AL and PM domains (**Figure 4C- D**). Thus, we conclude that DOI results in a domain-specific change in the variance of calcium activity, decreasing variance in AL and increasing variance in RSPd. Of these two, only the decrease in AL variance is consistent with the decrease in the independent activity contributing to the observed decrease in functional separation of the AL and PM domains.

Finally, if the independent component of activity variation in the AL domain is indeed what creates separation between the domains in the cortical activity and its reduction is responsible for the increase of AL-PM correlations, we should find the following relationship: AL variance (normalized to PM variance) should be anticorrelated with the strength of AL-PM correlations on an individual mouse basis, and the effect of DOI should be in the same direction, shifting each mouse towards lower relative AL variance and higher AL-PM correlation. This is indeed what we found (**Figure S6A**). Taken together, these results strongly support the hypothesis that the DOI-induced increase in AL-PM correlation is primarily driven by a targeted reduction of independent activity variation within the AL domain.

### Relationship to psychedelic-driven EEG changes in humans

One of the most robust neurophysiological effects of psychedelics in humans is a general reduction in EEG power in the theta and alpha range, across all of cortex but most pronounced over posterior cortex (Riba et al. 2004; Muthukumaraswamy et al. 2013; Kometer et al. 2015; Carhart-Harris et al. 2016; Timmermann et al. 2023; Ip et al. 2026; Subramani et al. 2026), often accompanied by a shift of the peak alpha frequency to higher values (Carhart-Harris et al. 2016; Subramani et al. 2026). To investigate how the domain-specific effects we find may relate to these EEG changes reported in humans, we first computed power spectra for ROIs in AL and RSPd. We found that DOI reduced the spectral power in AL activity in a 1-4 Hz window, while it increased power in RSPd in that same window (**Figure 5A-B**). This pattern of change cannot be explained by hemodynamics (**Figure S7**) and was abolished by pre- treatment with ketanserin (**Figure 5A-B**). We suspect this change in AL power in the 1-4 Hz window is related to the change in AL-PM correlations as it is a similar frequency window that carries the correlation change (**Figure S6B-C**). We then repeated our experiments in a cohort of mice in which we conducted EEG recordings using a 30-channel electrode array (**Figure 5C**). We found that EEG power was reduced in the 2-3Hz range by DOI and the peak was shifted towards higher frequencies (**Figure 5D-E**). Both of these effects are consistent with EEG signatures of psychedelic action in humans (Carhart-Harris et al. 2016; Subramani et al. 2026). Interestingly, these effects were similar in recordings over the AL domain and in recordings over RSPd. Thus, the EEG recordings captured the reduction in calcium activity power in AL, but were blind to the increase in power in RSPd. Thus, the opposing effects we observe in calcium activity are difficult to relate to psychedelic induced EEG effects in humans.

### Circuit mechanisms

Based on the above arguments, our data would indicate that the DOI-driven increase in similarity of activity patterns between AL and PM domains is mediated by a domain-specific effect in AL. Thus, we turned to investigating possible circuit mechanisms that could underlie domain-specific changes to AL inputs. We focused on characterizing DOI-induced changes to the two main external inputs: long-range cortico-cortical projections (**Figure 6A**) and thalamo-cortical input (**Figure 7A**). To do this, we combined optogenetic stimulation with widefield calcium imaging to perform a series of functional influence mapping experiments. We expressed Channelrhodopsin-2 (ChR2) in neurons of a source area and PinkyCaMP, a red calcium indicator, brain-wide. We could then activate the processes of ChR2 positive neurons on the cortical surface using a focused blue laser and measure evoked responses in the target regions (**Figure 6B**). In the first experiment, we measured the DOI-induced change to long-range cortical input to the AL domain from the PM domain (**Figure 6A**). To do this, we focused on L5 IT neurons for two reasons: they are the dominant contributor to long- range cortico-cortical projections, and they strongly express 5-HT2A receptors (Shao et al. 2021), making them both the likely mediators of inter-domain communication and a direct target of psychedelic action. To stimulate these neurons selectively, we crossed the Tlx3-Cre line with the Ai32 reporter line to express ChR2 in L5 IT neurons cortex-wide (**Figure 6B**). PinkyCaMP was expressed via a retroorbital deposit in all neurons (**Methods**). Consistent with a preferential influence on AL, we found that DOI increased responses in AL to stimulation in VISa (**Figure 6C-E**). While we found a similar trend to increased responses when stimulating directly within AL, this effect was not significant (**Figure 6F-H**). Responses in PM during the stimulation period were not changed by DOI for either stimulation location. Thus, we conclude that DOI preferentially increases cortical influence in the AL domain.

To measure the functional influence of thalamic input to PM and AL (**Figure 7A**), we injected an AAV to express ChR2 into the left and right thalamus (**Figure 7B**). The left injection targeted regions of the thalamus that preferentially target AL in cortex (**Figure S8**). In the right thalamus we targeted one of two nuclei, both projecting to areas within the PM domain: in a subset of mice, we injected AAV into the AV nucleus, which projects to RSP, and in another subset of mice we targeted the LP nucleus, which projects to visual areas. Under baseline conditions, we found that stimulation of thalamic axons in the target cortical area tended to produce a slight focal increase of activity at the location of stimulation and produced an average decrease in the areas further away from the stimulation location (**Figure 7C-D, S9**). Under the influence of DOI, the focal excitation increased and the average suppression in surrounding areas was reduced (**Figure 7C- D**). This change of response seems to be additive: regardless of the baseline response, DOI increases the influence by approximately the same amount (**Figure 7E**). While the cortical response to the thalamic stimulation exhibits some domain specificity, the net effect of DOI was symmetric for both thalamic influence in AL and PM domains resulting in an additive increase in both (**Figure 7E-G**). Thus, we conclude that DOI increases thalamo-cortical influence on both AL and PM domains in additive manner.

Long-range cortico-cortical communication is predominantly mediated by intratelencephalic (IT) excitatory neurons in cortex that are also the primary target of thalamo-cortical input (Petreanu et al. 2009; Audette et al. 2018). If the primary effect of DOI on cortical activity patterns is mediated by changes to long-range cortico-cortical and thalamocortical communication, we might expect to find stronger effects of DOI in IT neurons. To test for possible cell-type-specific effects, we thus repeated the widefield imaging experiments using Cre lines to express GCaMP selectively in layer 2/3 and layer 4 excitatory neurons (Cux2-Cre), layer 5 IT neurons (Tlx3-Cre), layer 5 ET neurons (Fezf2-Cre), parvalbumin positive interneurons (PV-Cre), and somatostatin positive interneurons (SST-Cre). In all cell types, DOI induced a general increase in correlations (**Figure S10, S11**). However, only in the Cux2, Tlx3, and PV cell types, did it drive an AL-specific decrease in variance (**Figure 8**). As in the brain-wide recordings, this decrease in variance was primarily driven by a decrease in spectral power in the 1-4 Hz range (**Figure S12**), and the negative correlation between the strength of the relative AL variance and AL-PM correlations was only apparent in Tlx3, Cux2, and SST recordings (**Figure S13**). Thus, we conclude that the domain-specific effects of DOI are primarily carried by excitatory intratelencephalic neurons, but not by excitatory L5 ET/L6 neurons, with inhibitory interneurons exhibiting intermediate effects.

**Figure 8.**
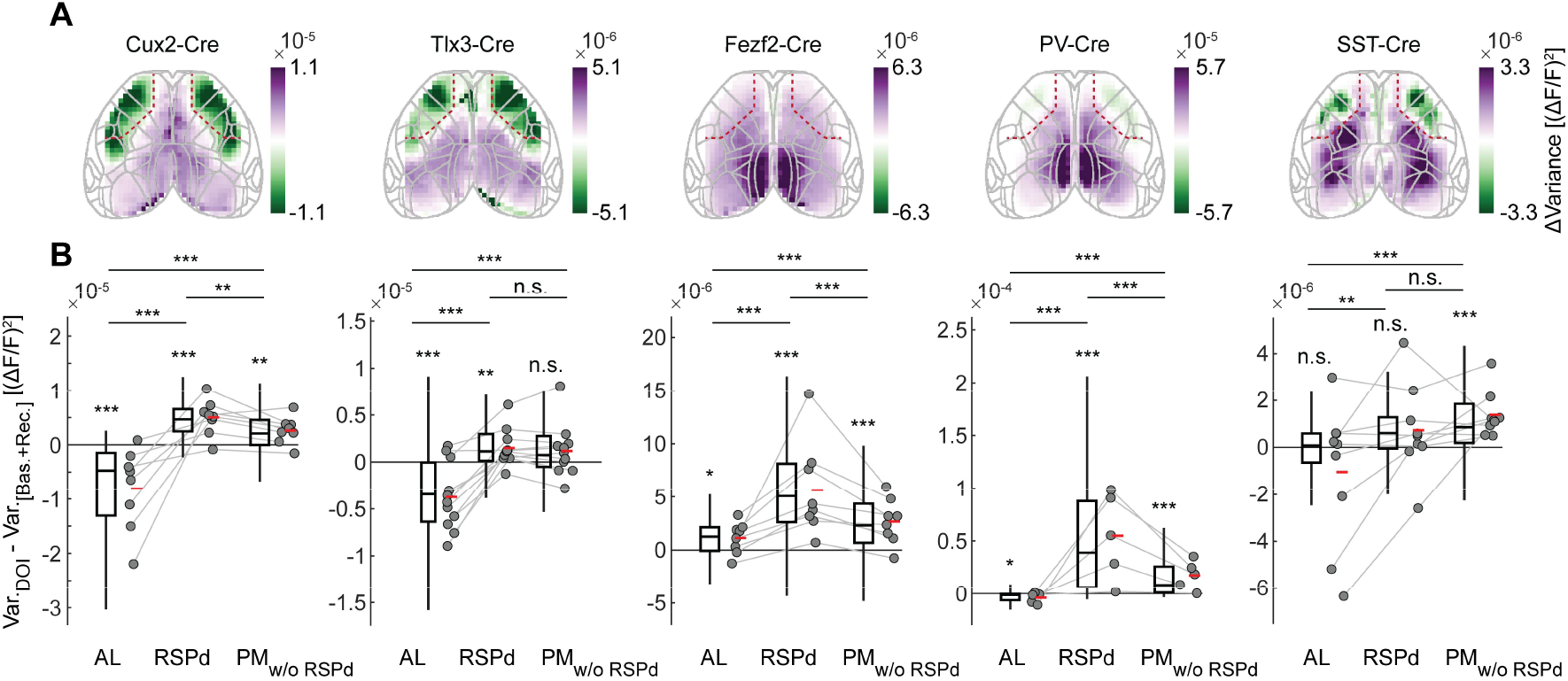
Cell-type-specific effects of DOI on AL and PM variance. (**A**) Difference in spatial maps of variance between DOI and combined baseline and recovery conditions for different cell types. (**B**) Boxplots for the difference in variance between DOI and combined baseline and recovery for the ROIs in the AL domain, RSPd, and the PM domain without RSPd for different cell types (**Methods**).

## DISCUSSION

In this work we show that the psychedelic DOI, via 5-HT2A agonism, acutely decreases the modularity of the cortical network by reducing the functional separation between the AL and PM cortical domains. In addition, we identify two candidate circuit mechanisms, a general increase in thalamo-cortical influence as well as a domain-specific increase of cortico-cortical influence onto the AL domain, that could plausibly account for this reorganization.

### Division of the cortex into AL and PM domains

The existence of anterolateral and posteromedial networks in mouse widefield imaging has been reported previously (Silasi et al. 2016; Wright et al. 2017; Vanni et al. 2017; Vafaii et al. 2024; Benisty et al. 2024). A similar partition into AL and PM domains is also visible in cortical projection patterns (Harris et al. 2019; Luo et al. 2019). What is peculiar is that the boundary between the AL and PM domains does not map neatly onto Allen atlas-defined brain areas – rather it runs right through the middle of primary motor and primary somatosensory cortex. What could the functional role of such a division be? The two domains receive differential neuromodulation (**Figure 1I**) (Nazari et al. 2023), with serotonergic projections from dorsal raphe specifically targeting the AL domain. The domains are also differentially engaged during goal-directed behavior: AL is preferentially active during anticipatory licking in a go/no-go task and drops after reward delivery despite continued licking (Allen et al. 2017). The sleep literature points to differential roles as well: during NREM sleep both AL and PM exhibit slow oscillatory activity, but during REM sleep only PM begins to desynchronize, possibly under cholinergic control (Nazari et al. 2023).

Functionally, the split into AL and PM domains is the dominant division of dorsal cortex. The PM domain is not itself a single uniform network, at finer parcellations it splits into multiple sub-networks (Vafaii et al. 2024; Kochalka et al. 2026), but the AL–PM separation emerges as the first and most robust axis of organization, appearing before these finer partitions (**Figure S1A**). The AL domain sits at the transmodal end of the first cortical gradient (**Figure 1F**) and during full-body fidgets, activity in the PM domain increases while it decreases in the AL domain (**Figure 1G-H**). In human fMRI imaging, these are the defining functional features of the division of cortex into a task-positive and a task-negative network – the task-negative network is also known as the default mode network (DMN) (Raichle et al. 2001; Margulies et al. 2016; Anticevic et al. 2012). In human cortex, the core consensus set of cortical areas that form the DMN is posterior cingulate, retrosplenial, and medial prefrontal cortex (Buckner et al. 2008). Thus, by anatomical criteria, AL is not an obvious DMN homolog, but by functional criteria, AL exhibits several of the defining DMN properties. Also indicative of such an analogy is the finding that there appears to be specific serotonergic influence in both the AL domain and the DMN: serotonergic fibers preferentially innervate AL (**Figure 1I**), while in humans 5-HT2A receptor density distribution positively correlates with DMN (Delli Pizzi et al. 2023). Based on this, we speculate that the AL domain is a functional analog of the DMN in humans. It is conceivable that the evolutionary origin of the DMN were parts of cortex responsible for self-directed movements.

One of the most consistent observations in the human psychedelic literature is increased functional connectivity between the DMN and sensory networks (Dai et al. 2023; Girn et al. 2026), with the changes in functional connectivity blocked by ketanserin (Preller et al. 2018). Because the AL domain shares functional features with the human DMN and the PM domain consists mostly of sensory cortices, the DOI-induced increase in AL–PM correlation we observe (**Figure 2, 3**) parallels effects reported in human cortex. Our results thus provide a candidate mouse correlate of this signature of psychedelics: a 5-HT2A- dependent increase in coupling between a transmodal, DMN-like domain and sensory cortex.

### EEG effects and changes in AL and RSPd variance

In mice, psychedelics are known to drive a decrease in low-frequency frontal EEG power (Olson et al. 2024; Golden and Chadderton 2022). The anatomical source of this frontal slow rhythm has remained unclear. In our widefield recordings (**Figure 4A, 5A, 8A**), the DOI-induced decrease of variance was largely restricted to the excitatory intratelencephalic neurons in the AL domain, with other cortical areas showing an increase. This regional and cell-type specificity raises the possibility that the excitatory intratelencephalic neurons in the AL domain are the cortical source of the frontal slow rhythm reported in these earlier studies (Olson et al. 2024; Golden and Chadderton 2022; Karalis and Sirota 2022). The increase of the RSPd widefield power we find is more puzzling, as we did not find a correspondence between the widefield and EEG changes in RSP. However, the increase in RSPd variance is consistent with recent findings of increased low-frequency LFP power under psilocybin (Jiang et al. 2026). Given that increases in RSPd variance are a signature of dissociative drugs (Vesuna et al. 2020), this may be a more general effect of hallucinogens.

### Thalamic stimulation effects

One prevalent theory of psychedelic action, the cortico-striato-thalamo-cortical loop model (CSTC), postulates that psychedelics modulate cortical and subcortical structures, resulting in aberrant thalamo-cortical coupling (Vollenweider and Geyer 2001; Doss et al. 2022). Prior support for this idea has largely come from human imaging studies that reported changes in thalamo-cortical resting-state functional connectivity under psychedelics (Preller et al. 2018; Gaddis et al. 2022; F. Müller et al. 2017). Additionally, slice physiology work has shown that psychedelics increase NMDA-receptor-mediated excitatory postsynaptic currents in thalamo-cortical synapses (Barre et al. 2016). What has remained untested is whether and how psychedelics alter the way the thalamus modulates large-scale cortical networks. Our optogenetic data provide direct evidence for such a change: DOI leads to an additive increase in thalamo-cortical influence in both the AL and PM domains.

A notable feature of the baseline thalamo-cortical influence we measured is that, beyond focal excitation, it is on average suppressive (**Figure 7C-D, F-G**). Although thalamo-cortical feedforward inhibition is well described at the single-cell level, it was less expected that activating associative thalamic projections would produce net suppression across large cortical territories. This net suppression might be a general feature of associative thalamic projections: activation of medial dorsal thalamus produces mild suppression of excitatory neurons in prelimbic cortex (Mukherjee et al. 2020; Lupori et al. 2026). The anatomical basis for this net suppression is speculated to be PV interneurons that also receive thalamic input but may respond faster. Additionally, L1 inhibitory interneurons are excited more strongly than pyramidal cells by matrix thalamo-cortical input (Cruikshank et al. 2012). Together, these features might explain the net reduction in activity distant from the stimulation site. DOI reduces this suppression. The mechanism behind this reduction is not yet clear, but the effect appears to be additive, and not multiplicative (**Figure 7E**).

How does this finding relate to the human imaging literature? Resting-state studies report that psychedelics increase thalamic functional connectivity with sensorimotor cortex while decreasing it with associative cortex (Preller et al. 2018). Our finding, an additive increase in thalamo-cortical influence, does not map cleanly onto this pattern. On first approximation, one would expect an increase in influence to result in an increase in correlation between thalamus and cortex in resting-state measures. Thus, our data would predict increase of thalamic functional connectivity with both sensory and associative cortices, different from the human data. This could reflect species differences, drug differences (DOI versus LSD), a divergence between resting-state correlation and direct functional influence measures, or a need to refine the mapping between mouse AL and human associative cortex.

Alterations of thalamo-cortical function may be a broader feature of hallucinogenic states and partially shared with psychosis. Microinjections of the dissociative MK-801, an NMDA receptor agonist, into the mediodorsal thalamus, but not into prefrontal cortex, reproduce the cortical signatures of a systemic injection, implicating thalamus as a primary locus of action for at least one non-serotonergic hallucinogen (Kiss et al. 2011). Moreover, patients with psychosis show altered thalamo-cortical functional connectivity with a pattern partially overlapping the psychedelic state: Increased connectivity with sensorimotor cortices and decreased connectivity with prefrontal regions (Anticevic and Halassa 2023; Keller and Sterzer 2024), consistent with the historical origin of the CSTC model in psychosis research - whether the underlying circuit alterations are shared remains an open question.

### Limitations

Our study has several limitations, key among these are: The first concerns causality. We show that DOI alters the correlation structure between the AL and PM domains, reduces AL variance, and modifies both cortico-cortical and thalamo-cortical functional influence. These cortico-cortical and thalamo-cortical changes are plausible circuit substrates for the observed correlation and activity changes, given the capacity of higher-order thalamus to reconfigure large-scale cortical networks (Shine 2021) and the capacity of L5 IT neurons to broadcast activity across the whole cortex via long-range projections. However, we do not demonstrate that they are the cause. Closing this gap will require selectively preventing DOI-induced changes to either cortico-cortical or thalamo-cortical influence directly – to the best of our knowledge, such an experiment, however, is not technically possible yet.

The second limitation concerns the relationship between the cortico-cortical and thalamo-cortical changes we report. These could be independent effects of DOI, or they could share a common origin. For example, if the primary action of DOI is on thalamo-cortical drive, the apparent change in cortico-cortical influence could arise indirectly through L5 IT neurons recruiting thalamus-projecting cortical neurons, which then engage the DOI-modulated thalamo-cortical pathway. The reverse confound is also possible: stimulating thalamic axons in cortex evokes cortical activity that propagates through cortico-cortical channels, so a change in evoked thalamo-cortical response could in part reflect altered cortico-cortical coupling. Disambiguating these effects would again require means to block the DOI effect selectively in one or the other pathway.

## Conclusion

Our findings point to two dimensions along which psychedelic action on large-scale cortical circuits can be described in the mouse: the AL–PM functional partition in cortex and thalamo-cortical influence. Together, these provide two circuit features that are sensitive to DOI and against which other interventions and conditions can be tested. Other non-serotonergic hallucinogens can now be compared by how they affect each domain: whether they dissolve the AL–PM separation as DOI does, and how they reshape thalamo-cortical influence onto each domain. This will test whether hallucinogens with different molecular targets converge on a shared cortical reorganization or act through distinct pathways. The same metrics can be applied to data from genetic risk models of schizophrenia, to ask whether their baseline cortical organization is already shifted toward the state DOI induces acutely. In this sense, our findings establish both a tractable mouse correlate of the large-scale network reorganization reported in human imaging and a circuit-level entry points for subsequent mechanistic work.

## ACKNOWLEDGMENTS

We thank all the members of the Keller lab for discussion and support, Tingjia Lu and FMI vector core for AAV production, and Jennifer Gröli for animal husbandry. Our work was supported by a team of core facilities at the FMI. We thank Olivia Masseck for sharing the PinkyCaMP plasmid prior to publication. We thank the Allen Institute for being awesome. The project has received funding from the Swiss National Science Foundation (GBK), the Novartis Research Foundation (GBK), and the European Research Council (ERC) under the European Union’s Horizon 2020 research and innovation programme (grant agreement No 865617) (GBK).

## DATA AVAILABILITY

Software for controlling the widefield microscopes and preprocessing of calcium imaging data is available at https://sourceforge.net/projects/iris-scanning/. Raw data and code to generate all figures will be deposited in public repositories upon publication.

## AUTHOR CONTRIBUTIONS

RD designed the study, performed widefield imaging and widefield stimulation experiments, and analyzed the data; RD and ES performed EEG experiments. GBK provided ice cream. All authors wrote the manuscript.

## METHODS

### Animals and surgery

All animal procedures were approved by and carried out in accordance with guidelines of the Veterinary Department of the Canton Basel-Stadt, Switzerland. For all surgical procedures, mice were anesthetized using a mix of fentanyl (0.05 mg/kg), medetomidine (0.5 mg/kg) and midazolam (5 mg/kg). Analgesics were applied perioperatively. Lidocaine was injected locally on the scalp (10 mg/kg subcutaneously) prior to surgery, while Metacam (5 mg/kg, subcutaneously), and buprenorphine (0.1 mg/kg subcutaneously) were injected just after completion of the surgery. In some experiments, we used Ethiqa XR (3.25 mg/kg subcutaneously) instead of buprenorphine and Metacam.

For widefield imaging experiments, we deposited an AAV-PHP.eB vector retroorbitally (5 μl per side) to express a fluorescent calcium indicator (GCaMP6s (Chen et al. 2013) or PinkyCaMP (Fink et al. 2026)) or EGFP, at least 1 week prior to the surgery. For cell-type-specific experiments to restrict the expression of a calcium indicator in different neuronal types, we used Cux2-CreERT2 (Franco et al. 2012), Tlx3-Cre (Gerfen et al. 2013), Fezf2-CreERT2 (Matho et al. 2021), PV-Cre (Hippenmeyer et al. 2005) and SST-Cre (Taniguchi et al. 2011) transgenic mouse lines. In a subset of mice, GCaMP6f was instead expressed using the Ai148 reporter line (Daigle et al. 2018). In the Cux2-CreERT2 and Fezf2-CreERT2 reporter lines, we administered tamoxifen food for at least two weeks to induce the activation of Cre. For all widefield imaging experiments we used thin-skull preparations. The skull over dorsal cortex was thinned with a dental drill to a thickness of approximately 50 µm. A custom-made acrylic window, thermoformed to match the curvature of the mouse skull, was glued to the skull with optical glue (Norland Optical Adhesive NOA 61). Finally, a custom titanium headbar was fixed to the skull using dental cement (Paladur, Heraeus Kulzer).

For thalamic stimulation experiments (**Figure 7, S9**) we injected an AAV to drive the expression of ChR2 (Boyden et al. 2005) into the left ventral antero-lateral complex of thalamus (VAL) (1.34 mm posterior to bregma, 1 mm lateral of bregma, and 3.25 mm below the cortical surface), right lateral posterior nucleus of thalamus (LP) (2.1 mm posterior to bregma, 1 mm lateral of bregma, and 2.5 mm below the cortical surface) or right anteroventral nucleus of thalamus (AV) (0.94 mm posterior to bregma, 0.9 mm lateral of bregma, and 2.6 mm below the cortical surface). To minimize backflow, the injection pipette was only retracted at least 5 min after the injection. For L5 IT neurons stimulation experiments (**Figure 6**), to restrict the expression of ChR2 in L5 IT neurons we crossed Tlx3-Cre transgenic line (Gerfen et al. 2013) with Ai32 line (Madisen et al. 2012).

For EEG experiments, the thinned skull was covered with a silicone casting compound (Kwik-Sil, WPI) before and between recording sessions.

### Drugs

2,5-dimethoxy-4-iodoamphetamine (DOI) hydrochloride (Sigma-Aldrich) was dissolved in 0.9% saline and administered intraperitoneally at dosages between 2.5 mg/kg and 5 mg/kg. Imaging experiments typically started 25 - 30 min after the injection. On baseline (Day 1) and recovery (Day 5) sessions, mice received an intraperitoneal injection of an equivalent volume of 0.9% saline. A subset of mice was used in multiple experimental series and received DOI multiple times spaced by at least 14 days. For ketanserin experiments, ketanserin was dissolved in 0.9% saline and administered intraperitoneally at 2.5 mg/kg 30 min prior to the DOI injection.

### Virtual reality setup

For all experiments, mice were head-fixed in a virtual reality setup (Leinweber et al. 2014) and free to locomote either on a spherical, air-supported treadmill or a wheel. A visual scene was projected (Samsung SP-F10M) onto a toroidal screen positioned in front of the mouse covering a field of view of approximately 160 degrees horizontally and 100 degrees vertically. The virtual reality setup was configured for one of the following paradigms: Closed loop, in which the visual stimulus consisted of a virtual corridor containing sinusoidal vertical gratings on the walls and a uniform gray floor. In this paradigm, movement in the virtual corridor was coupled to the locomotion speed of the mouse. Open loop, in which a replay of a previous closed loop session was projected. Dark, in which the projector was turned off., although residual dim ambient lighting was still present. In addition to these paradigms, we also presented full field drifting sinusoidal gratings (8 directions in randomized order, approximately 0.07 cycles/deg spatial frequency, 2 Hz temporal frequency). Each grating was presented for a randomized duration between 3 to 5 s with a randomized inter-trial interval of between 5 and 9 s during which a uniform gray screen was shown. All analyses except grating responses were performed for the data pooled for all types of recording sessions.

### Widefield imaging

Widefield imaging experiments were performed using a custom-built macroscope consisting of two objectives mounted face-to-face (Nikon 85 mm f/1.8 sample side, Nikon 50 mm f/1.4 sensor side). We used a 470 nm LED (M470L3 or M470L5, Thorlabs), powered by a custom-built LED driver for exciting GCaMP. Excitation light was passed through an excitation filter (FB470-10 or FBH470-10, Thorlabs) and combined with the emission pathway using a dichroic mirror (DMLP490L, Thorlabs). Green fluorescence was collected through a 525/50 nm emission filter (MF530-43, Thorlabs) on an sCMOS camera (PCO edge 4.2, Excelitas). We used collimator (SM2F32-A, Thorlabs) set to maximize the uniformity of the LED illumination across the surface of the cranial window. The resulting profile of the illumination cone was further trimmed with black tape on the sample side objective to avoid directly illuminating regions outside of the craniotomy. Raw images were acquired at 100 Hz using full dynamic range (16 bit) of the sensor with a resolution of 1107 by 1200 pixels, or 700 by 747 pixels, with an effective pixel size of either 10 by 10 µm or 16 by 16 µm. Raw images were cropped on-sensor and the resulting data were streamed to disk with custom software written in LabVIEW.

### Optogenetic stimulation and simultaneous widefield imaging

Simultaneous stimulation and imaging were performed on a custom-built macroscope consisting of two objectives mounted face-to-face (Samyang 85 mm f/1.8 sample side, Nikon 50 mm f/1.4 sensor side). PinkyCaMP (Fink et al. 2026) was excited with a 530 nm LED (M530L4, Thorlabs) powered by a custom-built LED driver. Excitation light was passed through a bandpass filter (FLH532-10, Thorlabs) and combined with the emission pathway using a dichroic mirror (DMLP567L, Thorlabs). Red fluorescence was collected through an emission filter (MF630-69, Thorlabs) on an sCMOS camera (PCO edge 4.2, Excelitas). Images were acquired at 25 Hz. For optogenetic stimulation, a 473 nm laser (OBIS LX 75 mW, Coherent) was directed via a pair of galvo mirrors (GVS002, Thorlabs) and a dichroic mirror (DMLP505L, Thorlabs) through the sample-side objective onto the cortical surface. Galvo mirrors were controlled using custom software written in LabVIEW, enabling targeting of the laser spot to arbitrary cortical locations within the imaging field of view. To avoid visual artifacts from the laser turning on and off, the laser was kept on continuously and parked on the headbar between trials, then steered to the cortical target only during stimulation. The laser spot diameter at the cortical surface was approximately 250 µm (full width at half maximum). To avoid contamination of the field of view by the reflected laser light, the laser was pulsed between the camera exposure times at 25 Hz with an 18% duty cycle. For cortical stimulation we used a laser irradiance between 3 and 6 mW/mm^2^. For stimulation of thalamic axons, laser irradiance was 30 mW/mm^2^. Stimulation duration was 1 s, with an inter-trial interval randomly sampled between 4 and 11 s. During the stimulation experiments mice were presented with a uniform gray screen.

### EEG recordings

For the duration of recordings, the silicone casting compound was removed and a 30-channel EEG electrode array (mouse-30-A-10-H32, NeuroNexus) was placed on top of the thinned skull and covered with a mix of saline and ultrasound gel (3:2 proportion). A ground wire was placed on the cement outside of the craniotomy and the reference wire was placed lateral to the EEG array. Signals were amplified using a 32 channel headstage (RHD 32ch, Intan Technologies) and digitized at 30 kHz using an Open Ephys acquisition board (Siegle et al. 2017). For analysis, EEG data were downsampled to 1 kHz.

### Image registration and preprocessing

Raw data were first automatically registered across days using the first recording as a target for alignment. In cases where automatic registrations failed, we manually registered the data. The dorsal cortex was then tiled with 818 ROIs each with a fixed location relative to bregma and lambda. This resulted in an average ROI size of 25 by 25 pixels (approximately 230 by 230 μm). ROIs were also registered against the Allen Mouse Brain Common Coordinate Framework (CCFv3-2017) (Wang et al. 2020). Fluorescence traces for each ROI were calculated by averaging all pixel values. Slow drifts were removed from the traces by high-pass filtering with an intensity-preserving algorithm: A filtered trace was generated using the 8^th^ percentile computed in a moving window of 62.5 s. This filtered trace was median subtracted and then subtracted from the raw trace to generate a drift corrected version. Activity was calculated as the ΔF/F_0_, where F_0_ was the median fluorescence of the entire trace.

### Behavioral state classification

Quiet rest was defined as periods where the mouse’s running speed, smoothed with a 1 s moving-average window, was below 0.33 cm/s. Transitions were excluded by discarding 100 ms flanking each rest epoch boundary. Unless otherwise noted, all analyses used only data from quiet rest periods. For optogenetic stimulation analyses, all behavioral states were included to maximize trial counts.

### Head-twitch response quantification

Mice were placed in a circular arena for 15 min after DOI injection and recorded in darkness under infrared illumination with a top-view behavioral camera at 60 Hz frame rate. Head-twitches were detected in these recordings by manual annotation.

### Analysis and visualization of correlation and variance

To minimize the contribution of hemodynamic effects, all correlation and variance analyses were performed on 2-8 Hz band-passed data. When visualizing data with a box plot, the central box indicates the interquartile range (IQR), spanning from the first (Q_1_) to the third (Q_3_) quartiles. Whiskers extend to data points not considered outliers. Outliers were defined as data points lying outside of the Q_1_ – 1.5 IQR to Q_3_ + 1.5 IQR range. A central horizontal line indicates the median of the distribution. To generate correlation seed maps we calculated the correlation of all ROIs with a given seed ROI. These correlation values were then visualized as a heatmap. To generate seed correlation contour maps (**Figure 1D, 2D, 3D, S11**), we generated isocontour lines for each of the 818 seed correlation maps at the level corresponding to 35% of the range between its minimum and maximum correlation values. All 818 isocontour lines were then overlaid to generate a seed correlation contour map. Average within and between domain correlations were computed as the average correlation between all possible pairs of ROIs within or between the domains.

### Spectral analysis

To calculate power spectra during the periods of quiet rest we only used contiguous periods of quiet rest of at least 4 s duration. This was done to avoid introducing stitching artifacts in the power spectra. Spectra were calculated for all ROIs separately and the resulting spectra averaged. For EEG data the signals were first common average re-referenced by subtracting the mean signal over all electrodes from each electrode.

### Clustering

For spectral clustering (**Figure 1E, S1**) we generated the average correlation matrix for all ROIs by computing individual correlation matrices for individual mice and averaging resulting correlation matrices. Then the clusters were computed using MATLAB’s function *spectralcluster* with the average correlation matrix as precomputed distance. ROIs for which silhouette coefficients were less than 0.5 were excluded from the clusters for all subsequent analyses (132 ROIs out of 818). To estimate the uncertainty in the mean silhouette coefficient (**Figure S1B, D**), we used a bootstrap over mice. For each of 10 000 bootstrap replicates, we resampled mice with replacement, computed the average correlation matrix across the resampled mice, clustered the ROIs, and computed the mean silhouette coefficient across ROIs for each cluster number. The error bars (bootstrap SEM) are the standard deviation of these mean silhouette coefficients across replicates.

### First cortical gradient computation

To compute the first gradient of the correlation matrix (**Figure 1F**), we fitted the *GradientMaps* model from the BrainSpace MATLAB package (Vos de Wael et al. 2020) to the average correlation matrix for combined baseline and recovery conditions using default parameters (normalized angle kernel with diffusion embedding approach).

### Onset responses

For each event type (grating, optogenetic stimulation, and fidget), for each ROI and mouse we averaged the responses across trials, and subtracted the mean baseline activity from the trial-averaged response; these trial-averaged, per-ROI-per-mouse responses formed the lowest-level unit for the hierarchical bootstrap. Optogenetic responses used the same −0.5 to 0 s baseline. Fidget events were defined as brief increases in running speed from rest (average smoothed speed had to be below 0.17 cm/s from −2.5 to 0 s, above 0.5 cm/s from 0 to 0.1 s, below 0.5 cm/s from 0.5 to 1.5 s after the event), with a baseline window of −2 to −1.5 s.

### Simulated uniform vascular modulation

To test whether a uniform increase in vascular fluctuations could account for the observed correlation changes (**Figure S4**), we added a synthetic global normal noise ξ of varying amplitude to the non-drug GCaMP data and asked whether this could reproduce the DOI-induced correlation increase. For each mouse we scaled a common-mode signal by a coefficient α and added it to the no-drug data, using a single α shared between baseline and recovery.

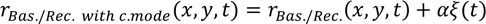

We then found the α at which the mean correlation on a fitting block (either AL–PM or PM–PM), averaged across the common-mode-injected baseline and recovery, equals the mean correlation of that block under DOI. This yields the common-mode amplitude needed to match the observed between-domain correlation increase. For mice which already had higher correlations for baseline and recovery than DOI, α was set to zero (two out of eleven mice for fitting to PM-PM correlations). We then compared the within-domain correlations produced by this simulation against those observed under DOI.

### Neuromodulatory projections analysis

Axonal projection density volumes were obtained from the Allen Mouse Brain Connectivity Atlas. For each neuromodulatory system, we selected experiments based on injection target and, where available, the appropriate Cre driver line: serotonergic line Slc6a4-Cre injections in dorsal raphe (DR); cholinergic line Chat-IRES-Cre injections in the Diagonal band nucleus (NDB) and in substantia innominata (SI) (analyzed separately); and, for ventral tegmental area (VTA) projections, all available lines targeting VTA, as no DAT-Cre experiments were available. To minimize contamination by fibers of passage, which appear in the atlas as spatially discontinuous or focally patchy cortical labeling rather than smoothly graded terminal innervation, we visually inspected the cortical projection patterns of each candidate experiment and excluded those lacking spatial continuity, retaining only experiments whose cortical labeling exhibited the smoothly graded distribution expected of terminal projections. For each retained experiment (5 out of 7 for serotonergic projections, 3 out of 3 for cholinergic projections from NDB, 8 out of 8 for cholinergic projections from SI, 6 out of 15 for projections from VTA), the projection density was normalized by injection density, and the resulting volumes were averaged across experiments. The averaged cortical volumes were then mean-projected onto a top-view representation of the cortical surface and averaged within the same 818 ROIs used for widefield imaging (**Figure 1I**). For visualization, the color scale for each projection density map was capped at the 99th percentile of values to prevent a small number of outlier ROIs from dominating the display.

### Mapping of AL- and PM-projecting thalamic regions

To identify thalamic locations preferentially projecting to the AL or PM cortical domains and guide our thalamic injections (**Figure S8**), we used cortical projection and injection density volumes from the Allen Mouse Brain Connectivity Atlas. For the analysis we used both wild-type experiments and all transgenic lines which had primary injection sites in thalamus. For each experiment, the cortical projection density volume was projected onto a top-view representation of the dorsal cortex and registered to the reference coordinates of our widefield recordings (**Figure 1C**). We then computed correlation coefficient between this top-view projection map and the AL–PM binary mask derived from our clustering of ROIs into the AL and PM domains (**Figure 1E**). This yielded, for each experiment, a single scalar with positive values indicating preferential projection to AL and negative values indicating preferential projection to PM. To then map this AL-vs-PM bias back into thalamus, we computed a weighted average of the thalamic injection density volumes across all retained experiments, using this correlation coefficient as the per-experiment weight.

### Statistical analysis

All quantitative information for the statistical analyses performed in this manuscript is provided in **Table S1**. For all statistical tests involving hierarchically structured data we used a hierarchical bootstrapping approach to account for the dependencies between groups of datapoints (Saravanan et al. 2020). In these cases, the data were first resampled with replacement at the level of mouse or recording site and then resampled with replacement at the level of ROIs or neurons. We performed 10 000 bootstrap repetitions and calculated a distribution of bootstrapped mean values. The p-value was calculated as the fraction of the bootstrap sample consistent with the null hypothesis. In the case of paired data (e.g. recordings from the same mice for DOI and non-DOI conditions), the paired version of hierarchical bootstrap test was used. For this, we computed the difference in mean values between pairs of bootstrap samples. For comparisons involving different groups of animals (e.g. ketanserin-treated vs. other conditions), a non paired hierarchical bootstrap was used instead. To quantify the significance of the difference of two average calcium responses as a function of time, hierarchical bootstrap was performed for every time bin. For visual clarity, a contiguous cluster of significant bins was marked only if it spanned at least 2% of the bins along the x axis, with a minimum of two bins; isolated significant bins (and shorter clusters) were left unmarked.

---

**Figure S1.**
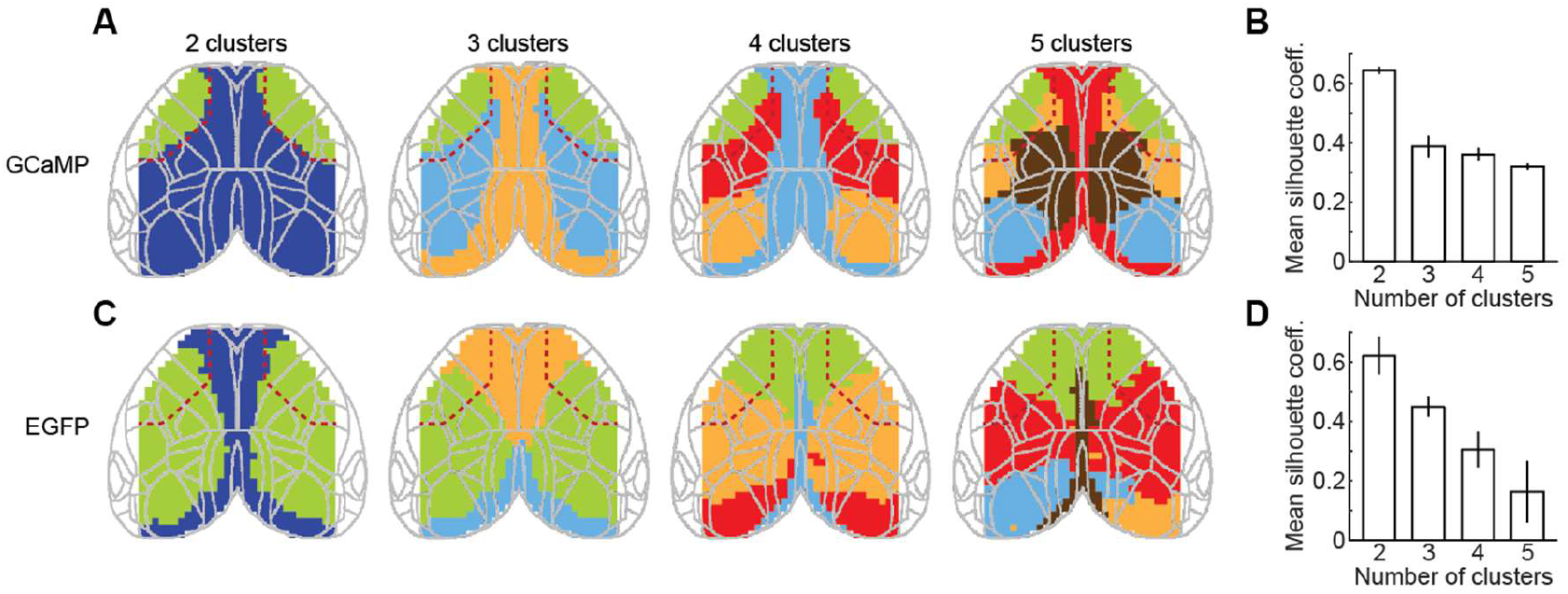
Hemodynamics filtered in 2-8 Hz cannot explain the partitioning of the ROIs into AL and PM domains. (**A**) Spectral clustering of the average GCaMP correlation matrix into 2 to 5 clusters. (**B**) Mean silhouette coefficient as a function of the number of clusters. Error bars denote the bootstrap SEM (**Methods**). (**C**) As in **A**, but for EGFP data. (**D**) As in **B**, but for EGFP data.

**Figure S2.**
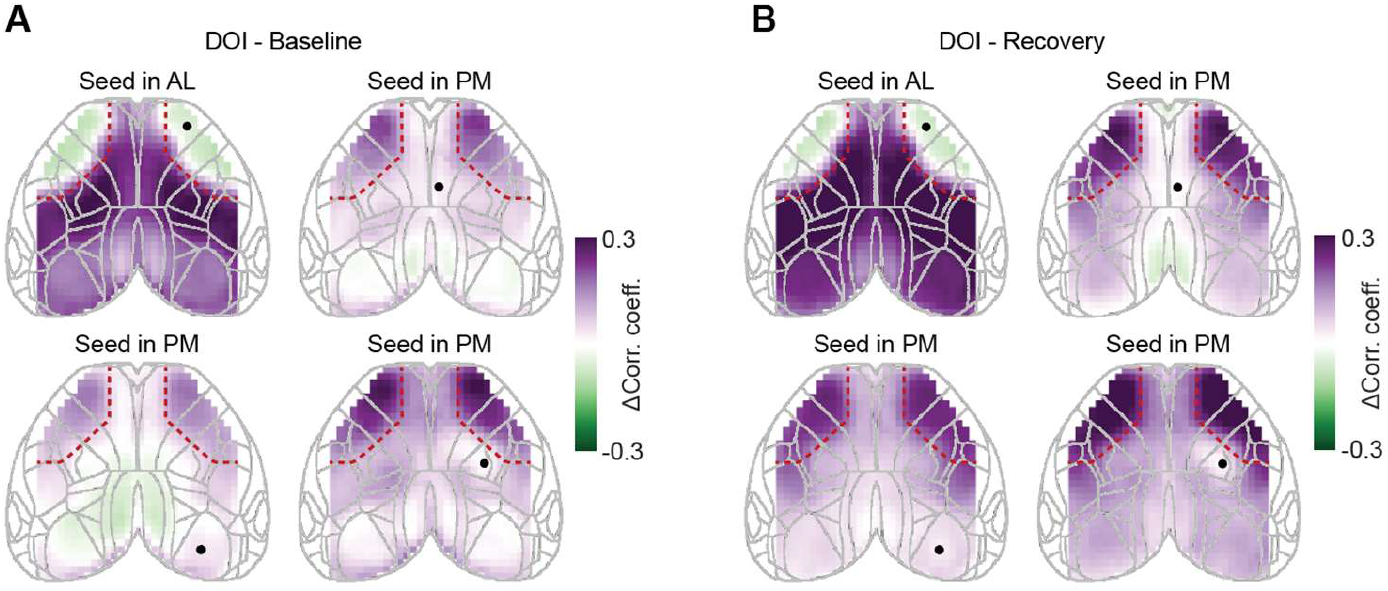
Difference of seed correlation maps between DOI and either baseline or recovery conditions. (**A**) Difference in seed correlation maps for four selected seeds between DOI and the baseline condition. (**B**) As in **A**, but between DOI and the recovery condition.

**Figure S3.**
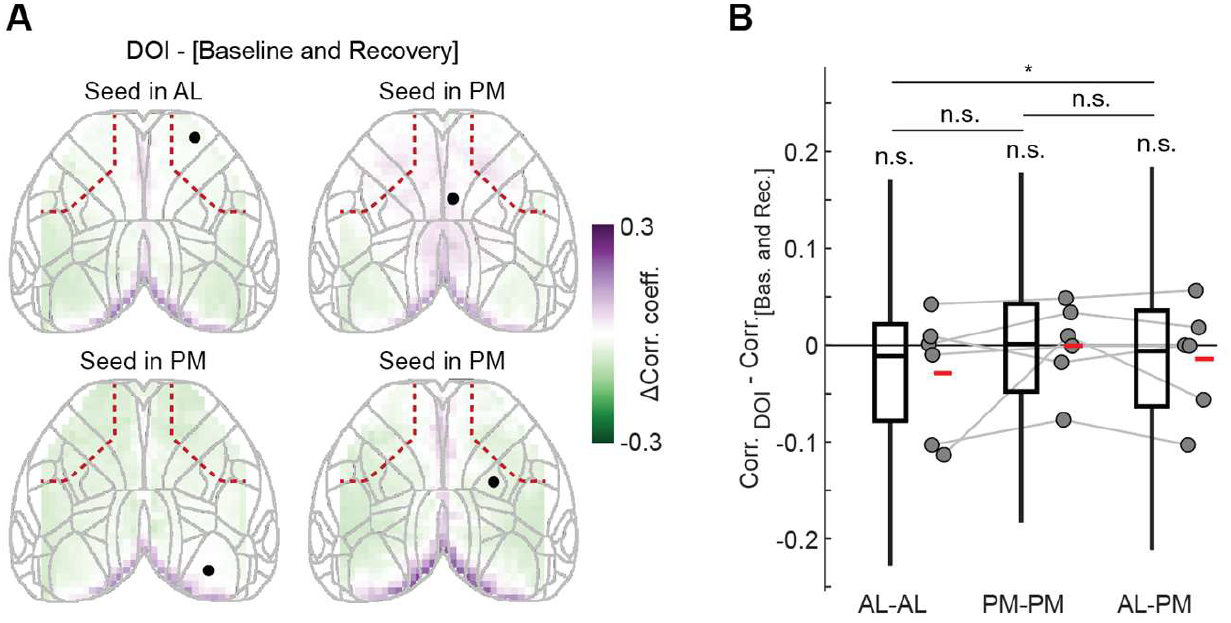
Hemodynamics cannot explain patterns of DOI-induced changes of correlations observed with GCaMP. (**A**) Difference in seed correlation maps for four example ROIs (black dots) between DOI and combined baseline and recovery conditions for mice expressing EGFP brain-wide. Limits are the same as for GCaMP data (Figure 2C). (**B**) Change in correlation of activities between DOI and combined baseline and recovery conditions for pairs of ROIs within the AL domain, within the PM domain, and between the AL and PM domains for mice expressing EGFP brain-wide. Boxplots show the distribution of differences in correlations for all pairs of ROIs within or between domains for all mice.

**Figure S4.**
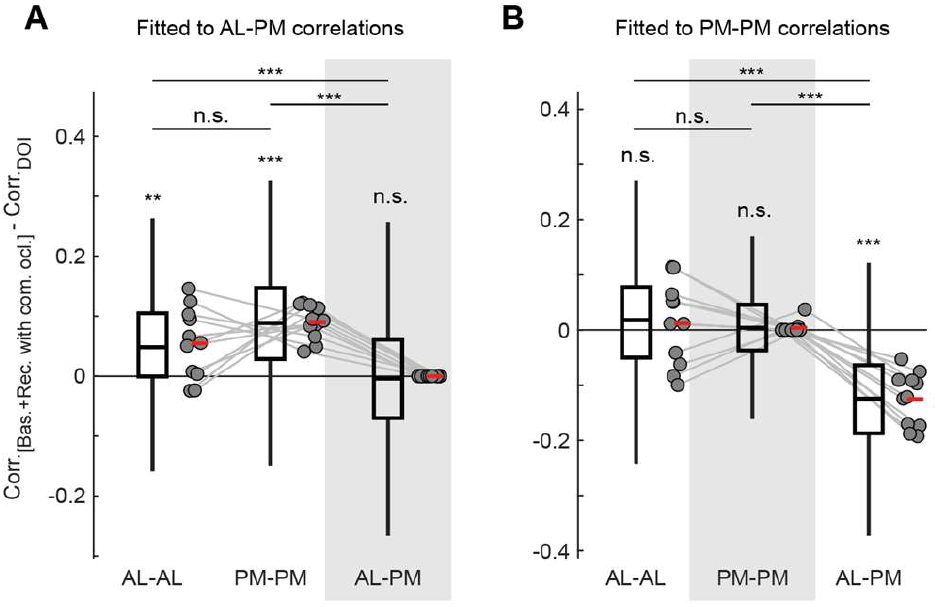
Addition of the common occluder cannot explain the pattern of the correlation change. (**A**) Difference between (i) the combined baseline and recovery activity correlations with an added common occluder, scaled by the optimal multiplier that matches AL-PM correlations between DOI and combined baseline and recovery (**Methods**), and (ii) the DOI correlations, shown for AL-AL, PM-PM, and AL-PM ROI pairs. Gray shading indicates a pair of domains, for which the AL-PM correlations were matched. (**B**) As in **A**, but for the optimal multiplier which matches PM-PM correlations between DOI and combined baseline and recovery.

**Figure S5.**
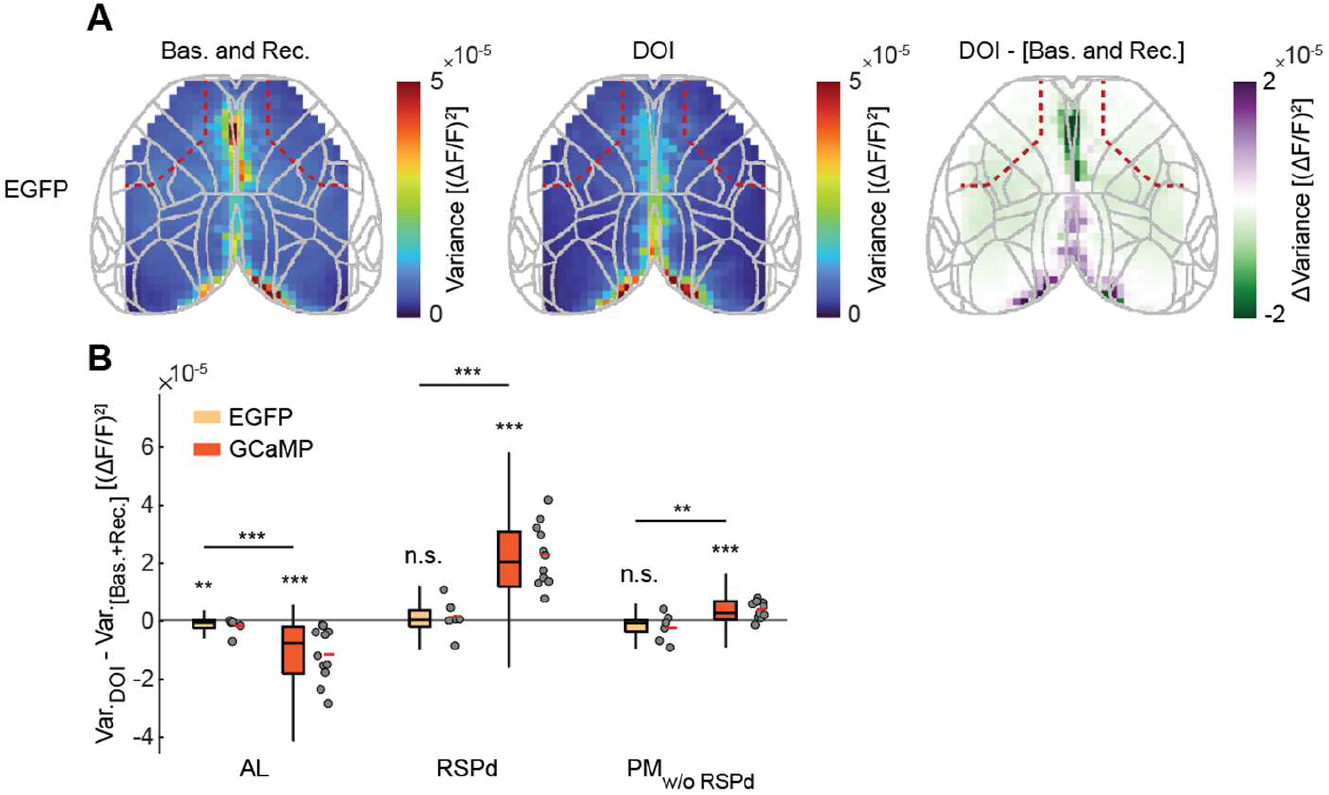
Hemodynamics cannot explain the DOI-induced changes in AL and RSPd variance. (**A**) Variance of the calcium activity for the combined baseline and recovery conditions, DOI, and the difference between the two for mice expressing EGFP brain-wide. (**B**) Difference between DOI variance and combined baseline and recovery variance for EGFP and GCaMP data for ROIs in AL, RSPd, and PM without RSPd. Boxplots show the distribution of variance across all ROIs within a given area of all mice.

**Figure S6.**
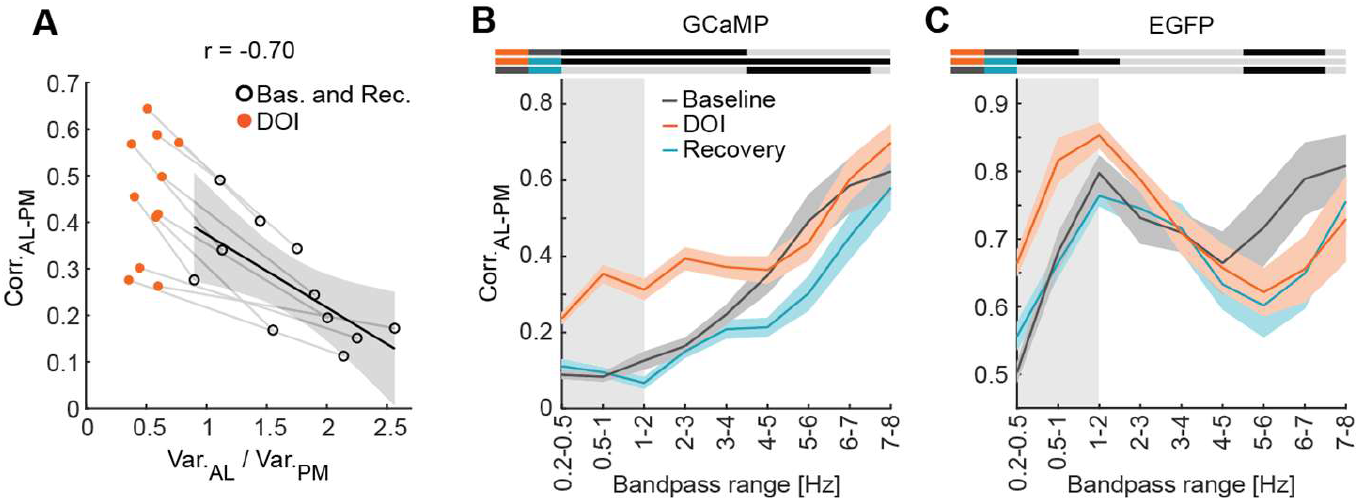
The level of the AL domain variance is associated with the strength of AL-PM correlations. (**A**) Correlation of the AL-to-PM variance ratio with the average correlation of activity between the AL and PM domains. Black open circles denote individual mice in the combined baseline and recovery conditions. Orange filled circles are data from the same mice in the DOI condition. Gray lines connect the same mice over conditions. Solid black line indicates the linear fit to the combined baseline and recovery datapoints. Gray shading indicates the 95% confidence interval of the fit. (**B**) Correlation between the AL and PM domains for GCaMP as a function of the frequency band for baseline, DOI, and recovery conditions. Gray shading indicates the region of the hemodynamic effects of DOI according to **C**. (**C**) As in **B**, but for EGFP data.

**Figure S7.**
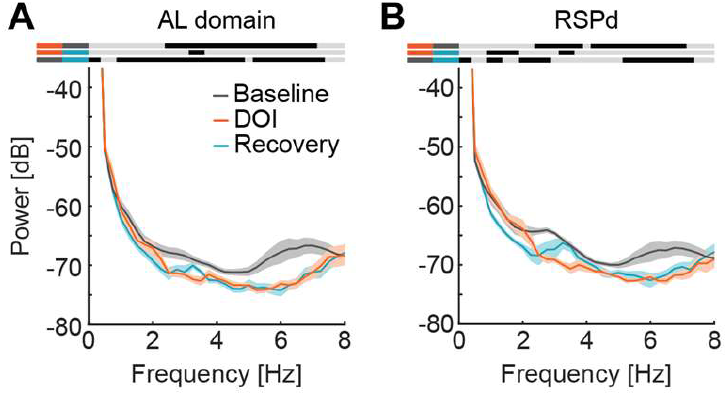
Hemodynamics cannot explain the change in the spectral power in the widefield recordings. (**A**) Power spectral density of the widefield calcium signal averaged for ROIs in the AL domain for baseline, DOI, and recovery conditions for mice expressing EGFP brain-wide. (**B**) As in **A**, but for ROIs in RSPd.

**Figure S8.**
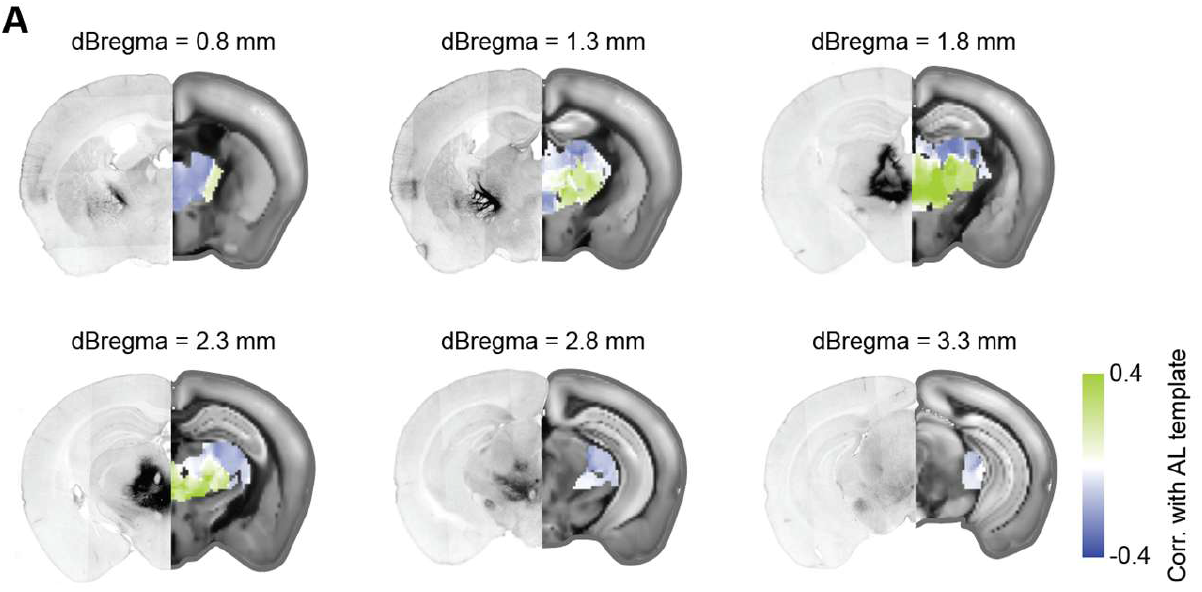
Injection sites and thalamic locations projecting to the AL and PM domains. (**A**) Identification of thalamic regions projecting preferentially to the AL or PM cortical domain, based on data from the Allen Mouse Brain Connectivity Atlas. Coronal sections at six distances posterior to bregma (dBregma). Left half of each section: Example histology image of an AAV injection targeting thalamic nuclei that project to the AL domain. Right half: Allen Mouse Brain Atlas template image overlaid with the thalamic AL–PM projection map. For each voxel, color indicates the AL-vs-PM bias of the thalamo-cortical projections originating from injections covering that voxel (**Methods**). Green: voxels where injections from the Allen Mouse Brain Connectivity Atlas labeled neurons projecting preferentially to AL; dark blue: the same, but projecting preferentially to PM.

**Figure S9.**
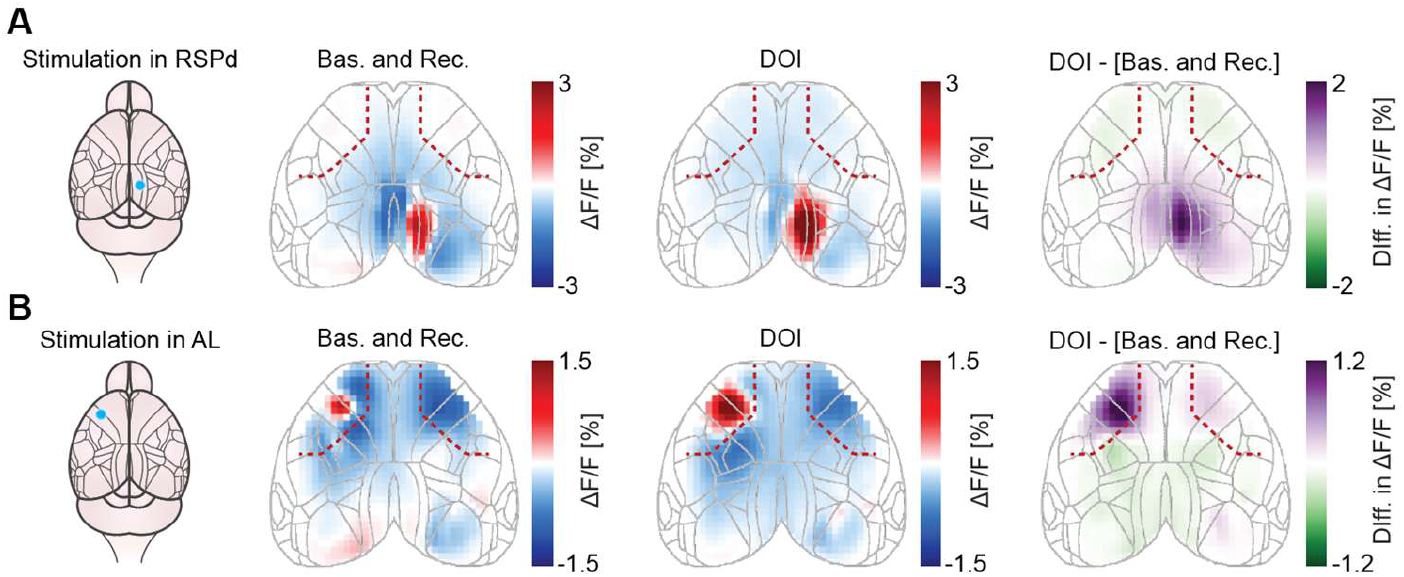
Spatial maps of responses to the optogenetic stimulation of the thalamic axons in RSPd and AL. (**A**) Left: Schematic of the stimulation location in RSPd. In these mice, an AAV injection was targeted to the AV thalamic nucleus. Right: Spatial maps of the calcium response to RSPd stimulation for the combined baseline and recovery condition, DOI condition, and the difference between the two in a 0.5-1 s window. (**B**) As in **A**, but for the stimulation in AL.

**Figure S10.**
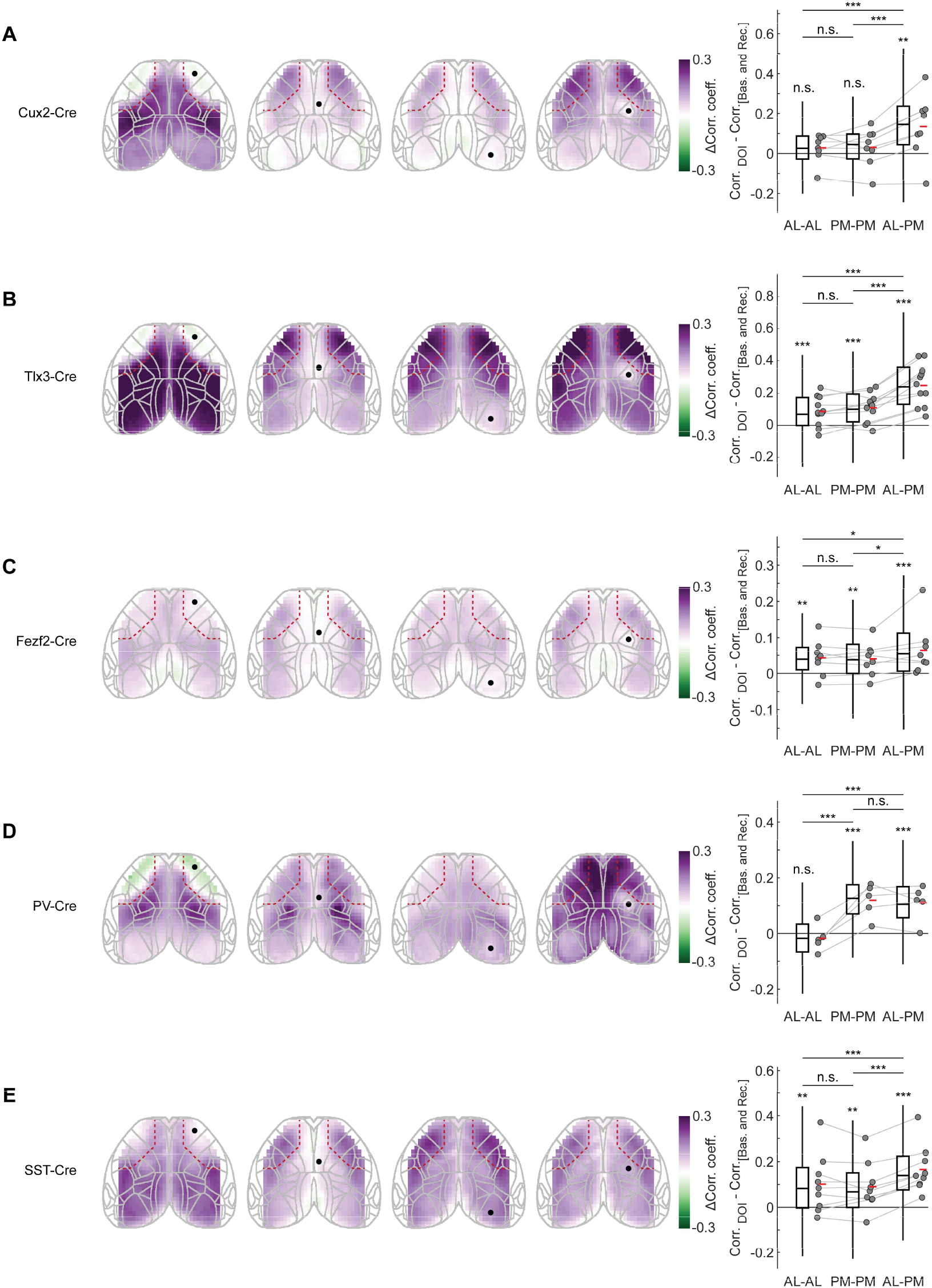
Patterns of correlation changes for different cell types. (**A**) Left: Difference in seed correlation maps for four example ROIs (black dots) between DOI and combined baseline and recovery conditions for Cux2-Cre mice with Cre-dependent GCaMP expression in a subset of L2/3 and L4 excitatory neurons. Right: change in correlation of activities between DOI and combined baseline and recovery conditions for pairs of ROIs within the AL domain, within the PM domain, and between the AL and PM domains. (**B**) As in **A**, but for Tlx3-Cre mice with Cre-dependent GCaMP expression in a subset of L5 IT excitatory neurons. (**C**) As in **A**, but for Fezf2-Cre mice with Cre-dependent GCaMP expression in a subset of L5 ET and L6 neurons. (**D**) As in **A**, but for PV-Cre mice with Cre-dependent GCaMP expression in PV neurons. (**E**) As in **A**, but for SST-Cre mice with Cre-dependent GCaMP expression in SST neurons.

**Figure S11.**
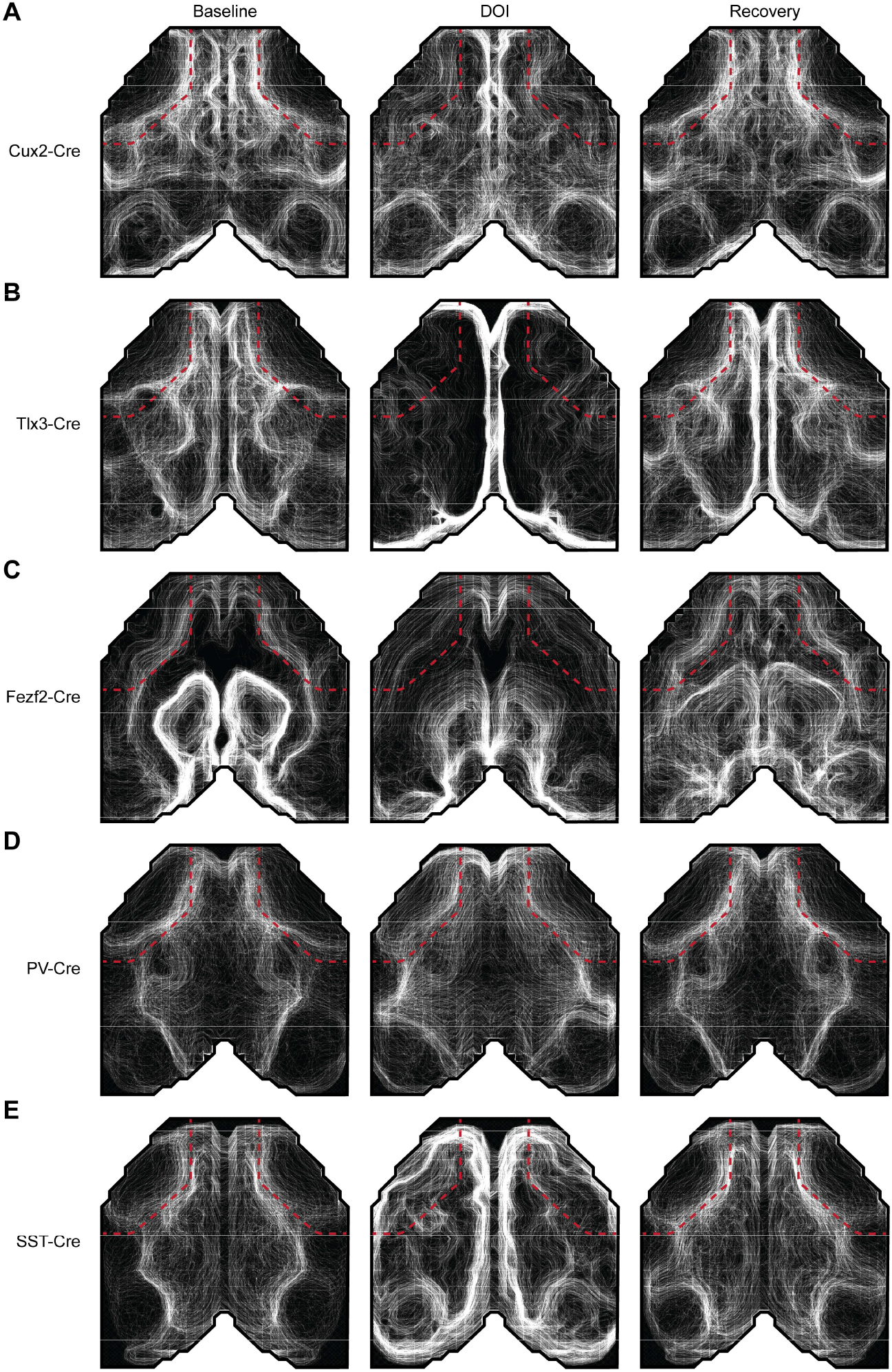
Seed correlation contour maps for different cell types. (**A**) Seed correlation contour maps for Cux2-Cre mice with Cre-dependent GCaMP expression in a subset of L2/3 and L4 excitatory neurons. (**B**) As in **A**, but for Tlx3-Cre mice with Cre-dependent GCaMP expression in a subset of L5 IT excitatory neurons. (**C**) As in **A**, but for Fezf2-Cre mice with Cre-dependent GCaMP expression in a subset of L5 ET and L6 neurons. (**D**) As in **A**, but for PV-Cre mice with Cre-dependent GCaMP expression in PV neurons. (**E**) As in **A**, but for SST-Cre mice with Cre-dependent GCaMP expression in SST neurons.

**Figure S12.**
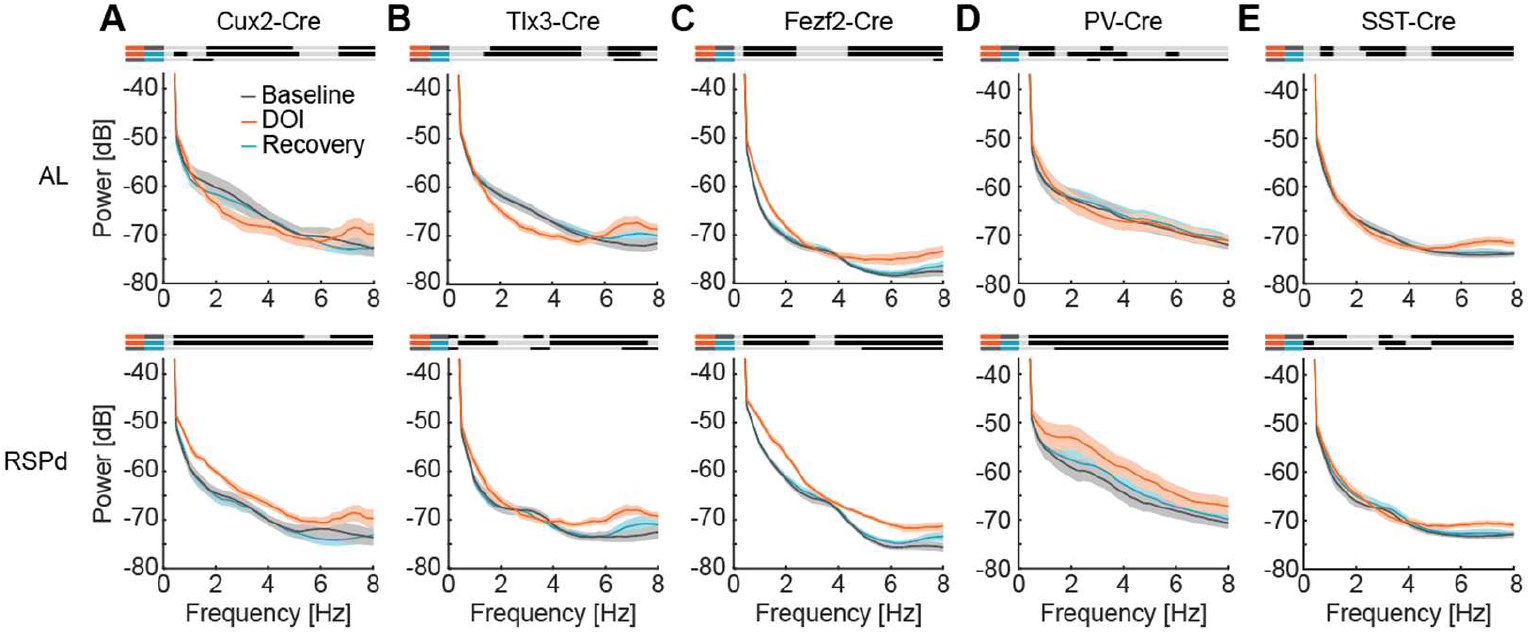
Effects of DOI on the power spectra for different cell types in the AL domain and RSPd. (**A**) Top: Power spectral density of the widefield calcium signal averaged for ROIs in the AL domain for baseline, DOI, and recovery conditions for Cux2-Cre mice with Cre-dependent GCaMP expression in a subset of L2/3 and L4 excitatory neurons. Bottom: The same as top, but for ROIs in RSPd. (**B**) As in **A**, but for Tlx3-Cre mice with Cre-dependent GCaMP expression in a subset of L5 IT excitatory neurons. (**C**) As in **A**, but for Fezf2-Cre mice with Cre-dependent GCaMP expression in a subset of L5 ET and L6 neurons. (**D**) As in **A**, but for PV-Cre mice with Cre-dependent GCaMP expression in PV neurons. (**E**) As in **A**, but for SST-Cre mice with Cre-dependent GCaMP expression in SST neurons.

**Figure S13.**
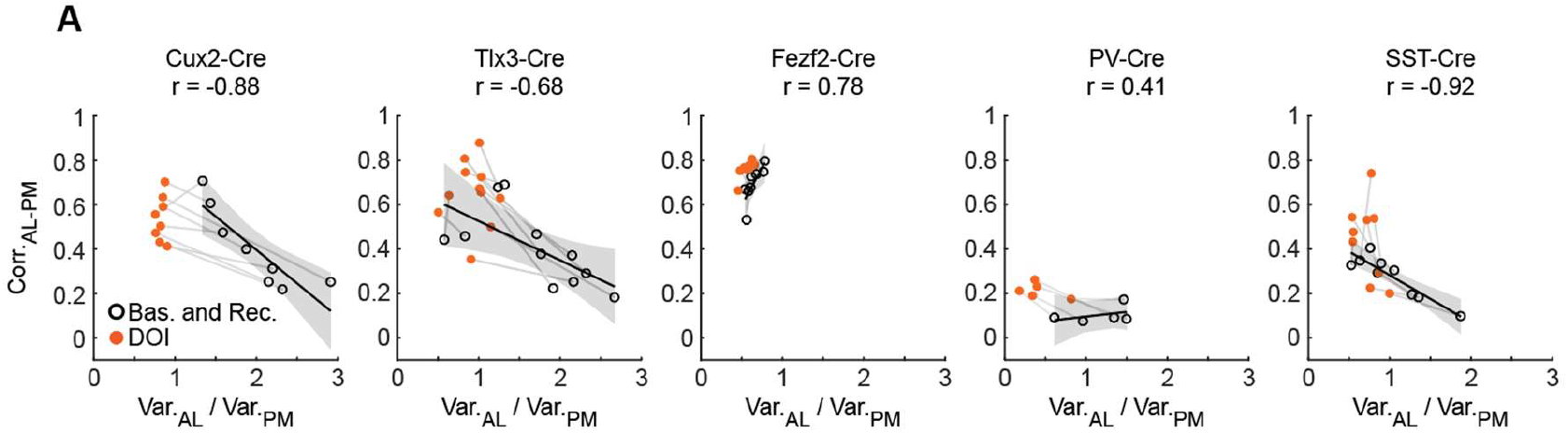
Correlation between AL-PM correlation and relative AL variance for different cell types. (**A**) Correlation of the AL-to-PM variance ratio with the average correlation of activity between the AL and PM domains for different cell types. Black open circles denote individual mice in the combined baseline and recovery conditions. Orange filled circles are the same mice in the DOI condition. Gray lines connect the same mice over conditions. Solid black lines indicate the linear fits to the combined baseline and recovery datapoints. Gray shading indicates the 95% confidence interval of the fits.

## Key Resource Table

| REAGENT or RESOURCE | Source | Identifier |
| --- | --- | --- |
| **Bacterial and virus strains** |  |  |
| AAV-PHP.eB-EF1α-GCaMP6s-WPRE (10<sup>14</sup> GC/ml) | FMI vector core | vector.fmi.ch |
| AAV-PHP.eB-EF1α-DIO-GCaMP6s (10<sup>15</sup> GC/ml) | FMI vector core | vector.fmi.ch |
| AAV-PHP.eB-EF1α-EGFP-WPRE (10<sup>14</sup> GC/ml) | FMI vector core | vector.fmi.ch |
| AAV-PHP.eB-hSyn-FLEX-jGCaMP8s-WPRE (10<sup>15</sup> GC/ml) | FMI vector core | vector.fmi.ch |
| AAV-PHP.eB-hSyn-PinkyCaMP (10<sup>14</sup> GC/ml) | FMI vector core | vector.fmi.ch |
| AAV-PHP.eB-Syn-ChR2(H134R)-GFP (10<sup>14</sup> GC/ml) | FMI vector core | vector.fmi.ch |
| AAV-2/1-Syn-ChR2(H134R)-GFP (10<sup>13</sup> GC/ml) | FMI vector core | vector.fmi.ch |
| **Chemicals, peptides, and recombinant proteins** |  |  |
| Fentanyl citrate | Actavis | CAS 990-73-8 |
| Midazolam (Dormicum) | Roche | CAS 59467-96-8 |
| Medetomidine (Domitor) | Orion Pharma | CAS 86347-14-0 |
| Ropivacaine | Presenius Kabi | CAS 132112-35-7 |
| Lidocaine | Bichsel | CAS 137-58-6 |
| Buprenorphine hydrochloride (Ethiq Xr) | Fidelis Pharmaceuticals | CAS 53152-21-9 |
| Ophthalmic gel (Humigel) | Virbac | N/A |
| Flumazenil (Anexate) | Roche | CAS 78755-81-4 |
| Atipamezole (Antisedan) | Orion Pharma | CAS 104054-27-5 |
| N-Butyl-2-cyanoacrylate (Vetbond) | 3M | CAS 6606-65-1 |
| Dental cement (Paladur) | Heraeus Kulzer | CAS 9066-86-8 |
| Mounting Medium (VectaShield Antifade) | Vector Laboratories | N/A |
| 2,5-dimethoxy-4-iodoamphetamine (DOI) hydrochloride | Sigma-Aldrich | CAS 42203-78-1 |
| Ketanserin tartrate | Sigma-Aldrich | CAS 83846-83-7 |
| Tamoxifen food | Envigo | CAS 10540-29-1 (TD.55125) |
| **Deposited data** |  |  |
| All mouse data and code used to generate manuscript figures | This paper | zenodo.org |
| **Experimental models: Organisms/strains** |  |  |
| Mus musculus: C57BL/6 | Charles River | N/A |
| Mus musculus: B6(Cg)-Cux2<sup>tm3.1(cre/ERT2)Mull</sup>/Mmmh | MMRRC | RRID:MMRRC_032779-MU |
| Mus musculus: Tg(Tlx3-cre)PL56Gsat/Mmucd | MMRRC | RRID:MMRRC_041158-UCD |
| Mus musculus: Fezf2<sup>tm1.1(cre/ERT2)Zjh</sup>/J | Jackson Laboratories | RRID:IMSR_JAX:036296 |
| Mus musculus: Pvalb<sup>tm1(cre)Arbr</sup> | Jackson Laboratories | RRID:IMSR_JAX:008069 |
| Mus musculus: Sst<sup>tm2.1(cre)Zjh</sup> | Jackson Laboratories | RRID:IMSR_JAX:018973 |
| Mus musculus: lgs7<sup>tm148.1(tetO-GCaMP6f,CAG-tTA2)Hze</sup>/J | Jackson Laboratories | RRID:IMSR_JAX:030328 |
| Mus musculus: Gt(ROSA)26Sor<sup>tm32(CAG-COP4*H134R/EYFP)Hze</sup>/J | Jackson Laboratories | RRID:IMSR_JAX:024109 |
| **Software and algorithms** |  |  |
| MATLAB (2023b) | The MathWorks | RRID:SCR_001622 |
| LabVIEW | National Instruments | RRID:SCR_014325 |
| Python | python.org | RRID:SCR_008394 |
| Panda3D | panda3d.org | N/A |
| Arduino IDE | arduino.cc | N/A |

---

**Table S1.**
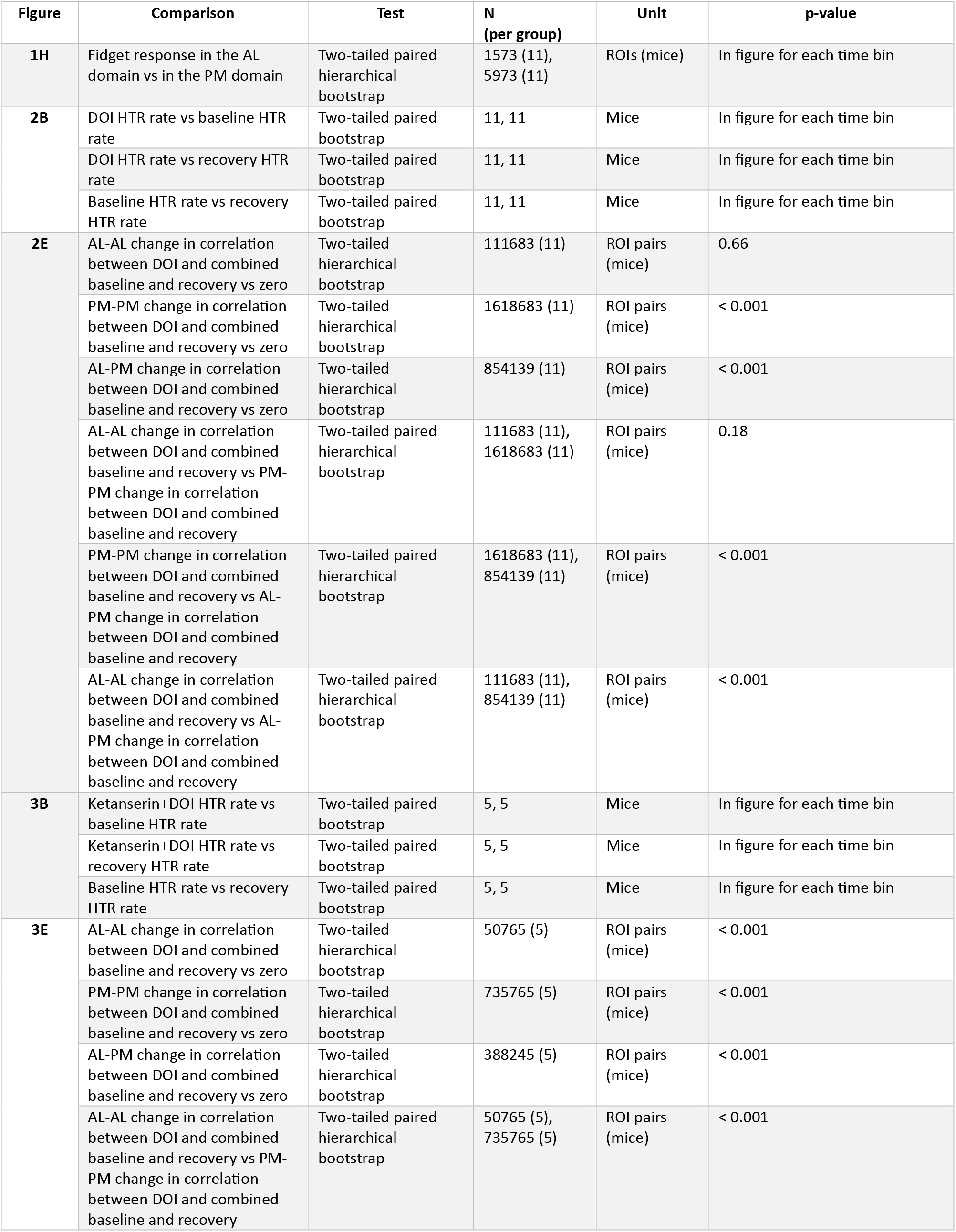
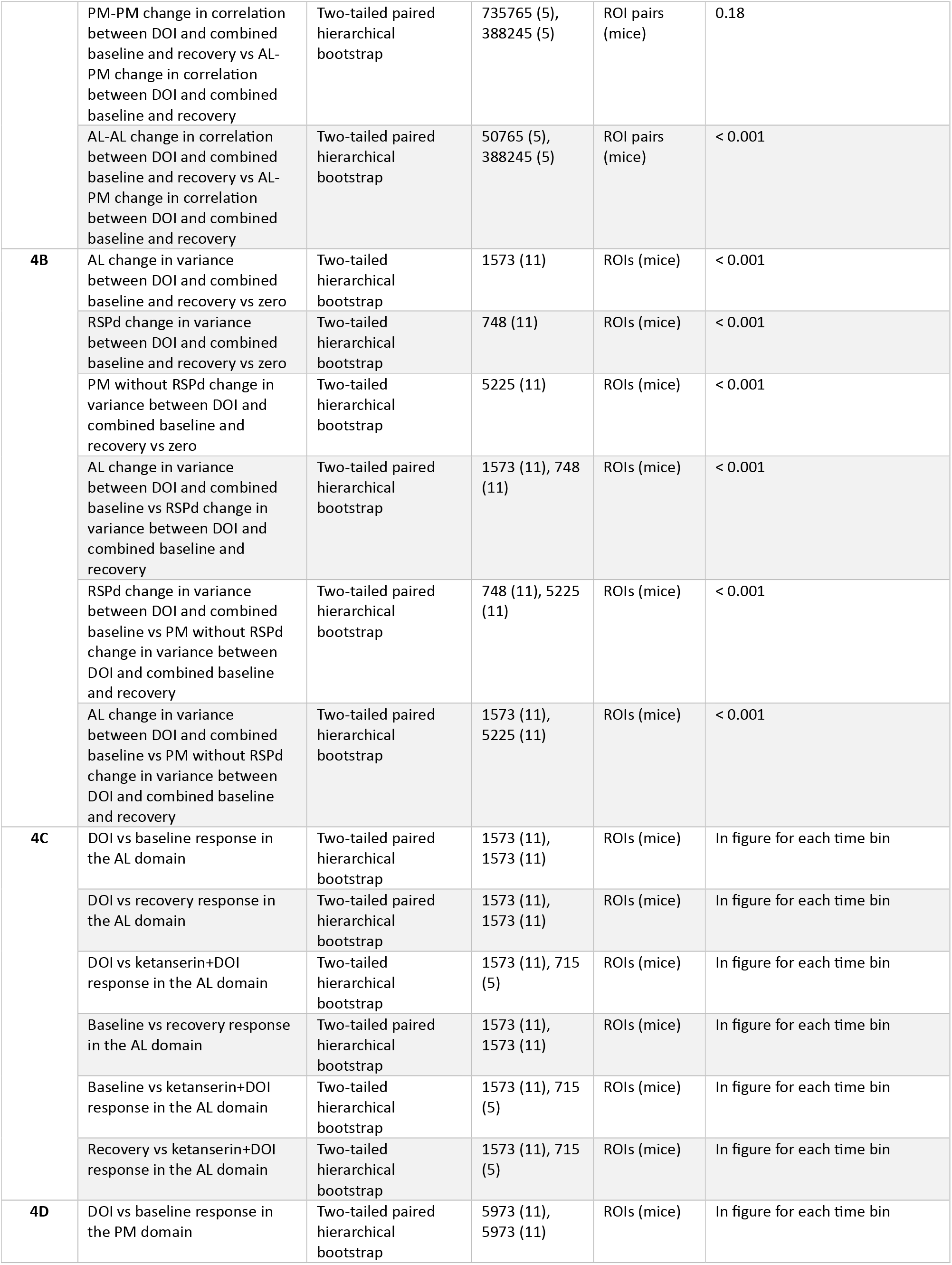
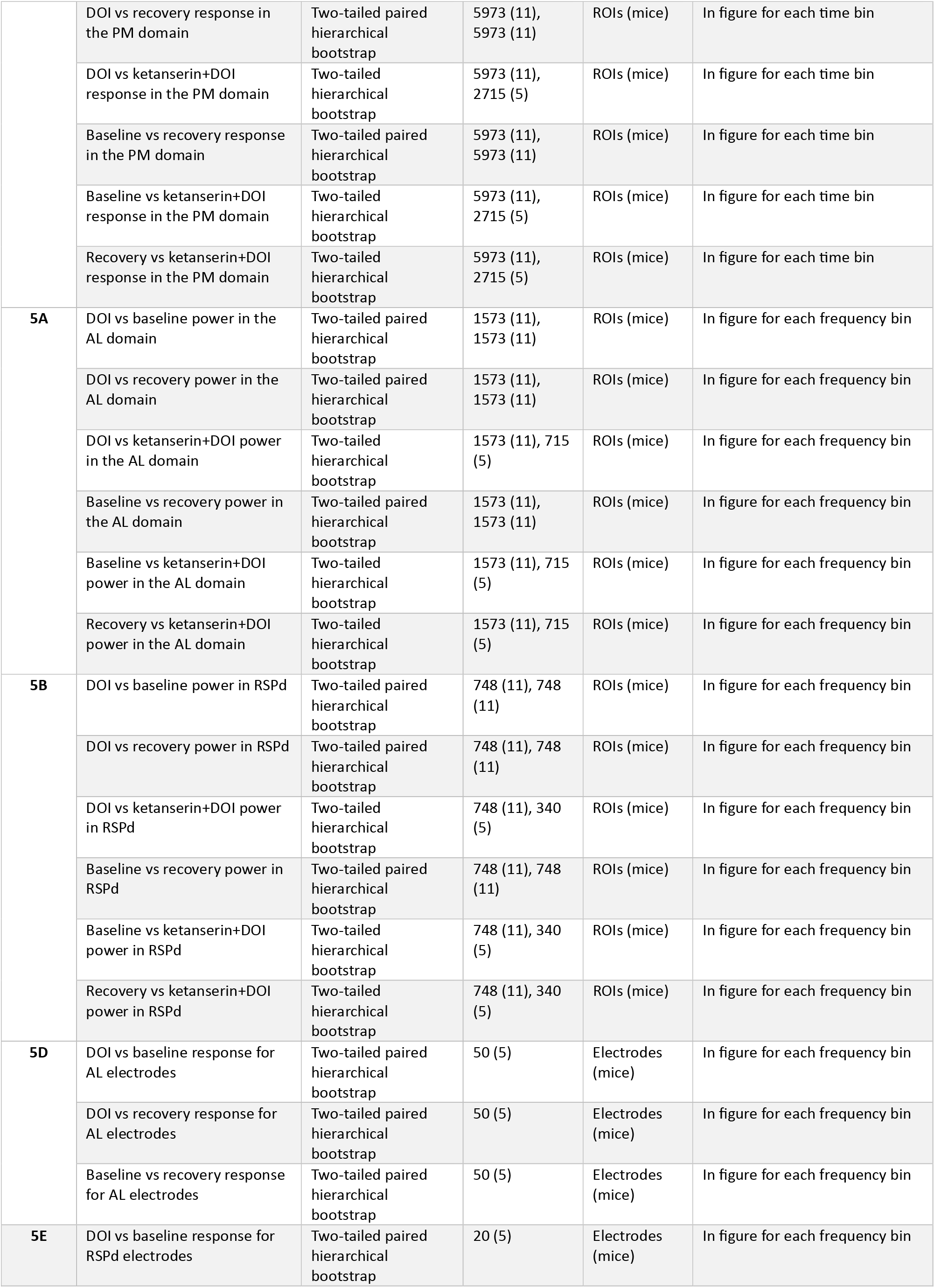
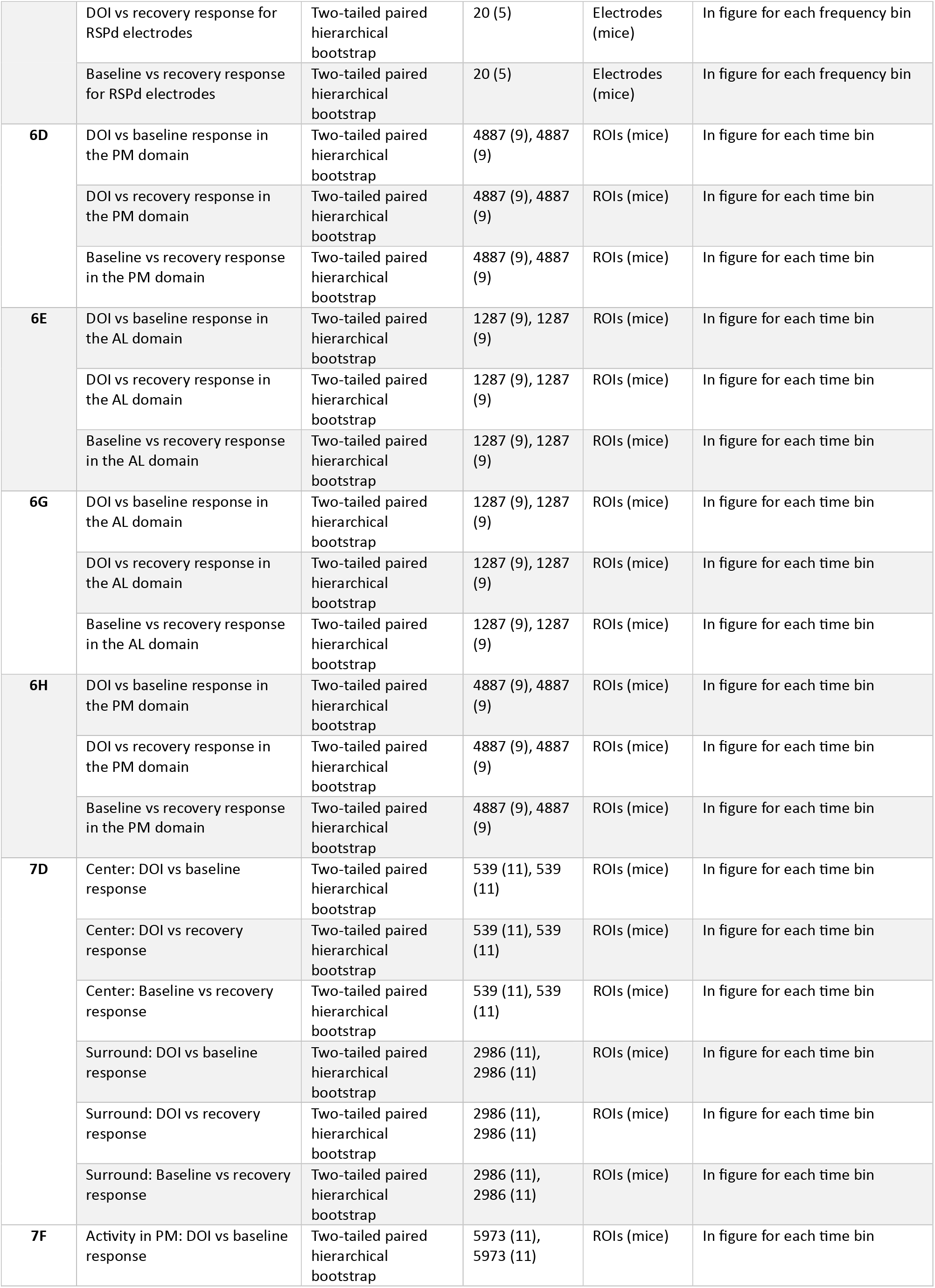
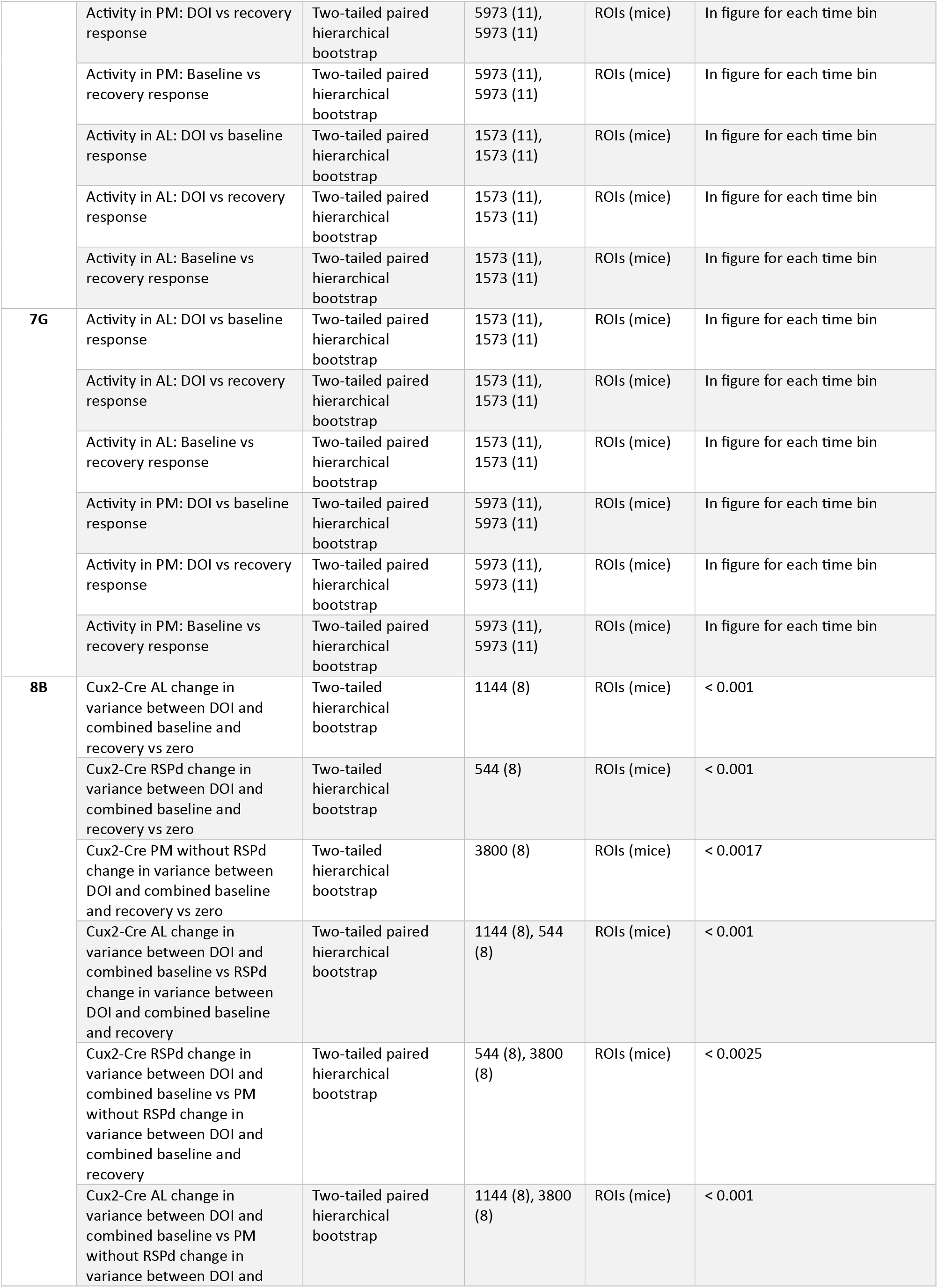
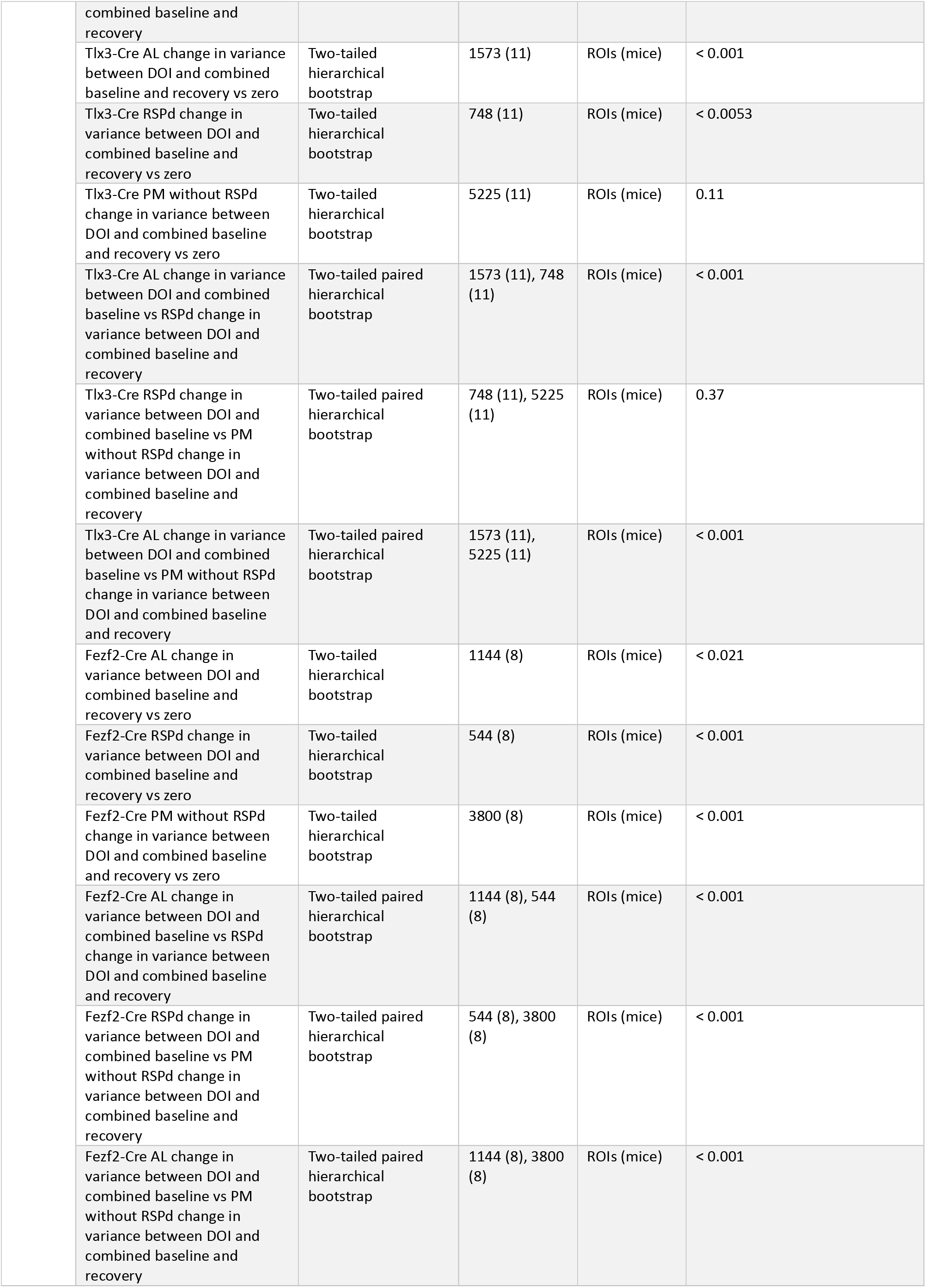
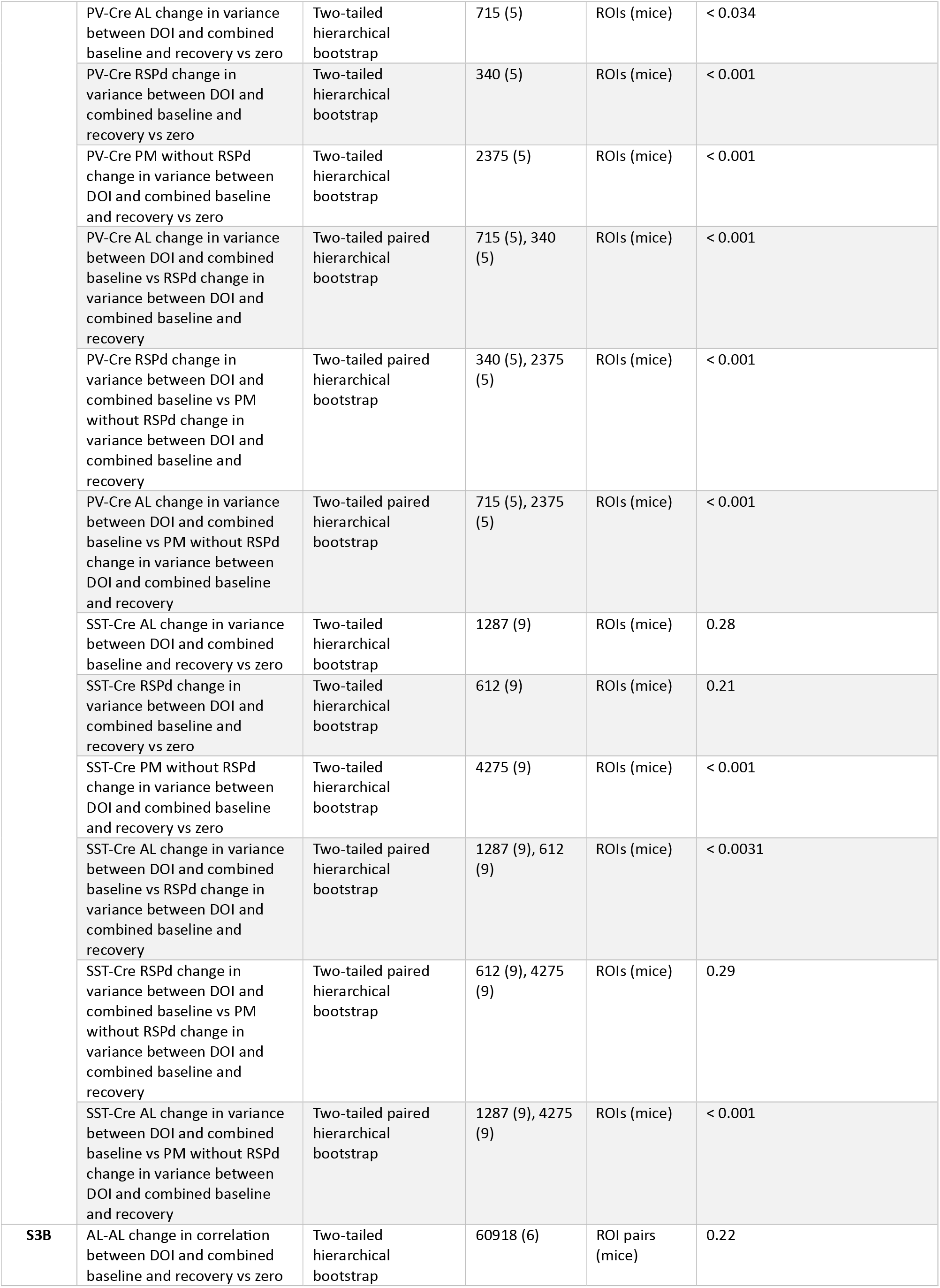
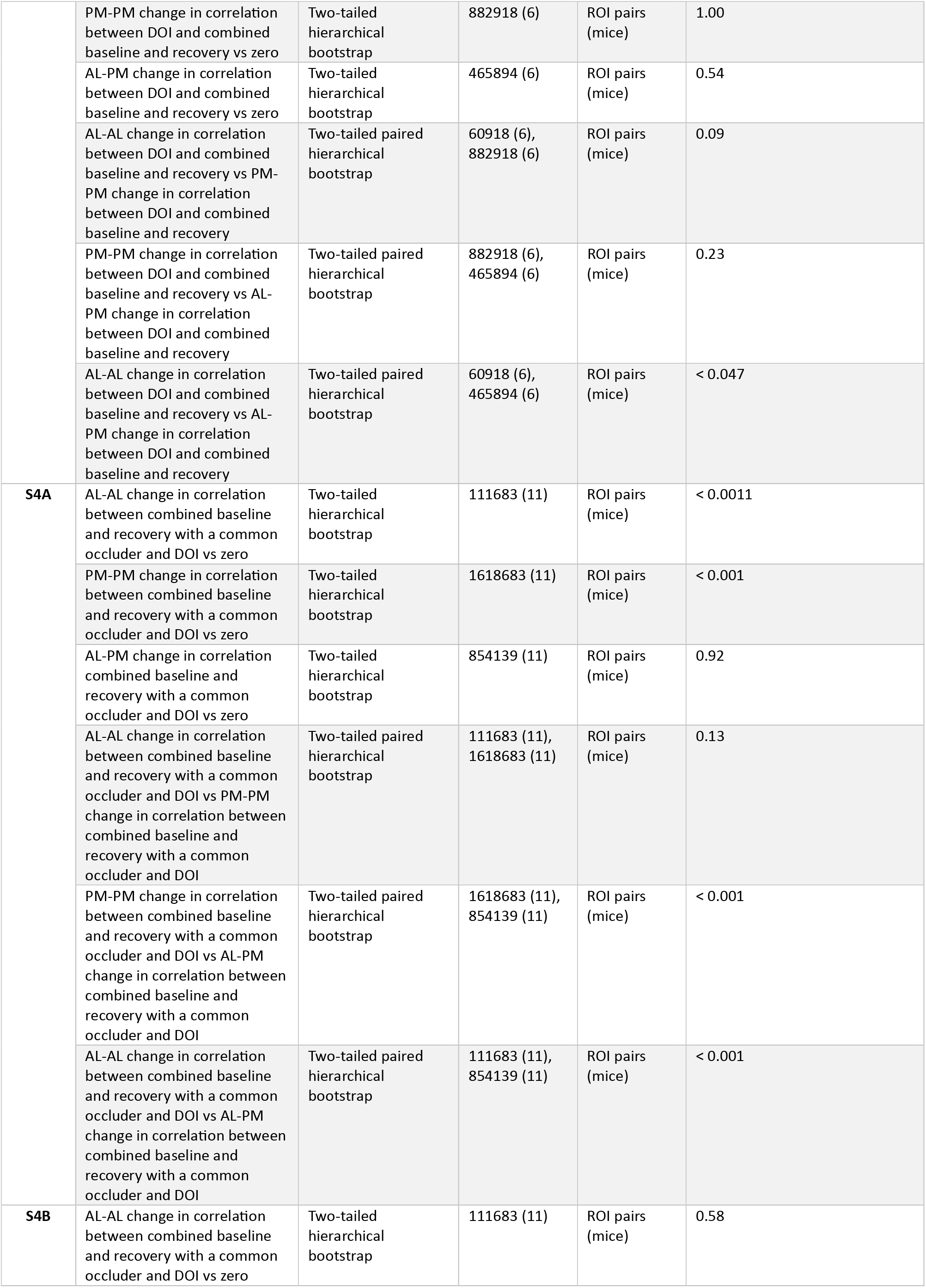
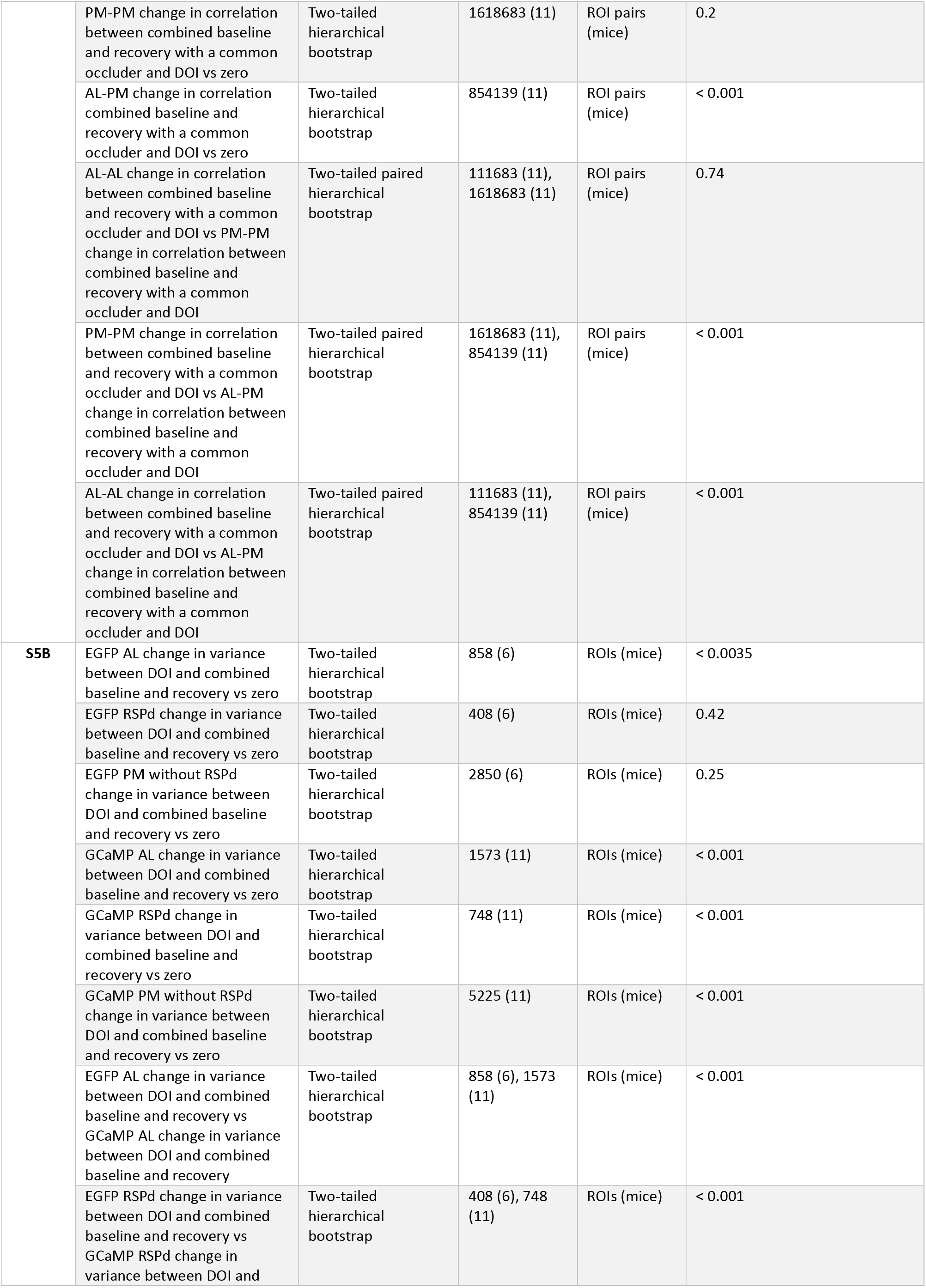
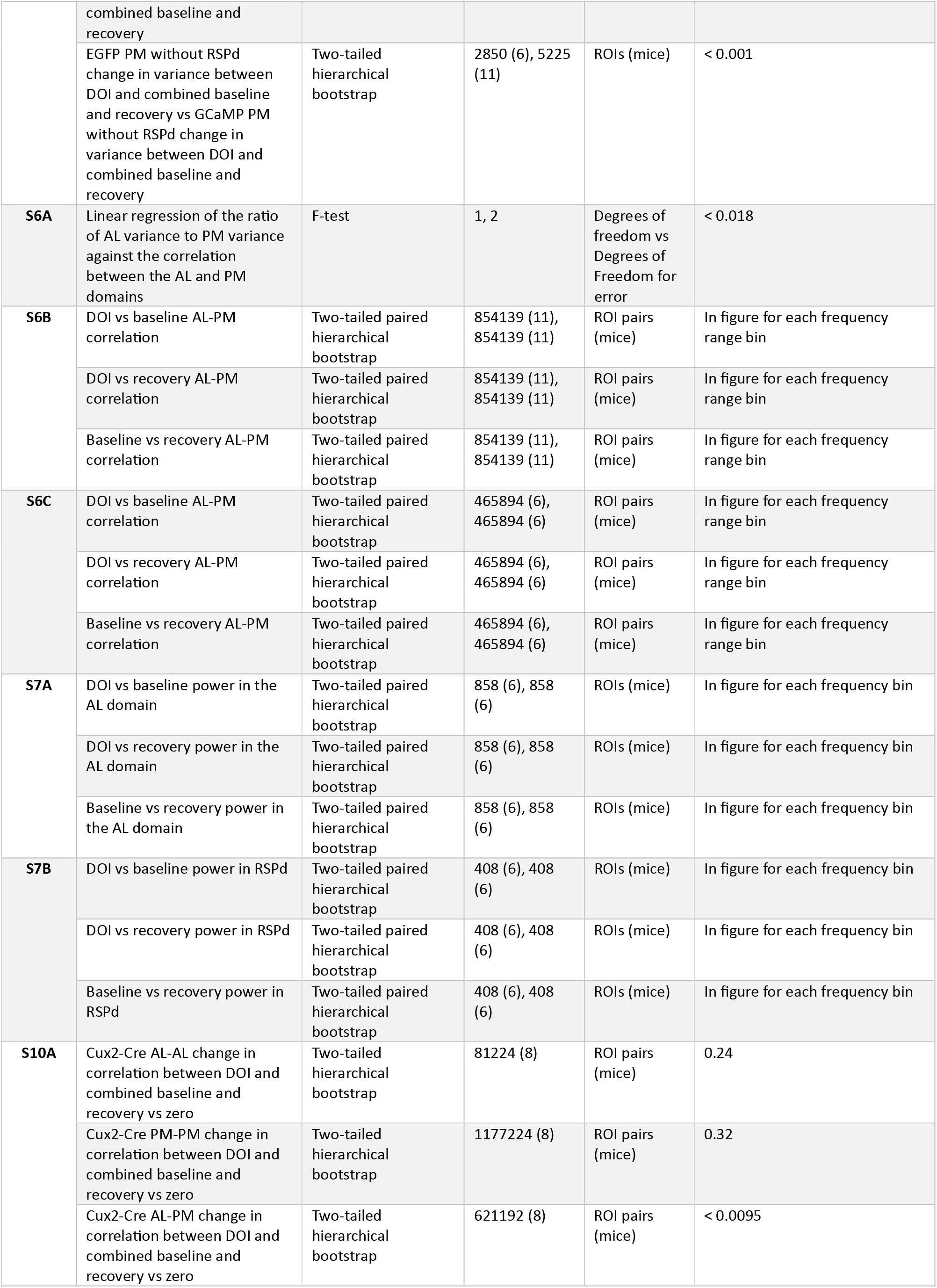
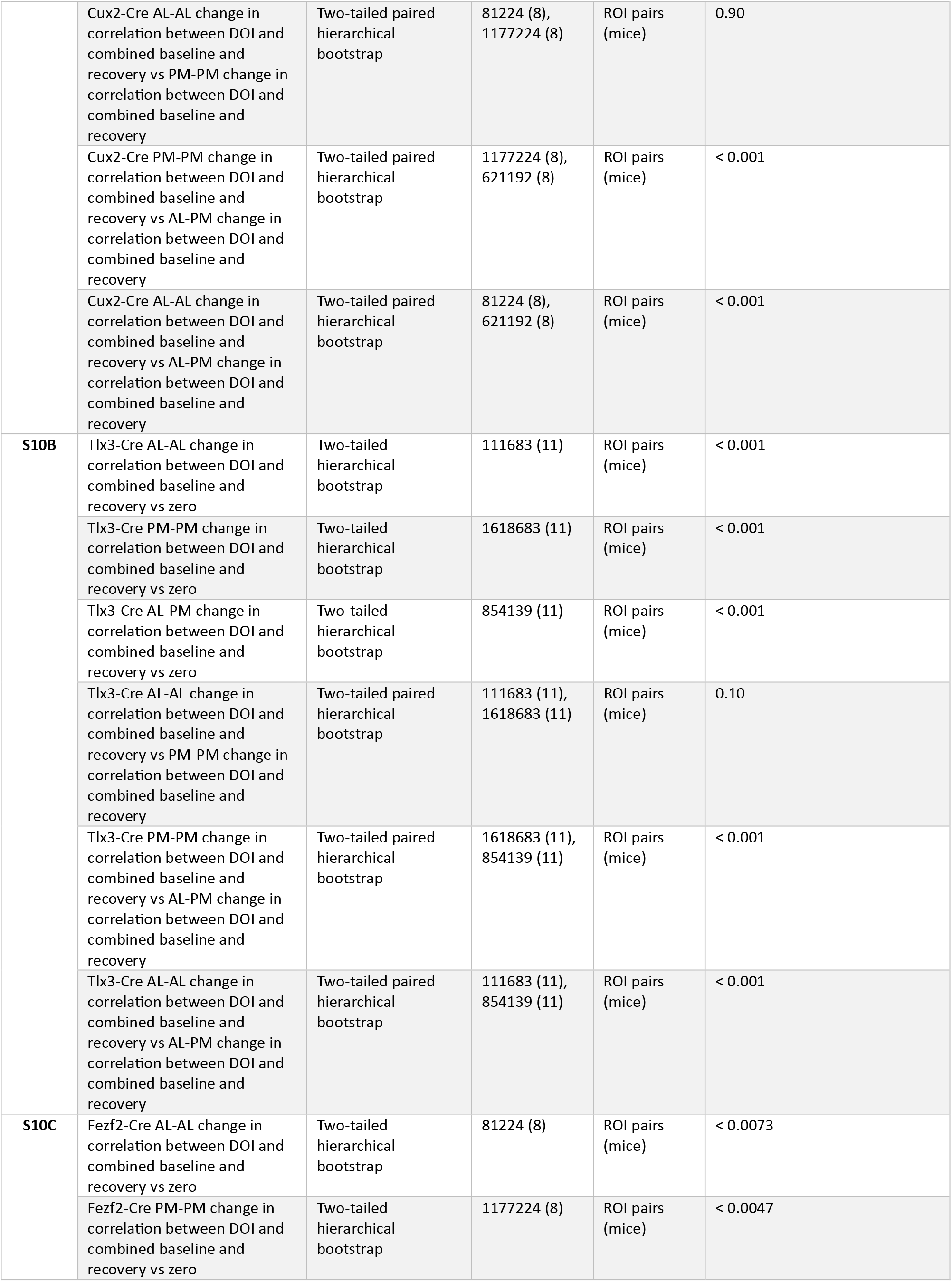
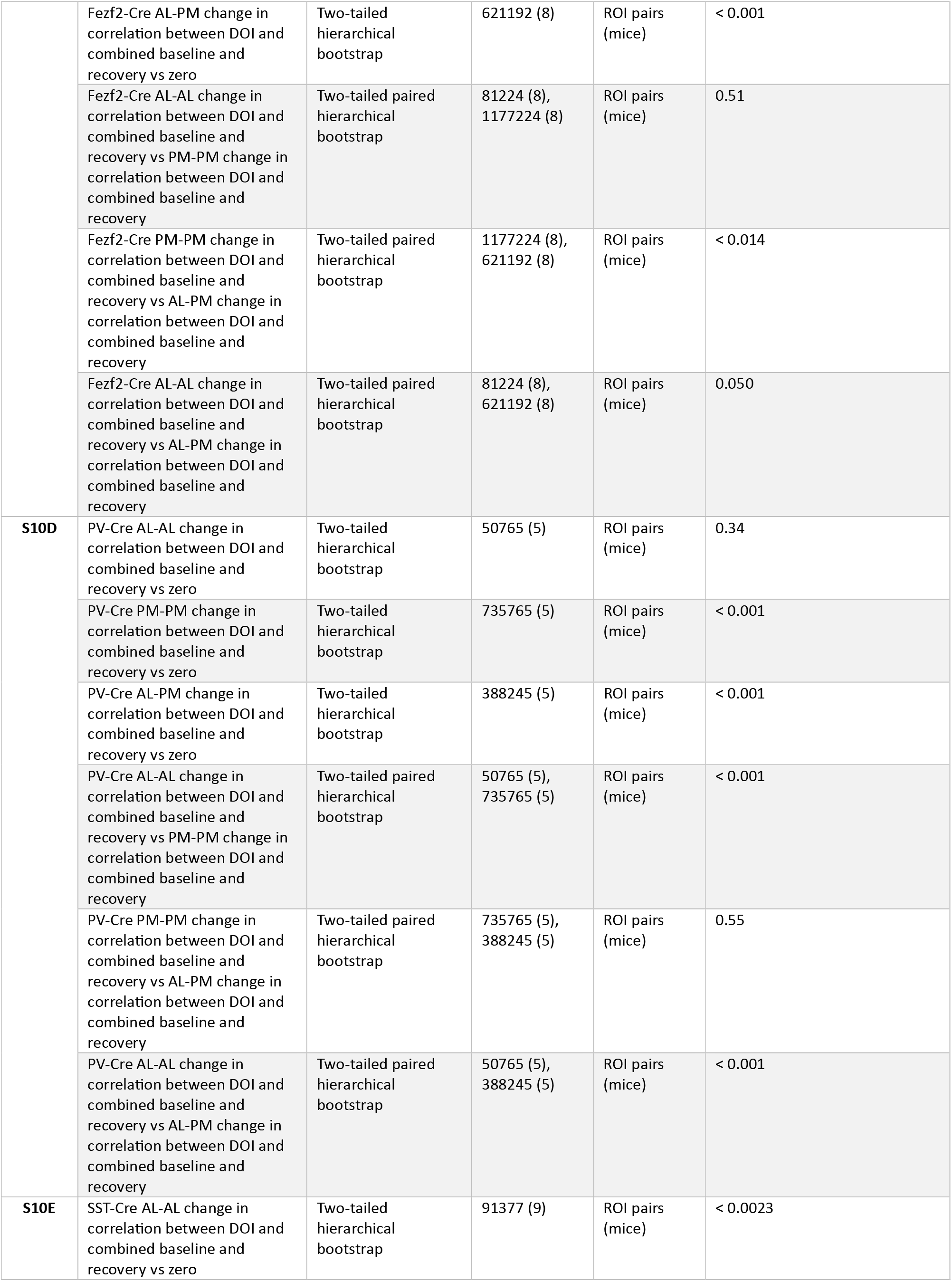
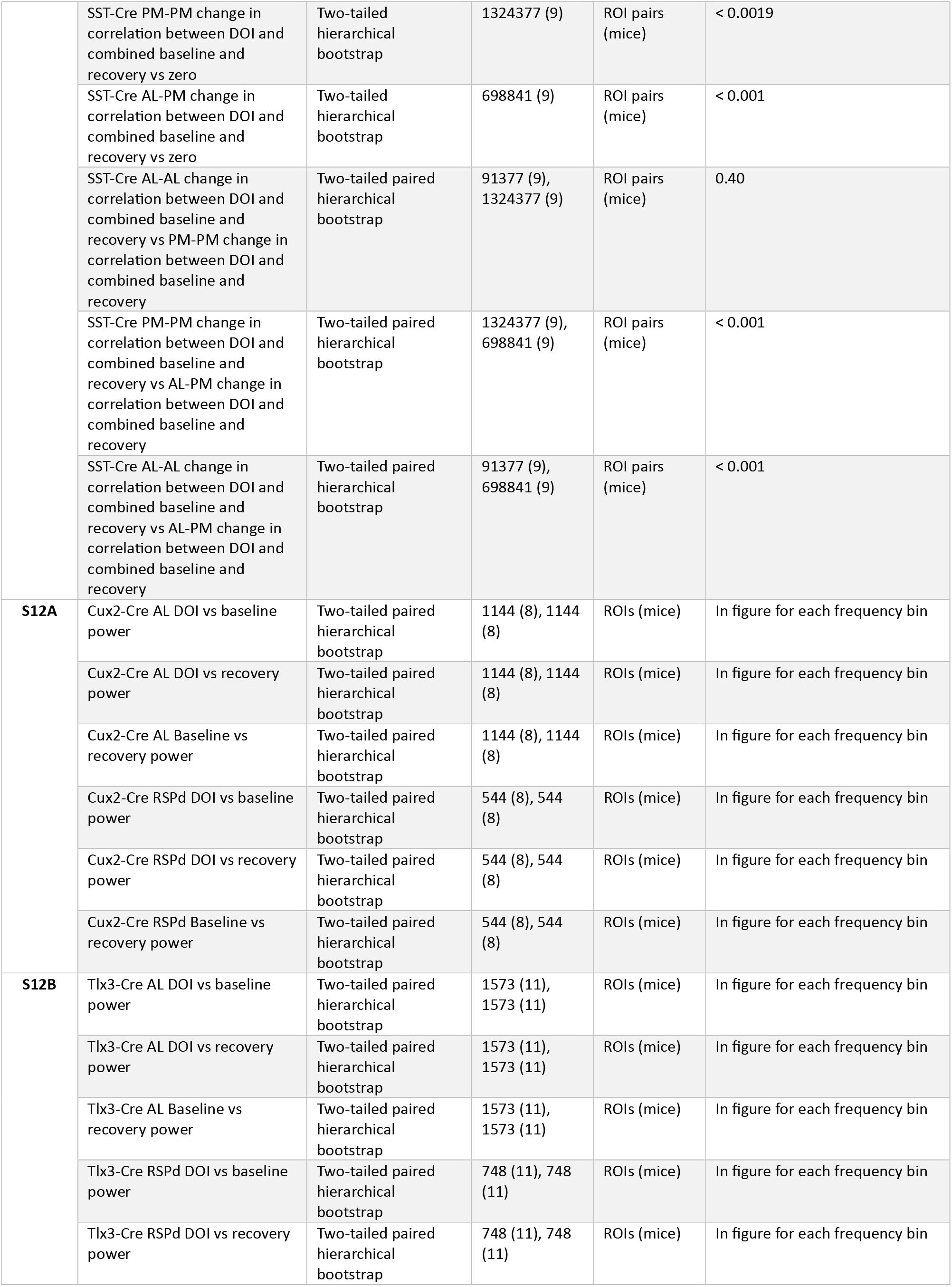
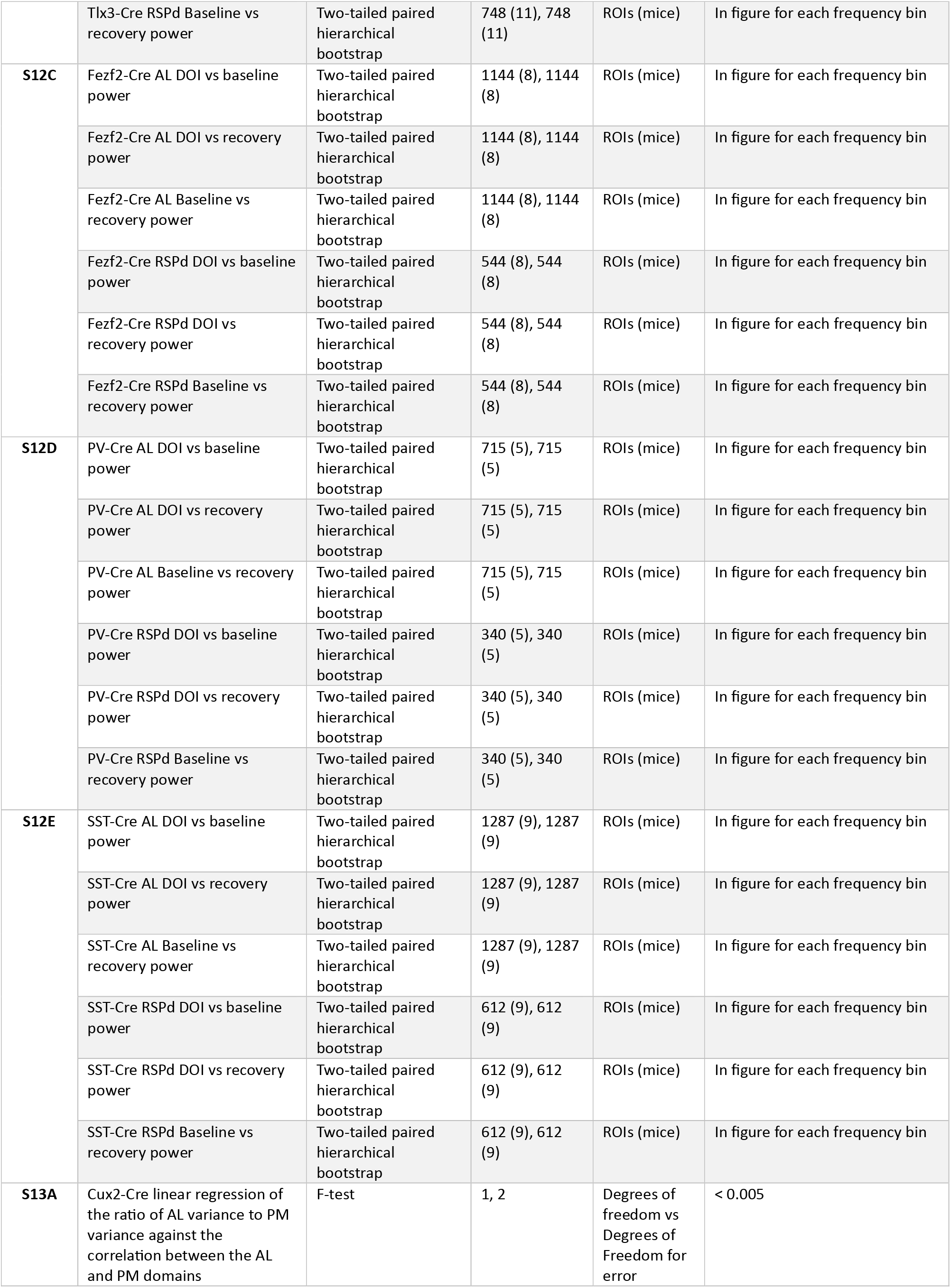
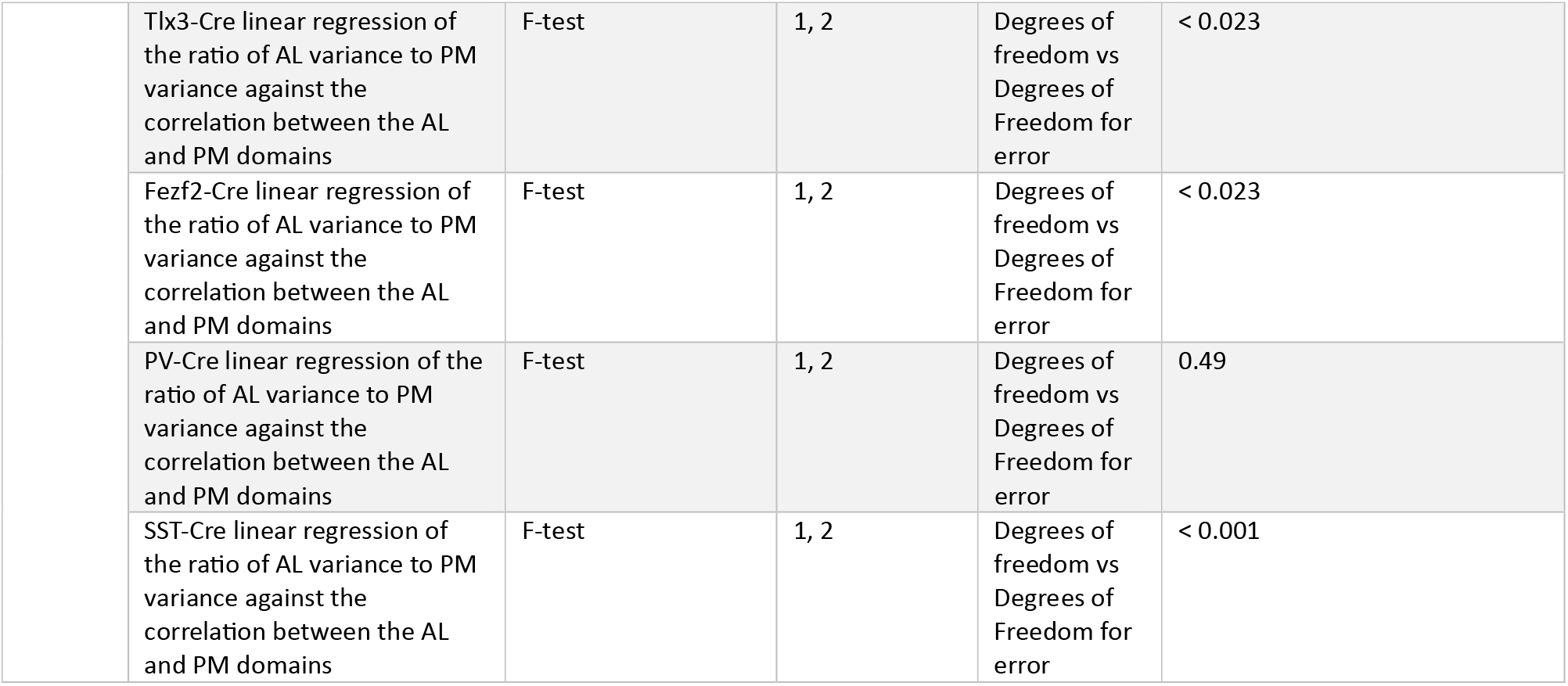
Statistical information. For a description of all the statistical tests used, please refer to the **Methods** section, paragraph **Statistical Analysis**.

| Figure | Comparison | Test | N<br>(per group) | Unit | p-value |
| --- | --- | --- | --- | --- | --- |
| **1H** | Fidget response in the AL domain vs in the PM domain | Two-tailed paired hierarchical bootstrap | 1573 (11),<br>5973 (11) | ROIs (mice) | In figure for each time bin |
| **2B** | DOI HTR rate vs baseline HTR rate | Two-tailed paired bootstrap | 11, 11 | Mice | In figure for each time bin |
|  | DOI HTR rate vs recovery HTR rate | Two-tailed paired bootstrap | 11, 11 | Mice | In figure for each time bin |
|  | Baseline HTR rate vs recovery HTR rate | Two-tailed paired bootstrap | 11, 11 | Mice | In figure for each time bin |
| **2E** | AL-AL change in correlation between DOI and combined baseline and recovery vs zero | Two-tailed hierarchical bootstrap | 111683 (11) | ROI pairs (mice) | 0.66 |
|  | PM-PM change in correlation between DOI and combined baseline and recovery vs zero | Two-tailed hierarchical bootstrap | 1618683 (11) | ROI pairs (mice) | < 0.001 |
|  | AL-PM change in correlation between DOI and combined baseline and recovery vs zero | Two-tailed hierarchical bootstrap | 854139 (11) | ROI pairs (mice) | < 0.001 |
|  | AL-AL change in correlation between DOI and combined baseline and recovery vs PM-PM change in correlation between DOI and combined baseline and recovery | Two-tailed paired hierarchical bootstrap | 111683 (11),<br>1618683 (11) | ROI pairs (mice) | 0.18 |
|  | PM-PM change in correlation between DOI and combined baseline and recovery vs AL-PM change in correlation between DOI and combined baseline and recovery | Two-tailed paired hierarchical bootstrap | 1618683 (11),<br>854139 (11) | ROI pairs (mice) | < 0.001 |
|  | AL-AL change in correlation between DOI and combined baseline and recovery vs AL-PM change in correlation between DOI and combined baseline and recovery | Two-tailed paired hierarchical bootstrap | 111683 (11),<br>854139 (11) | ROI pairs (mice) | < 0.001 |
| **3B** | Ketanserin+DOI HTR rate vs baseline HTR rate | Two-tailed paired bootstrap | 5, 5 | Mice | In figure for each time bin |
|  | Ketanserin+DOI HTR rate vs recovery HTR rate | Two-tailed paired bootstrap | 5, 5 | Mice | In figure for each time bin |
|  | Baseline HTR rate vs recovery HTR rate | Two-tailed paired bootstrap | 5, 5 | Mice | In figure for each time bin |
| **3E** | AL-AL change in correlation between DOI and combined baseline and recovery vs zero | Two-tailed hierarchical bootstrap | 50765 (5) | ROI pairs (mice) | < 0.001 |
|  | PM-PM change in correlation between DOI and combined baseline and recovery vs zero | Two-tailed hierarchical bootstrap | 735765 (5) | ROI pairs (mice) | < 0.001 |
|  | AL-PM change in correlation between DOI and combined baseline and recovery vs zero | Two-tailed hierarchical bootstrap | 388245 (5) | ROI pairs (mice) | < 0.001 |
|  | AL-AL change in correlation between DOI and combined baseline and recovery vs PM-PM change in correlation between DOI and combined baseline and recovery | Two-tailed paired hierarchical bootstrap | 50765 (5),<br>735765 (5) | ROI pairs (mice) | < 0.001 |
|  | PM-PM change in correlation between DOI and combined baseline and recovery vs AL-PM change in correlation between DOI and combined baseline and recovery | Two-tailed paired hierarchical bootstrap | 735765 (5), 388245 (5) | ROI pairs (mice) | 0.18 |
|  | AL-AL change in correlation between DOI and combined baseline and recovery vs AL-PM change in correlation between DOI and combined baseline and recovery | Two-tailed paired hierarchical bootstrap | 50765 (5), 388245 (5) | ROI pairs (mice) | < 0.001 |
| **4B** | AL change in variance between DOI and combined baseline and recovery vs zero | Two-tailed hierarchical bootstrap | 1573 (11) | ROIs (mice) | < 0.001 |
|  | RSPd change in variance between DOI and combined baseline and recovery vs zero | Two-tailed hierarchical bootstrap | 748 (11) | ROIs (mice) | < 0.001 |
|  | PM without RSPd change in variance between DOI and combined baseline and recovery vs zero | Two-tailed hierarchical bootstrap | 5225 (11) | ROIs (mice) | < 0.001 |
|  | AL change in variance between DOI and combined baseline vs RSPd change in variance between DOI and combined baseline and recovery | Two-tailed paired hierarchical bootstrap | 1573 (11), 748 (11) | ROIs (mice) | < 0.001 |
|  | RSPd change in variance between DOI and combined baseline vs PM without RSPd change in variance between DOI and combined baseline and recovery | Two-tailed paired hierarchical bootstrap | 748 (11), 5225 (11) | ROIs (mice) | < 0.001 |
|  | AL change in variance between DOI and combined baseline vs PM without RSPd change in variance between DOI and combined baseline and recovery | Two-tailed paired hierarchical bootstrap | 1573 (11), 5225 (11) | ROIs (mice) | < 0.001 |
| **4C** | DOI vs baseline response in the AL domain | Two-tailed paired hierarchical bootstrap | 1573 (11), 1573 (11) | ROIs (mice) | In figure for each time bin |
|  | DOI vs recovery response in the AL domain | Two-tailed paired hierarchical bootstrap | 1573 (11), 1573 (11) | ROIs (mice) | In figure for each time bin |
|  | DOI vs ketanserin+DOI response in the AL domain | Two-tailed hierarchical bootstrap | 1573 (11), 715 (5) | ROIs (mice) | In figure for each time bin |
|  | Baseline vs recovery response in the AL domain | Two-tailed paired hierarchical bootstrap | 1573 (11), 1573 (11) | ROIs (mice) | In figure for each time bin |
|  | Baseline vs ketanserin+DOI response in the AL domain | Two-tailed hierarchical bootstrap | 1573 (11), 715 (5) | ROIs (mice) | In figure for each time bin |
|  | Recovery vs ketanserin+DOI response in the AL domain | Two-tailed hierarchical bootstrap | 1573 (11), 715 (5) | ROIs (mice) | In figure for each time bin |
| **4D** | DOI vs baseline response in the PM domain | Two-tailed paired hierarchical bootstrap | 5973 (11), 5973 (11) | ROIs (mice) | In figure for each time bin |
|  | DOI vs recovery response in the PM domain | Two-tailed paired hierarchical bootstrap | 5973 (11), 5973 (11) | ROIs (mice) | In figure for each time bin |
|  | DOI vs ketanserin+DOI response in the PM domain | Two-tailed hierarchical bootstrap | 5973 (11), 2715 (5) | ROIs (mice) | In figure for each time bin |
|  | Baseline vs recovery response in the PM domain | Two-tailed paired hierarchical bootstrap | 5973 (11), 5973 (11) | ROIs (mice) | In figure for each time bin |
|  | Baseline vs ketanserin+DOI response in the PM domain | Two-tailed hierarchical bootstrap | 5973 (11), 2715 (5) | ROIs (mice) | In figure for each time bin |
|  | Recovery vs ketanserin+DOI response in the PM domain | Two-tailed hierarchical bootstrap | 5973 (11), 2715 (5) | ROIs (mice) | In figure for each time bin |
| **5A** | DOI vs baseline power in the AL domain | Two-tailed paired hierarchical bootstrap | 1573 (11), 1573 (11) | ROIs (mice) | In figure for each frequency bin |
|  | DOI vs recovery power in the AL domain | Two-tailed paired hierarchical bootstrap | 1573 (11), 1573 (11) | ROIs (mice) | In figure for each frequency bin |
|  | DOI vs ketanserin+DOI power in the AL domain | Two-tailed hierarchical bootstrap | 1573 (11), 715 (5) | ROIs (mice) | In figure for each frequency bin |
|  | Baseline vs recovery power in the AL domain | Two-tailed paired hierarchical bootstrap | 1573 (11), 1573 (11) | ROIs (mice) | In figure for each frequency bin |
|  | Baseline vs ketanserin+DOI power in the AL domain | Two-tailed hierarchical bootstrap | 1573 (11), 715 (5) | ROIs (mice) | In figure for each frequency bin |
|  | Recovery vs ketanserin+DOI power in the AL domain | Two-tailed hierarchical bootstrap | 1573 (11), 715 (5) | ROIs (mice) | In figure for each frequency bin |
| **5B** | DOI vs baseline power in RSPd | Two-tailed paired hierarchical bootstrap | 748 (11), 748 (11) | ROIs (mice) | In figure for each frequency bin |
|  | DOI vs recovery power in RSPd | Two-tailed paired hierarchical bootstrap | 748 (11), 748 (11) | ROIs (mice) | In figure for each frequency bin |
|  | DOI vs ketanserin+DOI power in RSPd | Two-tailed hierarchical bootstrap | 748 (11), 340 (5) | ROIs (mice) | In figure for each frequency bin |
|  | Baseline vs recovery power in RSPd | Two-tailed paired hierarchical bootstrap | 748 (11), 748 (11) | ROIs (mice) | In figure for each frequency bin |
|  | Baseline vs ketanserin+DOI power in RSPd | Two-tailed hierarchical bootstrap | 748 (11), 340 (5) | ROIs (mice) | In figure for each frequency bin |
|  | Recovery vs ketanserin+DOI power in RSPd | Two-tailed hierarchical bootstrap | 748 (11), 340 (5) | ROIs (mice) | In figure for each frequency bin |
| **5D** | DOI vs baseline response for AL electrodes | Two-tailed paired hierarchical bootstrap | 50 (5) | Electrodes (mice) | In figure for each frequency bin |
|  | DOI vs recovery response for AL electrodes | Two-tailed paired hierarchical bootstrap | 50 (5) | Electrodes (mice) | In figure for each frequency bin |
|  | Baseline vs recovery response for AL electrodes | Two-tailed paired hierarchical bootstrap | 50 (5) | Electrodes (mice) | In figure for each frequency bin |
| **5E** | DOI vs baseline response for RSPd electrodes | Two-tailed paired hierarchical bootstrap | 20 (5) | Electrodes (mice) | In figure for each frequency bin |
|  | DOI vs recovery response for RSPd electrodes | Two-tailed paired hierarchical bootstrap | 20 (5) | Electrodes (mice) | In figure for each frequency bin |
|  | Baseline vs recovery response for RSPd electrodes | Two-tailed paired hierarchical bootstrap | 20 (5) | Electrodes (mice) | In figure for each frequency bin |
| **6D** | DOI vs baseline response in the PM domain | Two-tailed paired hierarchical bootstrap | 4887 (9), 4887 (9) | ROIs (mice) | In figure for each time bin |
|  | DOI vs recovery response in the PM domain | Two-tailed paired hierarchical bootstrap | 4887 (9), 4887 (9) | ROIs (mice) | In figure for each time bin |
|  | Baseline vs recovery response in the PM domain | Two-tailed paired hierarchical bootstrap | 4887 (9), 4887 (9) | ROIs (mice) | In figure for each time bin |
| **6E** | DOI vs baseline response in the AL domain | Two-tailed paired hierarchical bootstrap | 1287 (9), 1287 (9) | ROIs (mice) | In figure for each time bin |
|  | DOI vs recovery response in the AL domain | Two-tailed paired hierarchical bootstrap | 1287 (9), 1287 (9) | ROIs (mice) | In figure for each time bin |
|  | Baseline vs recovery response in the AL domain | Two-tailed paired hierarchical bootstrap | 1287 (9), 1287 (9) | ROIs (mice) | In figure for each time bin |
| **6G** | DOI vs baseline response in the AL domain | Two-tailed paired hierarchical bootstrap | 1287 (9), 1287 (9) | ROIs (mice) | In figure for each time bin |
|  | DOI vs recovery response in the AL domain | Two-tailed paired hierarchical bootstrap | 1287 (9), 1287 (9) | ROIs (mice) | In figure for each time bin |
|  | Baseline vs recovery response in the AL domain | Two-tailed paired hierarchical bootstrap | 1287 (9), 1287 (9) | ROIs (mice) | In figure for each time bin |
| **6H** | DOI vs baseline response in the PM domain | Two-tailed paired hierarchical bootstrap | 4887 (9), 4887 (9) | ROIs (mice) | In figure for each time bin |
|  | DOI vs recovery response in the PM domain | Two-tailed paired hierarchical bootstrap | 4887 (9), 4887 (9) | ROIs (mice) | In figure for each time bin |
|  | Baseline vs recovery response in the PM domain | Two-tailed paired hierarchical bootstrap | 4887 (9), 4887 (9) | ROIs (mice) | In figure for each time bin |
| **7D** | Center: DOI vs baseline response | Two-tailed paired hierarchical bootstrap | 539 (11), 539 (11) | ROIs (mice) | In figure for each time bin |
|  | Center: DOI vs recovery response | Two-tailed paired hierarchical bootstrap | 539 (11), 539 (11) | ROIs (mice) | In figure for each time bin |
|  | Center: Baseline vs recovery response | Two-tailed paired hierarchical bootstrap | 539 (11), 539 (11) | ROIs (mice) | In figure for each time bin |
|  | Surround: DOI vs baseline response | Two-tailed paired hierarchical bootstrap | 2986 (11), 2986 (11) | ROIs (mice) | In figure for each time bin |
|  | Surround: DOI vs recovery response | Two-tailed paired hierarchical bootstrap | 2986 (11), 2986 (11) | ROIs (mice) | In figure for each time bin |
|  | Surround: Baseline vs recovery response | Two-tailed paired hierarchical bootstrap | 2986 (11), 2986 (11) | ROIs (mice) | In figure for each time bin |
| **7F** | Activity in PM: DOI vs baseline response | Two-tailed paired hierarchical bootstrap | 5973 (11), 5973 (11) | ROIs (mice) | In figure for each time bin |
|  | Activity in PM: DOI vs recovery response | Two-tailed paired hierarchical bootstrap | 5973 (11), 5973 (11) | ROIs (mice) | In figure for each time bin |
|  | Activity in PM: Baseline vs recovery response | Two-tailed paired hierarchical bootstrap | 5973 (11), 5973 (11) | ROIs (mice) | In figure for each time bin |
|  | Activity in AL: DOI vs baseline response | Two-tailed paired hierarchical bootstrap | 1573 (11), 1573 (11) | ROIs (mice) | In figure for each time bin |
|  | Activity in AL: DOI vs recovery response | Two-tailed paired hierarchical bootstrap | 1573 (11), 1573 (11) | ROIs (mice) | In figure for each time bin |
|  | Activity in AL: Baseline vs recovery response | Two-tailed paired hierarchical bootstrap | 1573 (11), 1573 (11) | ROIs (mice) | In figure for each time bin |
| **7G** | Activity in AL: DOI vs baseline response | Two-tailed paired hierarchical bootstrap | 1573 (11), 1573 (11) | ROIs (mice) | In figure for each time bin |
|  | Activity in AL: DOI vs recovery response | Two-tailed paired hierarchical bootstrap | 1573 (11), 1573 (11) | ROIs (mice) | In figure for each time bin |
|  | Activity in AL: Baseline vs recovery response | Two-tailed paired hierarchical bootstrap | 1573 (11), 1573 (11) | ROIs (mice) | In figure for each time bin |
|  | Activity in PM: DOI vs baseline response | Two-tailed paired hierarchical bootstrap | 5973 (11), 5973 (11) | ROIs (mice) | In figure for each time bin |
|  | Activity in PM: DOI vs recovery response | Two-tailed paired hierarchical bootstrap | 5973 (11), 5973 (11) | ROIs (mice) | In figure for each time bin |
|  | Activity in PM: Baseline vs recovery response | Two-tailed paired hierarchical bootstrap | 5973 (11), 5973 (11) | ROIs (mice) | In figure for each time bin |
| **8B** | Cux2-Cre AL change in variance between DOI and combined baseline and recovery vs zero | Two-tailed hierarchical bootstrap | 1144 (8) | ROIs (mice) | < 0.001 |
|  | Cux2-Cre RSPd change in variance between DOI and combined baseline and recovery vs zero | Two-tailed hierarchical bootstrap | 544 (8) | ROIs (mice) | < 0.001 |
|  | Cux2-Cre PM without RSPd change in variance between DOI and combined baseline and recovery vs zero | Two-tailed hierarchical bootstrap | 3800 (8) | ROIs (mice) | < 0.0017 |
|  | Cux2-Cre AL change in variance between DOI and combined baseline vs RSPd change in variance between DOI and combined baseline and recovery | Two-tailed paired hierarchical bootstrap | 1144 (8), 544 (8) | ROIs (mice) | < 0.001 |
|  | Cux2-Cre RSPd change in variance between DOI and combined baseline vs PM without RSPd change in variance between DOI and combined baseline and recovery | Two-tailed paired hierarchical bootstrap | 544 (8), 3800 (8) | ROIs (mice) | < 0.0025 |
|  | Cux2-Cre AL change in variance between DOI and combined baseline vs PM without RSPd change in variance between DOI and combined baseline and recovery | Two-tailed paired hierarchical bootstrap | 1144 (8), 3800 (8) | ROIs (mice) | < 0.001 |
|  | Tlx3-Cre AL change in variance between DOI and combined baseline and recovery vs zero | Two-tailed hierarchical bootstrap | 1573 (11) | ROIs (mice) | < 0.001 |
|  | Tlx3-Cre RSPd change in variance between DOI and combined baseline and recovery vs zero | Two-tailed hierarchical bootstrap | 748 (11) | ROIs (mice) | < 0.0053 |
|  | Tlx3-Cre PM without RSPd change in variance between DOI and combined baseline and recovery vs zero | Two-tailed hierarchical bootstrap | 5225 (11) | ROIs (mice) | 0.11 |
|  | Tlx3-Cre AL change in variance between DOI and combined baseline vs RSPd change in variance between DOI and combined baseline and recovery | Two-tailed paired hierarchical bootstrap | 1573 (11), 748 (11) | ROIs (mice) | < 0.001 |
|  | Tlx3-Cre RSPd change in variance between DOI and combined baseline vs PM without RSPd change in variance between DOI and combined baseline and recovery | Two-tailed paired hierarchical bootstrap | 748 (11), 5225 (11) | ROIs (mice) | 0.37 |
|  | Tlx3-Cre AL change in variance between DOI and combined baseline vs PM without RSPd change in variance between DOI and combined baseline and recovery | Two-tailed paired hierarchical bootstrap | 1573 (11), 5225 (11) | ROIs (mice) | < 0.001 |
|  | Fezf2-Cre AL change in variance between DOI and combined baseline and recovery vs zero | Two-tailed hierarchical bootstrap | 1144 (8) | ROIs (mice) | < 0.021 |
|  | Fezf2-Cre RSPd change in variance between DOI and combined baseline and recovery vs zero | Two-tailed hierarchical bootstrap | 544 (8) | ROIs (mice) | < 0.001 |
|  | Fezf2-Cre PM without RSPd change in variance between DOI and combined baseline and recovery vs zero | Two-tailed hierarchical bootstrap | 3800 (8) | ROIs (mice) | < 0.001 |
|  | Fezf2-Cre AL change in variance between DOI and combined baseline vs RSPd change in variance between DOI and combined baseline and recovery | Two-tailed paired hierarchical bootstrap | 1144 (8), 544 (8) | ROIs (mice) | < 0.001 |
|  | Fezf2-Cre RSPd change in variance between DOI and combined baseline vs PM without RSPd change in variance between DOI and combined baseline and recovery | Two-tailed paired hierarchical bootstrap | 544 (8), 3800 (8) | ROIs (mice) | < 0.001 |
|  | Fezf2-Cre AL change in variance between DOI and combined baseline vs PM without RSPd change in variance between DOI and combined baseline and recovery | Two-tailed paired hierarchical bootstrap | 1144 (8), 3800 (8) | ROIs (mice) | < 0.001 |
|  | PV-Cre AL change in variance between DOI and combined baseline and recovery vs zero | Two-tailed hierarchical bootstrap | 715 (5) | ROIs (mice) | < 0.034 |
|  | PV-Cre RSPd change in variance between DOI and combined baseline and recovery vs zero | Two-tailed hierarchical bootstrap | 340 (5) | ROIs (mice) | < 0.001 |
|  | PV-Cre PM without RSPd change in variance between DOI and combined baseline and recovery vs zero | Two-tailed hierarchical bootstrap | 2375 (5) | ROIs (mice) | < 0.001 |
|  | PV-Cre AL change in variance between DOI and combined baseline vs RSPd change in variance between DOI and combined baseline and recovery | Two-tailed paired hierarchical bootstrap | 715 (5), 340 (5) | ROIs (mice) | < 0.001 |
|  | PV-Cre RSPd change in variance between DOI and combined baseline vs PM without RSPd change in variance between DOI and combined baseline and recovery | Two-tailed paired hierarchical bootstrap | 340 (5), 2375 (5) | ROIs (mice) | < 0.001 |
|  | PV-Cre AL change in variance between DOI and combined baseline vs PM without RSPd change in variance between DOI and combined baseline and recovery | Two-tailed paired hierarchical bootstrap | 715 (5), 2375 (5) | ROIs (mice) | < 0.001 |
|  | SST-Cre AL change in variance between DOI and combined baseline and recovery vs zero | Two-tailed hierarchical bootstrap | 1287 (9) | ROIs (mice) | 0.28 |
|  | SST-Cre RSPd change in variance between DOI and combined baseline and recovery vs zero | Two-tailed hierarchical bootstrap | 612 (9) | ROIs (mice) | 0.21 |
|  | SST-Cre PM without RSPd change in variance between DOI and combined baseline and recovery vs zero | Two-tailed hierarchical bootstrap | 4275 (9) | ROIs (mice) | < 0.001 |
|  | SST-Cre AL change in variance between DOI and combined baseline vs RSPd change in variance between DOI and combined baseline and recovery | Two-tailed paired hierarchical bootstrap | 1287 (9), 612 (9) | ROIs (mice) | < 0.0031 |
|  | SST-Cre RSPd change in variance between DOI and combined baseline vs PM without RSPd change in variance between DOI and combined baseline and recovery | Two-tailed paired hierarchical bootstrap | 612 (9), 4275 (9) | ROIs (mice) | 0.29 |
|  | SST-Cre AL change in variance between DOI and combined baseline vs PM without RSPd change in variance between DOI and combined baseline and recovery | Two-tailed paired hierarchical bootstrap | 1287 (9), 4275 (9) | ROIs (mice) | < 0.001 |
| **S3B** | AL-AL change in correlation between DOI and combined baseline and recovery vs zero | Two-tailed hierarchical bootstrap | 60918 (6) | ROI pairs (mice) | 0.22 |
|  | PM-PM change in correlation between DOI and combined baseline and recovery vs zero | Two-tailed hierarchical bootstrap | 882918 (6) | ROI pairs (mice) | 1.00 |
|  | AL-PM change in correlation between DOI and combined baseline and recovery vs zero | Two-tailed hierarchical bootstrap | 465894 (6) | ROI pairs (mice) | 0.54 |
|  | AL-AL change in correlation between DOI and combined baseline and recovery vs PM-PM change in correlation between DOI and combined baseline and recovery | Two-tailed paired hierarchical bootstrap | 60918 (6), 882918 (6) | ROI pairs (mice) | 0.09 |
|  | PM-PM change in correlation between DOI and combined baseline and recovery vs AL-PM change in correlation between DOI and combined baseline and recovery | Two-tailed paired hierarchical bootstrap | 882918 (6), 465894 (6) | ROI pairs (mice) | 0.23 |
|  | AL-AL change in correlation between DOI and combined baseline and recovery vs AL-PM change in correlation between DOI and combined baseline and recovery | Two-tailed paired hierarchical bootstrap | 60918 (6), 465894 (6) | ROI pairs (mice) | < 0.047 |
| **S4A** | AL-AL change in correlation between combined baseline and recovery with a common occluder and DOI vs zero | Two-tailed hierarchical bootstrap | 111683 (11) | ROI pairs (mice) | < 0.0011 |
|  | PM-PM change in correlation between combined baseline and recovery with a common occluder and DOI vs zero | Two-tailed hierarchical bootstrap | 1618683 (11) | ROI pairs (mice) | < 0.001 |
|  | AL-PM change in correlation between combined baseline and recovery with a common occluder and DOI vs zero | Two-tailed hierarchical bootstrap | 854139 (11) | ROI pairs (mice) | 0.92 |
|  | AL-AL change in correlation between combined baseline and recovery with a common occluder and DOI vs PM-PM change in correlation between combined baseline and recovery with a common occluder and DOI | Two-tailed paired hierarchical bootstrap | 111683 (11), 1618683 (11) | ROI pairs (mice) | 0.13 |
|  | PM-PM change in correlation between combined baseline and recovery with a common occluder and DOI vs AL-PM change in correlation between combined baseline and recovery with a common occluder and DOI | Two-tailed paired hierarchical bootstrap | 1618683 (11), 854139 (11) | ROI pairs (mice) | < 0.001 |
|  | AL-AL change in correlation between combined baseline and recovery with a common occluder and DOI vs AL-PM change in correlation between combined baseline and recovery with a common occluder and DOI | Two-tailed paired hierarchical bootstrap | 111683 (11), 854139 (11) | ROI pairs (mice) | < 0.001 |
| **S4B** | AL-AL change in correlation between combined baseline and recovery with a common occluder and DOI vs zero | Two-tailed hierarchical bootstrap | 111683 (11) | ROI pairs (mice) | 0.58 |
|  | PM-PM change in correlation between combined baseline and recovery with a common occluder and DOI vs zero | Two-tailed hierarchical bootstrap | 1618683 (11) | ROI pairs (mice) | 0.2 |
|  | AL-PM change in correlation combined baseline and recovery with a common occluder and DOI vs zero | Two-tailed hierarchical bootstrap | 854139 (11) | ROI pairs (mice) | < 0.001 |
|  | AL-AL change in correlation between combined baseline and recovery with a common occluder and DOI vs PM-PM change in correlation between combined baseline and recovery with a common occluder and DOI | Two-tailed paired hierarchical bootstrap | 111683 (11), 1618683 (11) | ROI pairs (mice) | 0.74 |
|  | PM-PM change in correlation between combined baseline and recovery with a common occluder and DOI vs AL-PM change in correlation between combined baseline and recovery with a common occluder and DOI | Two-tailed paired hierarchical bootstrap | 1618683 (11), 854139 (11) | ROI pairs (mice) | < 0.001 |
|  | AL-AL change in correlation between combined baseline and recovery with a common occluder and DOI vs AL-PM change in correlation between combined baseline and recovery with a common occluder and DOI | Two-tailed paired hierarchical bootstrap | 111683 (11), 854139 (11) | ROI pairs (mice) | < 0.001 |
| **S5B** | EGFP AL change in variance between DOI and combined baseline and recovery vs zero | Two-tailed hierarchical bootstrap | 858 (6) | ROIs (mice) | < 0.0035 |
|  | EGFP RSPd change in variance between DOI and combined baseline and recovery vs zero | Two-tailed hierarchical bootstrap | 408 (6) | ROIs (mice) | 0.42 |
|  | EGFP PM without RSPd change in variance between DOI and combined baseline and recovery vs zero | Two-tailed hierarchical bootstrap | 2850 (6) | ROIs (mice) | 0.25 |
|  | GCaMP AL change in variance between DOI and combined baseline and recovery vs zero | Two-tailed hierarchical bootstrap | 1573 (11) | ROIs (mice) | < 0.001 |
|  | GCaMP RSPd change in variance between DOI and combined baseline and recovery vs zero | Two-tailed hierarchical bootstrap | 748 (11) | ROIs (mice) | < 0.001 |
|  | GCaMP PM without RSPd change in variance between DOI and combined baseline and recovery vs zero | Two-tailed hierarchical bootstrap | 5225 (11) | ROIs (mice) | < 0.001 |
|  | EGFP AL change in variance between DOI and combined baseline and recovery vs GCaMP AL change in variance between DOI and combined baseline and recovery | Two-tailed hierarchical bootstrap | 858 (6), 1573 (11) | ROIs (mice) | < 0.001 |
|  | EGFP RSPd change in variance between DOI and combined baseline and recovery vs GCaMP RSPd change in variance between DOI and combined baseline and recovery | Two-tailed hierarchical bootstrap | 408 (6), 748 (11) | ROIs (mice) | < 0.001 |
|  | EGFP PM without RSPd change in variance between DOI and combined baseline and recovery vs GCaMP PM without RSPd change in variance between DOI and combined baseline and recovery | Two-tailed hierarchical bootstrap | 2850 (6), 5225 (11) | ROIs (mice) | < 0.001 |
| **S6A** | Linear regression of the ratio of AL variance to PM variance against the correlation between the AL and PM domains | F-test | 1, 2 | Degrees of freedom vs Degrees of Freedom for error | < 0.018 |
| **S6B** | DOI vs baseline AL-PM correlation | Two-tailed paired hierarchical bootstrap | 854139 (11), 854139 (11) | ROI pairs (mice) | In figure for each frequency range bin |
|  | DOI vs recovery AL-PM correlation | Two-tailed paired hierarchical bootstrap | 854139 (11), 854139 (11) | ROI pairs (mice) | In figure for each frequency range bin |
|  | Baseline vs recovery AL-PM correlation | Two-tailed paired hierarchical bootstrap | 854139 (11), 854139 (11) | ROI pairs (mice) | In figure for each frequency range bin |
| **S6C** | DOI vs baseline AL-PM correlation | Two-tailed paired hierarchical bootstrap | 465894 (6), 465894 (6) | ROI pairs (mice) | In figure for each frequency range bin |
|  | DOI vs recovery AL-PM correlation | Two-tailed paired hierarchical bootstrap | 465894 (6), 465894 (6) | ROI pairs (mice) | In figure for each frequency range bin |
|  | Baseline vs recovery AL-PM correlation | Two-tailed paired hierarchical bootstrap | 465894 (6), 465894 (6) | ROI pairs (mice) | In figure for each frequency range bin |
| **S7A** | DOI vs baseline power in the AL domain | Two-tailed paired hierarchical bootstrap | 858 (6), 858 (6) | ROIs (mice) | In figure for each frequency bin |
|  | DOI vs recovery power in the AL domain | Two-tailed paired hierarchical bootstrap | 858 (6), 858 (6) | ROIs (mice) | In figure for each frequency bin |
|  | Baseline vs recovery power in the AL domain | Two-tailed paired hierarchical bootstrap | 858 (6), 858 (6) | ROIs (mice) | In figure for each frequency bin |
| **S7B** | DOI vs baseline power in RSPd | Two-tailed paired hierarchical bootstrap | 408 (6), 408 (6) | ROIs (mice) | In figure for each frequency bin |
|  | DOI vs recovery power in RSPd | Two-tailed paired hierarchical bootstrap | 408 (6), 408 (6) | ROIs (mice) | In figure for each frequency bin |
|  | Baseline vs recovery power in RSPd | Two-tailed paired hierarchical bootstrap | 408 (6), 408 (6) | ROIs (mice) | In figure for each frequency bin |
| **S10A** | Cux2-Cre AL-AL change in correlation between DOI and combined baseline and recovery vs zero | Two-tailed hierarchical bootstrap | 81224 (8) | ROI pairs (mice) | 0.24 |
|  | Cux2-Cre PM-PM change in correlation between DOI and combined baseline and recovery vs zero | Two-tailed hierarchical bootstrap | 1177224 (8) | ROI pairs (mice) | 0.32 |
|  | Cux2-Cre AL-PM change in correlation between DOI and combined baseline and recovery vs zero | Two-tailed hierarchical bootstrap | 621192 (8) | ROI pairs (mice) | < 0.0095 |
|  | Cux2-Cre AL-AL change in correlation between DOI and combined baseline and recovery vs PM-PM change in correlation between DOI and combined baseline and recovery | Two-tailed paired hierarchical bootstrap | 81224 (8), 1177224 (8) | ROI pairs (mice) | 0.90 |
|  | Cux2-Cre PM-PM change in correlation between DOI and combined baseline and recovery vs AL-PM change in correlation between DOI and combined baseline and recovery | Two-tailed paired hierarchical bootstrap | 1177224 (8), 621192 (8) | ROI pairs (mice) | < 0.001 |
|  | Cux2-Cre AL-AL change in correlation between DOI and combined baseline and recovery vs AL-PM change in correlation between DOI and combined baseline and recovery | Two-tailed paired hierarchical bootstrap | 81224 (8), 621192 (8) | ROI pairs (mice) | < 0.001 |
| **S10B** | Tlx3-Cre AL-AL change in correlation between DOI and combined baseline and recovery vs zero | Two-tailed hierarchical bootstrap | 111683 (11) | ROI pairs (mice) | < 0.001 |
|  | Tlx3-Cre PM-PM change in correlation between DOI and combined baseline and recovery vs zero | Two-tailed hierarchical bootstrap | 1618683 (11) | ROI pairs (mice) | < 0.001 |
|  | Tlx3-Cre AL-PM change in correlation between DOI and combined baseline and recovery vs zero | Two-tailed hierarchical bootstrap | 854139 (11) | ROI pairs (mice) | < 0.001 |
|  | Tlx3-Cre AL-AL change in correlation between DOI and combined baseline and recovery vs PM-PM change in correlation between DOI and combined baseline and recovery | Two-tailed paired hierarchical bootstrap | 111683 (11), 1618683 (11) | ROI pairs (mice) | 0.10 |
|  | Tlx3-Cre PM-PM change in correlation between DOI and combined baseline and recovery vs AL-PM change in correlation between DOI and combined baseline and recovery | Two-tailed paired hierarchical bootstrap | 1618683 (11), 854139 (11) | ROI pairs (mice) | < 0.001 |
|  | Tlx3-Cre AL-AL change in correlation between DOI and combined baseline and recovery vs AL-PM change in correlation between DOI and combined baseline and recovery | Two-tailed paired hierarchical bootstrap | 111683 (11), 854139 (11) | ROI pairs (mice) | < 0.001 |
| **S10C** | Fezf2-Cre AL-AL change in correlation between DOI and combined baseline and recovery vs zero | Two-tailed hierarchical bootstrap | 81224 (8) | ROI pairs (mice) | < 0.0073 |
|  | Fezf2-Cre PM-PM change in correlation between DOI and combined baseline and recovery vs zero | Two-tailed hierarchical bootstrap | 1177224 (8) | ROI pairs (mice) | < 0.0047 |
|  | Fezf2-Cre AL-PM change in correlation between DOI and combined baseline and recovery vs zero | Two-tailed hierarchical bootstrap | 621192 (8) | ROI pairs (mice) | < 0.001 |
|  | Fezf2-Cre AL-AL change in correlation between DOI and combined baseline and recovery vs PM-PM change in correlation between DOI and combined baseline and recovery | Two-tailed paired hierarchical bootstrap | 81224 (8), 1177224 (8) | ROI pairs (mice) | 0.51 |
|  | Fezf2-Cre PM-PM change in correlation between DOI and combined baseline and recovery vs AL-PM change in correlation between DOI and combined baseline and recovery | Two-tailed paired hierarchical bootstrap | 1177224 (8), 621192 (8) | ROI pairs (mice) | < 0.014 |
|  | Fezf2-Cre AL-AL change in correlation between DOI and combined baseline and recovery vs AL-PM change in correlation between DOI and combined baseline and recovery | Two-tailed paired hierarchical bootstrap | 81224 (8), 621192 (8) | ROI pairs (mice) | 0.050 |
| **S10D** | PV-Cre AL-AL change in correlation between DOI and combined baseline and recovery vs zero | Two-tailed hierarchical bootstrap | 50765 (5) | ROI pairs (mice) | 0.34 |
|  | PV-Cre PM-PM change in correlation between DOI and combined baseline and recovery vs zero | Two-tailed hierarchical bootstrap | 735765 (5) | ROI pairs (mice) | < 0.001 |
|  | PV-Cre AL-PM change in correlation between DOI and combined baseline and recovery vs zero | Two-tailed hierarchical bootstrap | 388245 (5) | ROI pairs (mice) | < 0.001 |
|  | PV-Cre AL-AL change in correlation between DOI and combined baseline and recovery vs PM-PM change in correlation between DOI and combined baseline and recovery | Two-tailed paired hierarchical bootstrap | 50765 (5), 735765 (5) | ROI pairs (mice) | < 0.001 |
|  | PV-Cre PM-PM change in correlation between DOI and combined baseline and recovery vs AL-PM change in correlation between DOI and combined baseline and recovery | Two-tailed paired hierarchical bootstrap | 735765 (5), 388245 (5) | ROI pairs (mice) | 0.55 |
|  | PV-Cre AL-AL change in correlation between DOI and combined baseline and recovery vs AL-PM change in correlation between DOI and combined baseline and recovery | Two-tailed paired hierarchical bootstrap | 50765 (5), 388245 (5) | ROI pairs (mice) | < 0.001 |
| **S10E** | SST-Cre AL-AL change in correlation between DOI and combined baseline and recovery vs zero | Two-tailed hierarchical bootstrap | 91377 (9) | ROI pairs (mice) | < 0.0023 |
|  | SST-Cre PM-PM change in correlation between DOI and combined baseline and recovery vs zero | Two-tailed hierarchical bootstrap | 1324377 (9) | ROI pairs (mice) | < 0.0019 |
|  | SST-Cre AL-PM change in correlation between DOI and combined baseline and recovery vs zero | Two-tailed hierarchical bootstrap | 698841 (9) | ROI pairs (mice) | < 0.001 |
|  | SST-Cre AL-AL change in correlation between DOI and combined baseline and recovery vs PM-PM change in correlation between DOI and combined baseline and recovery | Two-tailed paired hierarchical bootstrap | 91377 (9), 1324377 (9) | ROI pairs (mice) | 0.40 |
|  | SST-Cre PM-PM change in correlation between DOI and combined baseline and recovery vs AL-PM change in correlation between DOI and combined baseline and recovery | Two-tailed paired hierarchical bootstrap | 1324377 (9), 698841 (9) | ROI pairs (mice) | < 0.001 |
|  | SST-Cre AL-AL change in correlation between DOI and combined baseline and recovery vs AL-PM change in correlation between DOI and combined baseline and recovery | Two-tailed paired hierarchical bootstrap | 91377 (9), 698841 (9) | ROI pairs (mice) | < 0.001 |
| **S12A** | Cux2-Cre AL DOI vs baseline power | Two-tailed paired hierarchical bootstrap | 1144 (8), 1144 (8) | ROIs (mice) | In figure for each frequency bin |
|  | Cux2-Cre AL DOI vs recovery power | Two-tailed paired hierarchical bootstrap | 1144 (8), 1144 (8) | ROIs (mice) | In figure for each frequency bin |
|  | Cux2-Cre AL Baseline vs recovery power | Two-tailed paired hierarchical bootstrap | 1144 (8), 1144 (8) | ROIs (mice) | In figure for each frequency bin |
|  | Cux2-Cre RSPd DOI vs baseline power | Two-tailed paired hierarchical bootstrap | 544 (8), 544 (8) | ROIs (mice) | In figure for each frequency bin |
|  | Cux2-Cre RSPd DOI vs recovery power | Two-tailed paired hierarchical bootstrap | 544 (8), 544 (8) | ROIs (mice) | In figure for each frequency bin |
|  | Cux2-Cre RSPd Baseline vs recovery power | Two-tailed paired hierarchical bootstrap | 544 (8), 544 (8) | ROIs (mice) | In figure for each frequency bin |
| **S12B** | Tlx3-Cre AL DOI vs baseline power | Two-tailed paired hierarchical bootstrap | 1573 (11), 1573 (11) | ROIs (mice) | In figure for each frequency bin |
|  | Tlx3-Cre AL DOI vs recovery power | Two-tailed paired hierarchical bootstrap | 1573 (11), 1573 (11) | ROIs (mice) | In figure for each frequency bin |
|  | Tlx3-Cre AL Baseline vs recovery power | Two-tailed paired hierarchical bootstrap | 1573 (11), 1573 (11) | ROIs (mice) | In figure for each frequency bin |
|  | Tlx3-Cre RSPd DOI vs baseline power | Two-tailed paired hierarchical bootstrap | 748 (11), 748 (11) | ROIs (mice) | In figure for each frequency bin |
|  | Tlx3-Cre RSPd DOI vs recovery power | Two-tailed paired hierarchical bootstrap | 748 (11), 748 (11) | ROIs (mice) | In figure for each frequency bin |
|  | Tlx3-Cre RSPd Baseline vs recovery power | Two-tailed paired hierarchical bootstrap | 748 (11), 748 (11) | ROIs (mice) | In figure for each frequency bin |
| **S12C** | Fezf2-Cre AL DOI vs baseline power | Two-tailed paired hierarchical bootstrap | 1144 (8), 1144 (8) | ROIs (mice) | In figure for each frequency bin |
|  | Fezf2-Cre AL DOI vs recovery power | Two-tailed paired hierarchical bootstrap | 1144 (8), 1144 (8) | ROIs (mice) | In figure for each frequency bin |
|  | Fezf2-Cre AL Baseline vs recovery power | Two-tailed paired hierarchical bootstrap | 1144 (8), 1144 (8) | ROIs (mice) | In figure for each frequency bin |
|  | Fezf2-Cre RSPd DOI vs baseline power | Two-tailed paired hierarchical bootstrap | 544 (8), 544 (8) | ROIs (mice) | In figure for each frequency bin |
|  | Fezf2-Cre RSPd DOI vs recovery power | Two-tailed paired hierarchical bootstrap | 544 (8), 544 (8) | ROIs (mice) | In figure for each frequency bin |
|  | Fezf2-Cre RSPd Baseline vs recovery power | Two-tailed paired hierarchical bootstrap | 544 (8), 544 (8) | ROIs (mice) | In figure for each frequency bin |
| **S12D** | PV-Cre AL DOI vs baseline power | Two-tailed paired hierarchical bootstrap | 715 (5), 715 (5) | ROIs (mice) | In figure for each frequency bin |
|  | PV-Cre AL DOI vs recovery power | Two-tailed paired hierarchical bootstrap | 715 (5), 715 (5) | ROIs (mice) | In figure for each frequency bin |
|  | PV-Cre AL Baseline vs recovery power | Two-tailed paired hierarchical bootstrap | 715 (5), 715 (5) | ROIs (mice) | In figure for each frequency bin |
|  | PV-Cre RSPd DOI vs baseline power | Two-tailed paired hierarchical bootstrap | 340 (5), 340 (5) | ROIs (mice) | In figure for each frequency bin |
|  | PV-Cre RSPd DOI vs recovery power | Two-tailed paired hierarchical bootstrap | 340 (5), 340 (5) | ROIs (mice) | In figure for each frequency bin |
|  | PV-Cre RSPd Baseline vs recovery power | Two-tailed paired hierarchical bootstrap | 340 (5), 340 (5) | ROIs (mice) | In figure for each frequency bin |
| **S12E** | SST-Cre AL DOI vs baseline power | Two-tailed paired hierarchical bootstrap | 1287 (9), 1287 (9) | ROIs (mice) | In figure for each frequency bin |
|  | SST-Cre AL DOI vs recovery power | Two-tailed paired hierarchical bootstrap | 1287 (9), 1287 (9) | ROIs (mice) | In figure for each frequency bin |
|  | SST-Cre AL Baseline vs recovery power | Two-tailed paired hierarchical bootstrap | 1287 (9), 1287 (9) | ROIs (mice) | In figure for each frequency bin |
|  | SST-Cre RSPd DOI vs baseline power | Two-tailed paired hierarchical bootstrap | 612 (9), 612 (9) | ROIs (mice) | In figure for each frequency bin |
|  | SST-Cre RSPd DOI vs recovery power | Two-tailed paired hierarchical bootstrap | 612 (9), 612 (9) | ROIs (mice) | In figure for each frequency bin |
|  | SST-Cre RSPd Baseline vs recovery power | Two-tailed paired hierarchical bootstrap | 612 (9), 612 (9) | ROIs (mice) | In figure for each frequency bin |
| **S13A** | Cux2-Cre linear regression of the ratio of AL variance to PM variance against the correlation between the AL and PM domains | F-test | 1, 2 | Degrees of freedom vs Degrees of Freedom for error | < 0.005 |

|  |  |  |  |  |
| --- | --- | --- | --- | --- |
| Tlx3-Cre linear regression of the ratio of AL variance to PM variance against the correlation between the AL and PM domains | F-test | 1, 2 | Degrees of freedom vs Degrees of Freedom for error | < 0.023 |
| Fezf2-Cre linear regression of the ratio of AL variance to PM variance against the correlation between the AL and PM domains | F-test | 1, 2 | Degrees of freedom vs Degrees of Freedom for error | < 0.023 |
| PV-Cre linear regression of the ratio of AL variance to PM variance against the correlation between the AL and PM domains | F-test | 1, 2 | Degrees of freedom vs Degrees of Freedom for error | 0.49 |
| SST-Cre linear regression of the ratio of AL variance to PM variance against the correlation between the AL and PM domains | F-test | 1, 2 | Degrees of freedom vs Degrees of Freedom for error | < 0.001 |

**Table S2.** Analyses performed with each mouse. List of all mice used in this paper along with how they contribute to the different figures. Mice for which all entries in this table are identical are collapsed to a single row.

| Number of mice | Genotype | Experiment | AAV vectors | Delivery method | Figure |
| --- | --- | --- | --- | --- | --- |
| 11 | C57BL6/J | Widefield imaging | PHP.eB-EF1α-GCaMP6s-WPRE | Retroorbital deposit | 1C-H, 2B-F, 4, 5D-E, S1A-B, S2, S4, S5B, S6A,B |
| 5 | C57BL6/J | Widefield imaging<br>ketanserin+DOI | PHP.eB-EF1α-GCaMP6s-WPRE | Retroorbital deposit | 3B-F, 4C-D, 5A-B |
| 4 | C57BL6/J | Widefield EGFP imaging | PHP.eB-EF1α-EGFP-WPRE | Retroorbital deposit | S1C-D, S3, S5, S6C, S7 |
| 2 | Tlx3 x Ai32 wt/wt ki/wt | Widefield EGFP imaging | PHP.eB-EF1α-EGFP-WPRE | Retroorbital deposit | S1C-D, S3, S5, S6C, S7 |
| 5 | C57BL6/J | EEG recordings | PHP.eB-EF1α-GCaMP6s-WPRE | Retroorbital deposit | 5D-E |
| 9 | Tlx3-Cre x Ai32 | Widefield L5 neurons stimulation imaging | PHP.eB-hSyn-PinkyCaMP | Retroorbital deposit | 6C-H |
| 4 | C57BL6/J | Widefield thalamic stimulation imaging | PHP.eB-hSyn-PinkyCaMP<br>PHP.eB-Syn-ChR2(H134R)-GFP | PinkyCaMP: retroorbital deposit, ChR2: injection into left VAL and right LP | 7C-G, S8, S9B |
| 1 | C57BL6/J | Widefield thalamic stimulation imaging | PHP.eB-hSyn-PinkyCaMP<br>PHP.eB-Syn-ChR2(H134R)-GFP<br>PHP.eB-EF1α-EGFP-WPRE | PinkyCaMP: retroorbital deposit, ChR2: injection into left VAL and right LP, EGFP: injection into right VAL and left LP | 7C-G, S9B |
| 6 | C57BL6/J | Widefield thalamic stimulation imaging | PHP.eB-hSyn-PinkyCaMP 2/1-Syn-ChR2(H134R)-GFP | PinkyCaMP: retroorbital deposit, ChR2: injection into left VAL and right AV | 7D-G, S9 |
| 4 | Cux2-CreERT2 | Widefield imaging | PHP.eB-hSyn-FLEX-jGCaMP8s-WPRE | Retroorbital deposit | 8A,B, S10A, S11A, S12A, S13 |
| 4 | Cux2-CreERT2 x Ai148 | Widefield imaging | - | - | 8A,B, S10A, S11A, S12A, S13 |
| 11 | Tlx3-Cre x Ai148 | Widefield imaging | - | - | 8A,B, S10B, S11B, S12B, S13 |
| 8 | Fezf2-CreERT2 x Ai148 | Widefield imaging | - | - | 8A,B, S10C, S11C, S12C, S13 |
| 5 | PV-Cre | Widefield imaging | PHP.eB-hSyn-FLEX- jGCaMP8s-WPRE | Retroorbital deposit | 8A,B, S10D, S11D, S12D, S13 |
| 4 | SST-Cre | Widefield imaging | PHP.eB-hSyn-FLEX-jGCaMP8s-WPRE | Retroorbital deposit | 8A,B, S10E, S11E, S12E, S13 |
| 5 | SST-Cre | Widefield imaging | PHP.eB-EF1α-DIO-GCaMP6s | Retroorbital deposit | 8A,B, S10E, S11E, S12E, S13 |

